# Molecular Basis for G-protein-Coupled Receptor (GPCR) Activation and Biased Signalling at the Platelet Thrombin Receptor Proteinase Activated Receptor-4 (PAR4)

**DOI:** 10.1101/746693

**Authors:** Pierre E. Thibeault, Jordan C. LeSarge, D’Arcy Arends, Michaela Fernandes, Peter Chidiac, Peter B. Stathopulos, Leonard G. Luyt, Rithwik Ramachandran

**Author notes:** For correspondence: Rithwik Ramachandran, Department of Physiology and Pharmacology, Schulich School of Medicine and Dentistry, University of Western Ontario, London, Ontario, Canada.

## Abstract

Proteinase Activated Receptor-4 (PAR4) is a member of the proteolytically-activated PAR family of G-Protein-coupled Receptors (GPCRs). PARs are activated following proteolytic cleavage of the receptor N-terminus by enzymes such as thrombin, trypsin, and cathepsin-G to reveal the receptor-activating motif termed the tethered ligand. The tethered ligand binds intramolecularly to the receptor and triggers receptor signalling and cellular responses. In spite of this unusual mechanism of activation, PARs are fundamentally peptide receptors and can also be activated by exogenous application of short synthetic peptides derived from the tethered ligand sequence. In order to gain a better understanding of the molecular basis for PAR4-dependent signalling, we examined signalling responses to a library of peptides derived from the canonical PAR4 activating peptide (PAR4-AP), AYPGKF-NH_2_. We examined peptide residues involved in activation of the G*α*_q/11_-coupled calcium signalling pathway, β-arrestin recruitment, and mitogen-activated protein kinase pathway activation. The peptide *N*-methyl-alanine-YPGKF-NH_2_ was identified as a compound that is a poor activator of PAR4-dependent calcium signalling but was fully competent in recruiting β-arrestin-1 and -2. In order to gain a better understanding of the ligand-binding pocket, we used *in silico* docking to identify key residues involved in PAR4 interaction with AYPGKF-NH_2_. The predicted interactions were verified by site-directed mutagenesis and analysis of calcium signalling and β-arrestin-1/-2 recruitment following proteolytic activation (with thrombin) or activation with the synthetic agonist peptide (AYPGKF-NH_2_). We determined that a key extracellular loop-2 aspartic acid residue (Asp^230^) is critical for signalling following both proteolytic and peptide activation of PAR4. Finally, we investigated platelet aggregation in response to A**y**PGKF-NH_2_ (a peptide with D-tyrosine in position two) which is unable to activate calcium signalling, and AYPG**R**F-NH_2_ a peptide that is equipotent to the parental peptide AYPGKF-NH_2_ for calcium signalling but is more potent at recruiting β-arrestins. We found that A**y**PGKF-NH_2_ fails to activate platelets while AYPG**R**F-NH_2_ causes a platelet aggregation response that is greater than that seen with the parental peptide and is comparable to that seen with thrombin stimulation. Overall, these studies uncover molecular determinants for agonist binding and signalling through a non-canonically activated GPCR and provide a template for development of small molecule modulators of PAR4.

## Introduction

G-Protein-coupled receptors (GPCRs) are the largest family of cell surface receptors and regulate a host of important physiological responses (Fredriksson and Schioth, 2005; Pierce et al., 2002). GPCRs respond to a variety of extracellular signals and regulate cellular behaviour through engaging intracellular effector molecules to activate various cell-signalling pathways (Erlandson et al., 2018; Hilger et al., 2018). In drug discovery, GPCRs are a valuable and highly tractable class of drug targets with over 30% of all currently approved drugs acting on GPCRs (Sriram and Insel, 2018). GPCR activation occurs typically through binding of a soluble ligand and conformational changes that enable engagement of G-protein-dependent or β-arrestin-mediated signal transduction. The Proteinase Activated Receptors (PARs) are a four-member family of GPCRs with a distinct mechanism of activation that involves limited proteolysis and unmasking of a receptor activating motif called the tethered ligand. PARs were first discovered in an effort to identify the cellular receptors responsible for actions of thrombin that were independent from its role in the coagulation cascade (Vu et al., 1991). Various proteinases have now been described as activators of the different members of the PAR family, but canonically PAR1 and PAR4 are described as thrombin-activated receptors, while PAR2 is activated by trypsin and other trypsin-like serine proteinases. PAR4 can also be activated by trypsin. Whether PAR3 signals independently is unclear, though thrombin can cleave the PAR3 N-terminus to reveal a tethered ligand (Adams et al., 2011; Ishihara et al., 1997). Following the discovery of PARs it was quickly realized that two receptors, PAR1 and PAR4, serve as the thrombin receptors on human platelets (Coughlin, 2005, 1999). PAR1 and PAR4 both trigger platelet activation and aggregation. Given that anti-platelet agents are an important class of drugs for the treatment of various cardiovascular diseases, there has been a concerted effort aimed at understanding the role of these receptors in platelet activation and in identifying molecules that can act as PAR1 and PAR4 antagonists. A PAR1 antagonist was recently identified and approved for clinical use, however significant bleeding side effects were noted in patients administered this drug (Morrow et al., 2012, 2009; Scirica et al., 2012). While bleeding is a frequently encountered side effect of therapeutically targeting pathways involved in blood coagulation, it was hoped that targeting PAR1 would not present such liability. Much attention has now shifted to understanding and targeting the other thrombin receptor PAR4. Though both PAR1 and PAR4 are activated by thrombin, there are some key differences reported in the activation of these two receptors on platelets that argue for PAR4 being a better anti-platelet drug target than PAR1. PAR1 possesses a hirudin-like binding site that enables it to bind thrombin with relatively high affinity (Jacques et al., 2000; Liu et al., 1991). In lacking this thrombin-binding site, PAR4 can only be activated at much higher concentrations of thrombin, such as may be encountered on a growing platelet thrombus. There are also reported differences in the kinetics of PAR1 and PAR4-dependent calcium signalling, with PAR4 causing a delayed but more sustained signal compared to PAR1 (Shapiro et al., 2000). This is thought to be due to dual proline residues in the PAR4 exodomain that provide low-affinity interactions with the thrombin active site and an anionic cluster that slows dissociation from thrombin (Jacques and Kuliopulos, 2003). Overall, PAR1 is thought to initiate platelet activation in response to low concentrations of thrombin while PAR4 is engaged at higher concentrations to further consolidate and stabilize the clot.

Given the liabilities seen with targeting PAR1 as a platelet thrombin receptor there has been renewed interest in developing small molecule PAR4 antagonists (Rwibasira Rudinga et al., 2018; Wu et al., 2000; Young et al., 2010). Recently small molecule antagonists of PAR4 have been described with efficacy in inhibiting injury induced thrombosis in both non-human primates and humans (Miller et al., 2019; Wilson et al., 2017; Wong et al., 2017). Despite these promising advances there remains the question of whether a complete inhibition of PAR4 might also lead to bleeding liability. Our efforts have therefore focused on examining biased signalling through PAR4 in order to identify pathways that are responsible for platelet aggregation and that could be selectively manipulated for therapeutic efficacy. PAR4 can activate multiple signalling pathways through coupling to G*α*_q/11_, G*α*_12/13_ and β-arrestin (Holinstat et al., 2006; Li et al., 2011; Ramachandran et al., 2017). In previous work, we and others showed that pepducins targeting different intracellular motifs in PAR4 could be employed to differentially inhibit platelet activation by blocking G-protein-coupled or β-arrestin-dependent signalling (Covic et al., 2002; Ramachandran et al., 2017). The ability to selectively activate or inhibit GPCR signalling in a biased manner is of great interest and has been proposed as a strategy for obtaining therapeutically superior drugs for a number of conditions (Bar-Shavit et al., 2016; Hollenberg et al., 2014; Seyedabadi et al., 2019). In the case of the finely balanced signalling systems involved in coagulation and platelet activation, such subtle perturbations may be key to obtaining drugs which do not result in bleeding liability.

Here we seek to gain an understanding of the molecular interactions responsible for agonist docking to PAR4 and to identify interactions that can be manipulated to obtain biased ligands for PAR4. By screening a peptide library derived from the most widely used PAR4 agonist peptide, AYPGKF-NH_2_, we have identified key residues that confer selective activation of G*α*_q/11_ coupled calcium signalling and/or β-arrestin recruitment. Through *in silico* docking and confirmatory mutagenesis experiments we further identify an extracellular loop 2 residue (Asp^230^) that is critical for receptor activation and signalling. We also examine platelet activation by peptides that show differential activation of PAR4 signalling. Overall, these studies advance our understanding of PAR4 activation and signalling and will help guide future drug discovery efforts targeting this receptor.

## Results

### Experimental design and data analysis

In order to understand the rules governing agonist peptide activation of PAR4 we synthesized 37 hexapeptides through modification to the synthetic PAR4 agonist peptide AYPGKF-NH_2_. A number of different modifications were examined at each of the six positions, including alanine, D-isomer, and *N-*methyl substitutions, as well as other modifications aimed at further probing each of the six agonist peptide positions. Henceforth, the parental PAR4 agonist peptide for this study, AYPGKF-NH_2_ (Ala^1^-Tyr^2^-Pro^3^-Gly^4^-Lys^5^-Phe^6^-NH_2_; superscript denotes position from N-terminus), will be referred to as PAR4-AP. G*α*_q/11_-coupled calcium signalling, β-arrestin-1/-2 recruitment (β-Arr-1-rluc, β-Arrestin-2-rluc respectively), and p44/42 mitogen-activated protein kinase (MAPK) signalling by each of these peptides was monitored and compared to responses triggered by PAR4-AP (Table 1). Whereas PAR4-AP triggered G*α*_q/11_ signalling reached a clear plateau over the range of concentrations tested (0.3 – 300 µM), β-arrestin recruitment did not and therefore the effective concentration required to achieve half-maximal response (EC_50_) values could not be accurately assessed. In contrast, with the substituted peptides, G*α*_q/11_ signalling concentration-effect curves in some cases did not clearly plateau, whereas β-arrestin recruitment did. Where possible, EC_50_ values are compared between PAR4-AP and substituted peptides. However, since clear maxima were not always obtained, comparisons of signal magnitude throughout the present study are presented as the response elicited by a given compound at 100 µM, shown as a percentage of responses elicited by PAR4-AP at the same concentration. Since the concentration-effect curve for PAR4-AP exhibits a saturating calcium signal, peptides that failed to saturate calcium signalling responses are depicted as having EC_50_ values that were ‘not determined’ (*n.d.*). Maximal calcium signal responses are shown as a percentage of the response obtained with the calcium ionophore A23187 (Table 1, Max; % of A23187 calcium ionophore) and are referred to throughout the results as a change in maximal calcium signalling as a measure of the efficacy of the peptides. Throughout the results, β-arrestin recruitment values are presented as a percentage (%) of the PAR4-AP-induced recruitment at 100 µM concentration. For ease of comparison, concentration-effect curves for PAR4-AP-mediated G*α*_q/11_ calcium signalling and β-arrestin recruitment are displayed as a repeated reference value in each of the Figures 1 through 7.

**Figure 1.**
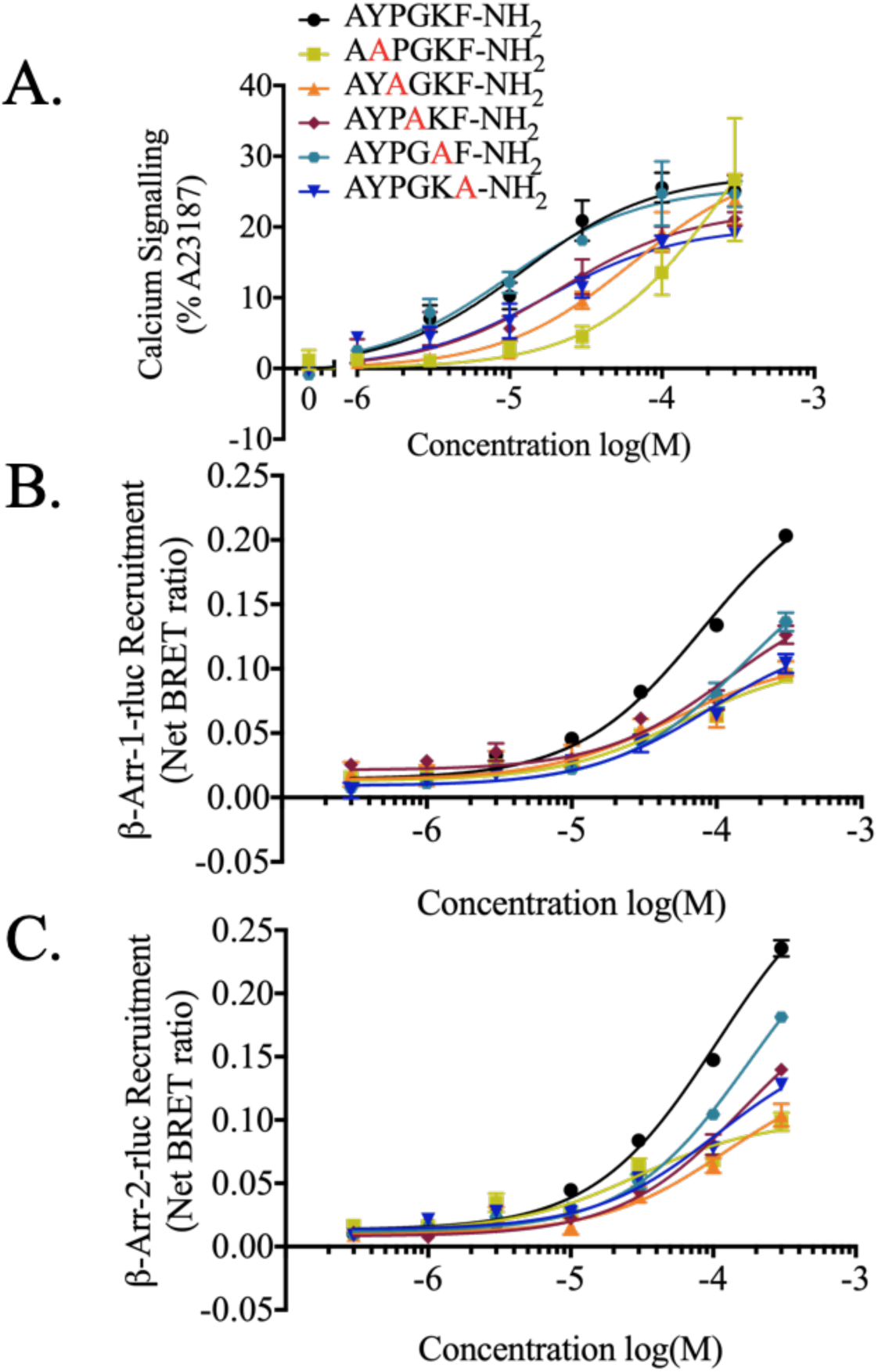
Calcium and β-arrestin recruitment in response to alanine substitutions in the PAR4-AP. **(A) Agonist-stimulated calcium signalling.** Calcium signalling is shown as a percentage of the maximum calcium signalling obtained by application of calcium ionophore (A23187, 3 µM). (*n = 3-7*) Agonist-stimulated recruitment of β-arrestin-1 **(B) and β-arrestin-2 (C) to PAR4.** Concentration effect curves of BRET values from cells treated increasing concentrations of peptide are presented as normalized net BRET ratio. (*n = 3*).

**Table 1.**
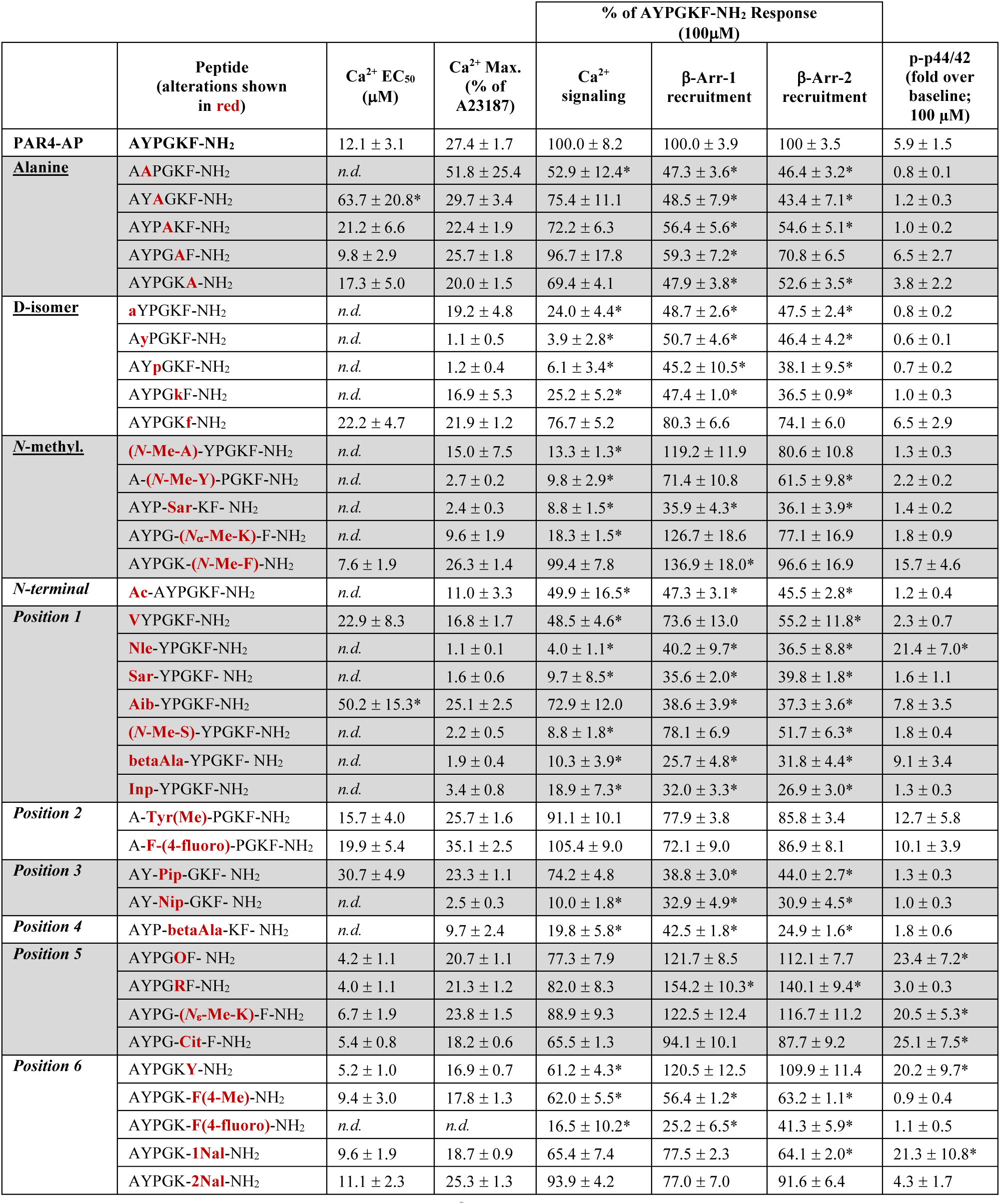
Summary table of all calcium, β-arrestin, and p44/42 data. Calcium data is displayed as EC_50_ or not determined (*n.d.*) if EC_50_ could not be calculated. Maximum calcium response achieved with each peptide is shown as a percentage of the maximum calcium response achievable in a given experiment revealed by calcium ionophore (% of A23187 calcium ionophore). Statistically significant shifts in EC_50_ compared to PAR4-AP, AYPGKF-NH_2_, were found using the F-statistic (*p < 0.05). Given that saturation did not always occur, all peptides are compared to the parental PAR4-AP, AYPGKF-NH_2_, and shown as a percent of the response obtained with PAR4-AP at a given concentration (100 µM). Finally, fold increase over unstimulated baseline is shown for phosphorylation of p44/42 MAP kinase activation. Statistical significance for comparisons at 100 µM and p44/42 phosphorylation are calculated using one-way ANOVA of response compared to PAR4-AP (*p < 0.05).

|  | Peptide<br>(alterations shown<br>in red) | Ca<sup>2+</sup> EC<sub>50</sub><br>(μM) | Ca<sup>2+</sup> Max.<br>(% of<br>A23187) | % of AYPGKF-NH<sub>2</sub> Response<br>(100μM) |  |  | p-p44/42<br>(fold over<br>baseline;<br>100 μM) |
| --- | --- | --- | --- | --- | --- | --- | --- |
|  |  |  |  | Ca<sup>2+</sup><br>signaling | β-Arr-1<br>recruitment | β-Arr-2<br>recruitment |  |
| **PAR4-AP** | **AYPGKF-NH<sub>2</sub>** | 12.1 ± 3.1 | 27.4 ± 1.7 | 100.0 ± 8.2 | 100.0 ± 3.9 | 100 ± 3.5 | 5.9 ± 1.5 |
| **Alanine** | **A**APGKF-NH<sub>2</sub> | *n.d.* | 51.8 ± 25.4 | 52.9 ± 12.4* | 47.3 ± 3.6* | 46.4 ± 3.2* | 0.8 ± 0.1 |
|  | AY**A**GKF-NH<sub>2</sub> | 63.7 ± 20.8* | 29.7 ± 3.4 | 75.4 ± 11.1 | 48.5 ± 7.9* | 43.4 ± 7.1* | 1.2 ± 0.3 |
|  | AYP**A**KF-NH<sub>2</sub> | 21.2 ± 6.6 | 22.4 ± 1.9 | 72.2 ± 6.3 | 56.4 ± 5.6* | 54.6 ± 5.1* | 1.0 ± 0.2 |
|  | AYPG**A**F-NH<sub>2</sub> | 9.8 ± 2.9 | 25.7 ± 1.8 | 96.7 ± 17.8 | 59.3 ± 7.2* | 70.8 ± 6.5 | 6.5 ± 2.7 |
|  | AYPGK**A**-NH<sub>2</sub> | 17.3 ± 5.0 | 20.0 ± 1.5 | 69.4 ± 4.1 | 47.9 ± 3.8* | 52.6 ± 3.5* | 3.8 ± 2.2 |
| **D-isomer** | **a**YPGKF-NH<sub>2</sub> | *n.d.* | 19.2 ± 4.8 | 24.0 ± 4.4* | 48.7 ± 2.6* | 47.5 ± 2.4* | 0.8 ± 0.2 |
|  | A**y**PGKF-NH<sub>2</sub> | *n.d.* | 1.1 ± 0.5 | 3.9 ± 2.8* | 50.7 ± 4.6* | 46.4 ± 4.2* | 0.6 ± 0.1 |
|  | AY**p**GKF-NH<sub>2</sub> | *n.d.* | 1.2 ± 0.4 | 6.1 ± 3.4* | 45.2 ± 10.5* | 38.1 ± 9.5* | 0.7 ± 0.2 |
|  | AYPG**k**F-NH<sub>2</sub> | *n.d.* | 16.9 ± 5.3 | 25.2 ± 5.2* | 47.4 ± 1.0* | 36.5 ± 0.9* | 1.0 ± 0.3 |
|  | AYPGK**f**-NH<sub>2</sub> | 22.2 ± 4.7 | 21.9 ± 1.2 | 76.7 ± 5.2 | 80.3 ± 6.6 | 74.1 ± 6.0 | 6.5 ± 2.9 |
| **N-methyl.** | **(N-Me-A)**-YPGKF-NH<sub>2</sub> | *n.d.* | 15.0 ± 7.5 | 13.3 ± 1.3* | 119.2 ± 11.9 | 80.6 ± 10.8 | 1.3 ± 0.3 |
|  | A-**(N-Me-Y)**-PGKF-NH<sub>2</sub> | *n.d.* | 2.7 ± 0.2 | 9.8 ± 2.9* | 71.4 ± 10.8 | 61.5 ± 9.8* | 2.2 ± 0.2 |
|  | AYP-**Sar**-KF- NH<sub>2</sub> | *n.d.* | 2.4 ± 0.3 | 8.8 ± 1.5* | 35.9 ± 4.3* | 36.1 ± 3.9* | 1.4 ± 0.2 |
|  | AYPG-**(N<sub>α</sub>-Me-K)**-F-NH<sub>2</sub> | *n.d.* | 9.6 ± 1.9 | 18.3 ± 1.5* | 126.7 ± 18.6 | 77.1 ± 16.9 | 1.8 ± 0.9 |
|  | AYPGK-**(N-Me-F)**-NH<sub>2</sub> | 7.6 ± 1.9 | 26.3 ± 1.4 | 99.4 ± 7.8 | 136.9 ± 18.0* | 96.6 ± 16.9 | 15.7 ± 4.6 |
| **N-terminal** | **Ac**-AYPGKF-NH<sub>2</sub> | *n.d.* | 11.0 ± 3.3 | 49.9 ± 16.5* | 47.3 ± 3.1* | 45.5 ± 2.8* | 1.2 ± 0.4 |
| **Position 1** | **V**YPGKF-NH<sub>2</sub> | 22.9 ± 8.3 | 16.8 ± 1.7 | 48.5 ± 4.6* | 73.6 ± 13.0 | 55.2 ± 11.8* | 2.3 ± 0.7 |
|  | **Nle**-YPGKF-NH<sub>2</sub> | *n.d.* | 1.1 ± 0.1 | 4.0 ± 1.1* | 40.2 ± 9.7* | 36.5 ± 8.8* | 21.4 ± 7.0* |
|  | **Sar**-YPGKF- NH<sub>2</sub> | *n.d.* | 1.6 ± 0.6 | 9.7 ± 8.5* | 35.6 ± 2.0* | 39.8 ± 1.8* | 1.6 ± 1.1 |
|  | **Aib**-YPGKF-NH<sub>2</sub> | 50.2 ± 15.3* | 25.1 ± 2.5 | 72.9 ± 12.0 | 38.6 ± 3.9* | 37.3 ± 3.6* | 7.8 ± 3.5 |
|  | **(N-Me-S)**-YPGKF-NH<sub>2</sub> | *n.d.* | 2.2 ± 0.5 | 8.8 ± 1.8* | 78.1 ± 6.9 | 51.7 ± 6.3* | 1.8 ± 0.4 |
|  | **betaAla**-YPGKF- NH<sub>2</sub> | *n.d.* | 1.9 ± 0.4 | 10.3 ± 3.9* | 25.7 ± 4.8* | 31.8 ± 4.4* | 9.1 ± 3.4 |
|  | **Inp**-YPGKF-NH<sub>2</sub> | *n.d.* | 3.4 ± 0.8 | 18.9 ± 7.3* | 32.0 ± 3.3* | 26.9 ± 3.0* | 1.3 ± 0.3 |
| **Position 2** | A-**Tyr(Me)**-PGKF-NH<sub>2</sub> | 15.7 ± 4.0 | 25.7 ± 1.6 | 91.1 ± 10.1 | 77.9 ± 3.8 | 85.8 ± 3.4 | 12.7 ± 5.8 |
|  | A-**F-(4-fluoro)**-PGKF-NH<sub>2</sub> | 19.9 ± 5.4 | 35.1 ± 2.5 | 105.4 ± 9.0 | 72.1 ± 9.0 | 86.9 ± 8.1 | 10.1 ± 3.9 |
| **Position 3** | AY-**Pip**-GKF- NH<sub>2</sub> | 30.7 ± 4.9 | 23.3 ± 1.1 | 74.2 ± 4.8 | 38.8 ± 3.0* | 44.0 ± 2.7* | 1.3 ± 0.3 |
|  | AY-**Nip**-GKF- NH<sub>2</sub> | *n.d.* | 2.5 ± 0.3 | 10.0 ± 1.8* | 32.9 ± 4.9* | 30.9 ± 4.5* | 1.0 ± 0.3 |
| **Position 4** | AYP-**betaAla**-KF- NH<sub>2</sub> | *n.d.* | 9.7 ± 2.4 | 19.8 ± 5.8* | 42.5 ± 1.8* | 24.9 ± 1.6* | 1.8 ± 0.6 |
| **Position 5** | AYPG**O**F- NH<sub>2</sub> | 4.2 ± 1.1 | 20.7 ± 1.1 | 77.3 ± 7.9 | 121.7 ± 8.5 | 112.1 ± 7.7 | 23.4 ± 7.2* |
|  | AYPG**R**F-NH<sub>2</sub> | 4.0 ± 1.1 | 21.3 ± 1.2 | 82.0 ± 8.3 | 154.2 ± 10.3* | 140.1 ± 9.4* | 3.0 ± 0.3 |
|  | AYPG-**(N<sub>ε</sub>-Me-K)**-F-NH<sub>2</sub> | 6.7 ± 1.9 | 23.8 ± 1.5 | 88.9 ± 9.3 | 122.5 ± 12.4 | 116.7 ± 11.2 | 20.5 ± 5.3* |
|  | AYPG-**Cit**-F-NH<sub>2</sub> | 5.4 ± 0.8 | 18.2 ± 0.6 | 65.5 ± 1.3 | 94.1 ± 10.1 | 87.7 ± 9.2 | 25.1 ± 7.5* |
| **Position 6** | AYPGK**Y**-NH<sub>2</sub> | 5.2 ± 1.0 | 16.9 ± 0.7 | 61.2 ± 4.3* | 120.5 ± 12.5 | 109.9 ± 11.4 | 20.2 ± 9.7* |
|  | AYPGK-**F(4-Me)**-NH<sub>2</sub> | 9.4 ± 3.0 | 17.8 ± 1.3 | 62.0 ± 5.5* | 56.4 ± 1.2* | 63.2 ± 1.1* | 0.9 ± 0.4 |
|  | AYPGK-**F(4-fluoro)**-NH<sub>2</sub> | *n.d.* | *n.d.* | 16.5 ± 10.2* | 25.2 ± 6.5* | 41.3 ± 5.9* | 1.1 ± 0.5 |
|  | AYPGK-**1NaI**-NH<sub>2</sub> | 9.6 ± 1.9 | 18.7 ± 0.9 | 65.4 ± 7.4 | 77.5 ± 2.3 | 64.1 ± 2.0* | 21.3 ± 10.8* |
|  | AYPGK-**2NaI**-NH<sub>2</sub> | 11.1 ± 2.3 | 25.3 ± 1.3 | 93.9 ± 4.2 | 77.0 ± 7.0 | 91.6 ± 6.4 | 4.3 ± 1.7 |

### Effect of alanine substitution on PAR4-AP-mediated signalling responses

Our first set of changes involved alanine substitutions at positions 2-6 of the PAR4-AP (calcium EC_50_ = 12.1 ± 3.1 µM; β-arrestin-1 recruitment at 100 µM for comparison is considered 100 ± 3.9%, and 100 ± 3.5% for β-arrestin-2) (Fig. 1, Table 1). Potency was decreased with respect to calcium signalling when tyrosine^2^ was substituted with alanine and failed to reach a plateau over the concentration range tested (A**A**PGKF-NH_2_; EC_50_ = *n.d.*). Correspondingly, β-arrestin recruitment was reduced compared to PAR4-AP at the same concentration (β-arr-1 47.3 ± 3.6%; β-arr-2 46.4 ± 3.2%). Substitution of proline^3^ with alanine resulted in a significant decrease in potency for calcium signalling (AY**A**GKF-NH_2_ EC_50_ = 63.7 ± 20.8 µM) and yielded reduced β-arrestin recruitment (β-arr-1 48.5 ± 7.9%; β-arr-2 43.4 ± 7.1%). Either glycine^4^ or phenylalanine^6^ substitution to alanine resulted in modest decreases in maximal calcium signalling compared to PAR4-AP at the same concentration, with no significant effect on potency (EC_50_ = 21.2 ± 6.6 µM; EC_50_ = 17.3 ± 5.0 µM, respectively). Additionally, alanine substitutions in these positions resulted in significantly reduced β-arrestin recruitment, to levels approximately half of those observed with PAR4-AP (AYP**A**KF-NH_2_ β-arr-1 56.4 ± 5.6%, β-arr-2 54.6 ± 5.1%; AYPGK**A**-NH_2_ β-arr-1 47.9 ± 3.8%, β-arr-2 52.6 ± 3.5%). Interestingly, we observed that lysine^5^ to alanine substitution had no appreciable impact on potency with respect to calcium signalling (AYPG**A**F-NH_2_ EC_50_ = 9.8 ± 2.9 µM); and modestly reduced recruitment of β-arrestin-1 (59.3 ± 7.2%) and -2 (70.8 ± 6.5%). Collectively, alanine substitutions of positions 2-6 resulted in decreased β-arrestin recruitment compared to the parental peptide (Fig. 1 B. and C.; Table 1). Interestingly, we observe differential effects on calcium signalling with significant detriment to potency attributed to tyrosine substitution (Fig. 1 A.). Additionally, substitution of glycine and phenylalanine moderately decreased maximal calcium signalling responses, perhaps indicating that these residues are important for full agonism of this pathway comparative to that achieved with the parental peptide. Notably, all single alanine substitutions of positions 2-6 of the PAR4-AP retained some agonist activity indicating that signalling is not wholly reliant on any single residue alone.

### Effect of D-isomer amino acid substitutions on PAR4-AP-mediated signalling responses

To assess the impact of stereochemical inversion at each residue we generated five peptides with single amino acid substitutions to their respective D-isomers (Fig. 2, Table 1). Glycine possesses only one isomeric form and thus is not substituted. Both L-alanine to D-alanine (**a**YPGKF-NH_2_) and L-lysine to D-lysine (AYPG**k**F-NH_2_) substitutions resulted in significant losses of potency in calcium signalling assays compared to PAR4-AP. Consistent with calcium data, we observed a significant decrease in β-arrestin recruitment compared to the parental PAR4-AP (**a**YPGKF-NH_2_ β-arr-1 48.7 ± 2.6%, β-arr-2 47.5 ± 2.4%; AYPG**k**F-NH_2_ β-arr-1 47.4 ± 1.0%, β-arr-2 36.5 ± 0.9%). Interestingly, both D-tyrosine (A**y**PGKF-NH_2_) and D-proline (AY**p**GKF-NH_2_) substitutions completely abolished calcium signalling. Comparably, β-arrestin recruitment was significantly reduced with stimulation by either peptide (A**y**PGKF-NH_2_ β-arr-1 50.7 ± 4.6%, β-arr-2 46.4 ± 4.2%; AY**p**GKF-NH_2_ β-arr-1 45.2 ± 10.5%, β-arr-2 38.1 ± 9.5%). In keeping with what we observed with phenylalanine to alanine substitution (Fig. 1), D-phenylalanine substitution in position 6 (AYPGK**f**-NH_2_) resulted in less efficacious agonism of both calcium (EC_50_ = 22.2 ± 4.7 µM) and β-arrestin recruitment (β-arr-1 80.3 ± 6.6%, β-arr-2 74.1 ± 6.0%). Overall, D-isomers substitutions of amino acids alanine^1^, tyrosine^2^, proline^3^, and lysine^5^ all result in significant reduction of (D-alanine, D-lysine) or abolition of (D-tyrosine, D-proline) calcium signalling while causing significant reductions in β-arrestin recruitment. Substitution of L-phenylalanine^6^ for D-phenylalanine was the only D-isomer substitution that is somewhat tolerated, resulting in modest reductions in potency with respect to both calcium signalling and β-arrestin recruitment. These data reveal that the sidechain stereochemistry of residues within the AYPGKF-NH_2_ peptide perform a role in agonism.

**Figure 2.**
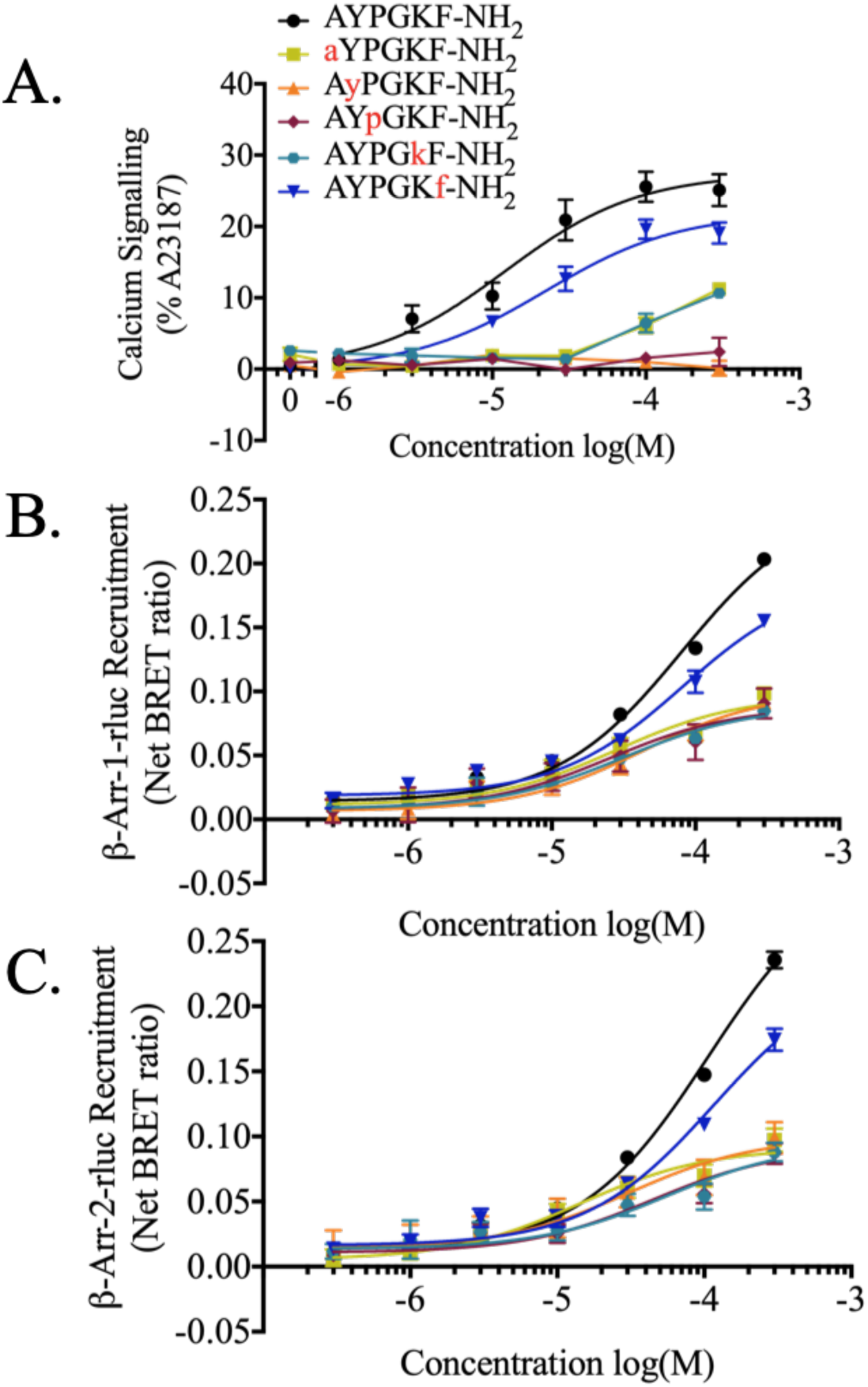
Calcium and β-arrestin recruitment in response to D-isomer substitutions in the PAR4-AP. **(A) Agonist-stimulated calcium signalling.** Calcium signalling is shown as a percentage of the maximum calcium signalling obtained by application of calcium ionophore (A23187, 3 µM). (*n = 3-4*) **Agonist-stimulated recruitment of β-arrestin-1 (B) and β-arrestin-2 (C) to PAR4.** Concentration effect curves of BRET values from cells treated increasing concentrations of peptide are presented as normalized net BRET ratio. (*n = 3*).

### Effect of *N*-methylation on PAR4-AP-mediated signalling responses

We generated a series of peptides to investigate the pharmacological impact of *N*-methylation along the peptide backbone (Fig. 3, Table 1). In contrast to the parental PAR4-AP, all but one of the N-methylated peptides evaluated yielded clear plateaus and discernible EC_50_ values in β-arrestin recruitment assays, and where applicable these are indicated below along with their relative responses compared to PAR4-AP at 100 µM. Substitution of alanine^1^ with *N*-methyl-alanine [**(*N*-Me-A)**-YPGKF-NH_2_] almost completely abolished calcium signalling (EC_50_ = *n.d.*). Interestingly, this peptide was able to initiate β-arrestin-1/-2 recruitment comparable to levels obtained with PAR4-AP (β-arr-1 119.2 ± 11.9%, EC_50_ = 6.8 ± 2.6 µM; β-arr-2 80.6 ± 10.8%; EC_50_ = 2.0 ± 0.7 µM). These results suggest that increasing steric bulk and hydrophobicity of the backbone at amide positions through the addition of an N-terminal methyl group drives PAR4 signalling towards β-arrestin pathways and causes only modest activation of calcium signalling. Substitution of tyrosine^2^ for *N*-methyl-tyrosine [A-**(*N*-Me-Y)**-PGKF-NH_2_] abolished calcium signalling up to 300 µM (EC_50_ = *n.d.*). Consistent with calcium signalling, we observed decreased signal magnitude in β-arrestin recruitment assays compared to PAR4-AP (β-arr-1 71.4 ± 10.8%, EC_50_ = 14.2 ± 8.6 µM; β-arr-2 61.5 ± 9.8%, EC_50_ = 11.8 ± 5.7 µM). Substitution of the PAR4-AP glycine^4^ with sarcosine (Sar., a.k.a *N*-methyl-glycine; AYP-**Sar**-KF-NH_2_) resulted in significantly reduced calcium potency (EC_50_ = *n.d.*) and significantly reduced β-arrestin-1/-2 recruitment (35.9 ± 4.3%, 36.1 ± 3.9% respectively). *N*-methylation of the alpha (α) amine of lysine^5^[AYPG-**(*N*α-Me-K)**-F-NH_2_] resulted in significantly reduced calcium signalling (EC_50_ = *n.d.*) with no significant effect on maximal β-arrestin recruitment (β-arr-1 126.7 ± 18.6%, EC_50_ = 12.2 ± 4.3 µM; β-arr-2 77.1 ± 16.9%, EC_50_ = 2.5 ± 1.0 µM). Substitution of phenylalanine^6^ with *N*-methyl-phenylalanine [AYPGK-**(*N*-Me-F)**-NH_2_] was equipotent for calcium signalling compared to PAR4-AP (EC_50_ = 7.6 ± 1.9 µM). Interestingly, this peptide showed increased recruitment of both β-arrestins (β-arr-1 136.9 ± 18.0%, EC_50_ = 10.4 ± 3.3 µM; β-arr-2 96.6 ± 16.4%, EC_50_ = 15.3 ± 4.6 µM). Together, these data reveal that *N*-methylation of positions 1, 2, 4, and 5 are deleterious to calcium signalling.

**Figure 3.**
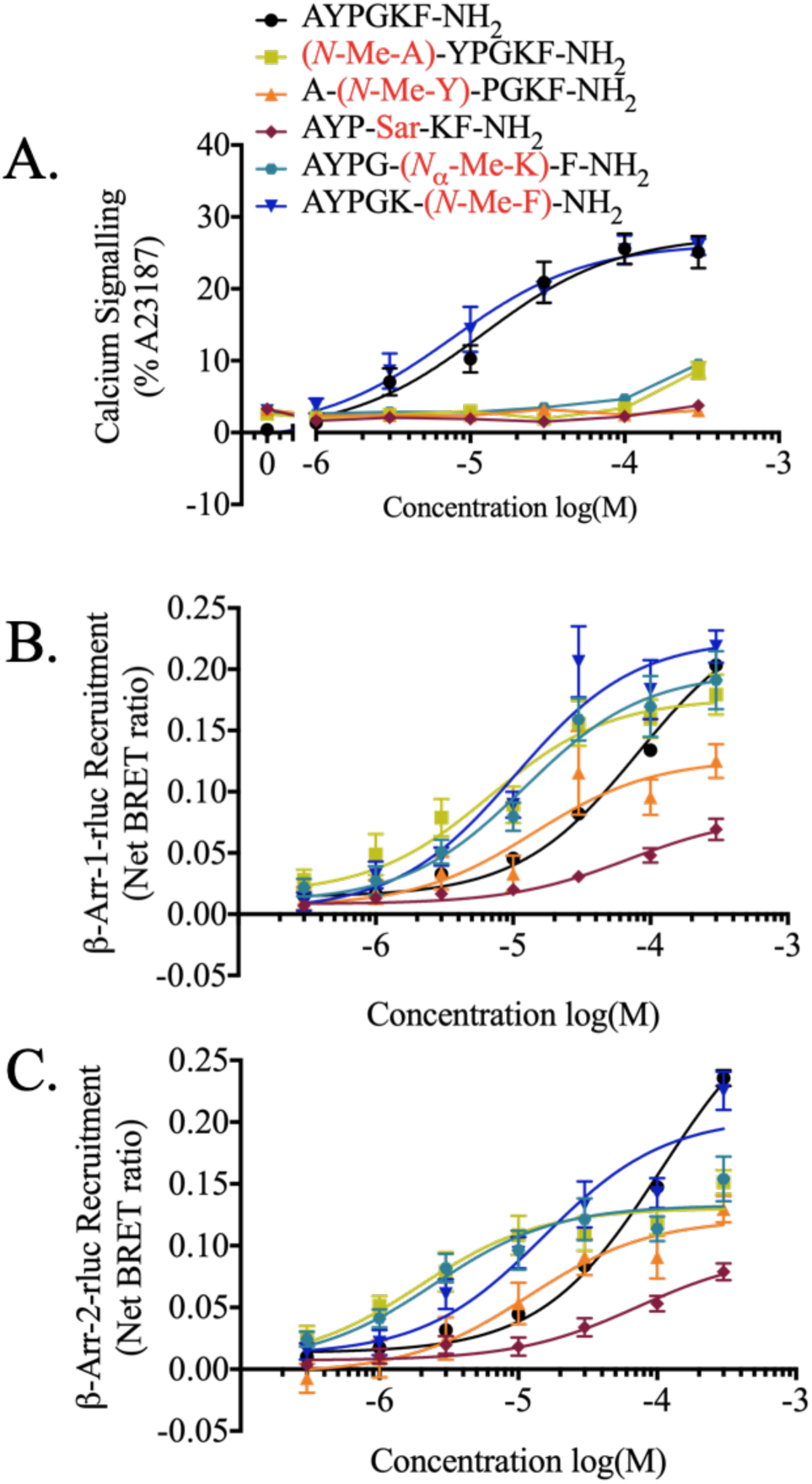
Calcium and β-arrestin recruitment in response to *N*-methylation substitutions in the PAR4-AP. **(A) Agonist-stimulated calcium signalling.** Calcium signalling is shown as a percentage of the maximum calcium signalling obtained by application of calcium ionophore (A23187, 3 µM). (*n = 3-7)* **Agonist-stimulated recruitment of β-arrestin-1 (B) and β-arrestin-2 (C) to PAR4.** Concentration effect curves of BRET values from cells treated increasing concentrations of peptide are presented as normalized net BRET ratio. (*n = 3*).

We were excited to observe evidence of β-arrestin-biased peptides within this group, all of which showed decreased calcium signalling and/or increased β-arrestin recruitment. Moreover, the leftward shifts in their associated β-arrestin concentration-effect profiles imply that substitution of *N*-methylated alanine^1^, tyrosine^2^, lysine^5^, or phenylalanine^6^ residues increases peptide potency with respect to G-protein-independent signalling. Only *N*-methylation of glycine^4^ (sarcosine) resulted in a reduced overall ability to activate PAR4, with relative calcium and β-arrestin signals being reduced by approximately 90% and 60%, respectively.

### Effect of N-terminal and position 1 modifications on PAR4-AP-mediated signalling responses

To further investigate the impact of *N*-terminal alterations to the PAR4-AP we examined the impact of N-terminal acetylation and substitutions of position 1 (Fig. 4, Table 1). Previously it was demonstrated that protonation of the N-terminus of the PAR1 tethered ligand (SFLLRN) is critical for agonist function (Scarborough et al., 1992). Additionally, substitution of glycine for mercaptoproprionic acid (Mpr), which lacks an amino-terminal protonated amine found in the native tethered-ligand-mimicking peptide GYPGQV, resulted in complete loss of agonist activity (Faruqi et al., 2000). Given the evidence for N-terminal protonation in PAR agonism, we sought to determine the impact of N-terminal acetylation on PAR4-AP activity. N-terminal acetylation of the PAR4-AP (**Ac**-AYPGKF-NH2) negatively impacted both calcium signalling (EC_50_ = *n.d.*) as well as the magnitude of β-arrestin recruitment to PAR4 as expected (β-arr-1 47.3 ± 3.1%, EC_50_ = 15.0 ± 4.3 µM; β-arr-2 45.5 ± 2.8%, EC_50_ = 26.7 ± 5.6 µM). In contrast to the previously reported complete loss of activity with Mpr-YPGQV peptide (Mpr = mercaptopropionic acid), we observed partial agonism with the loss of protonation through acetylation of PAR4-AP. This could also be influenced by the additional methyl group of the Ala in Ac-AYPGKF-NH2, which is not present in the Mpr-YPGQV peptide.

**Figure 4.**
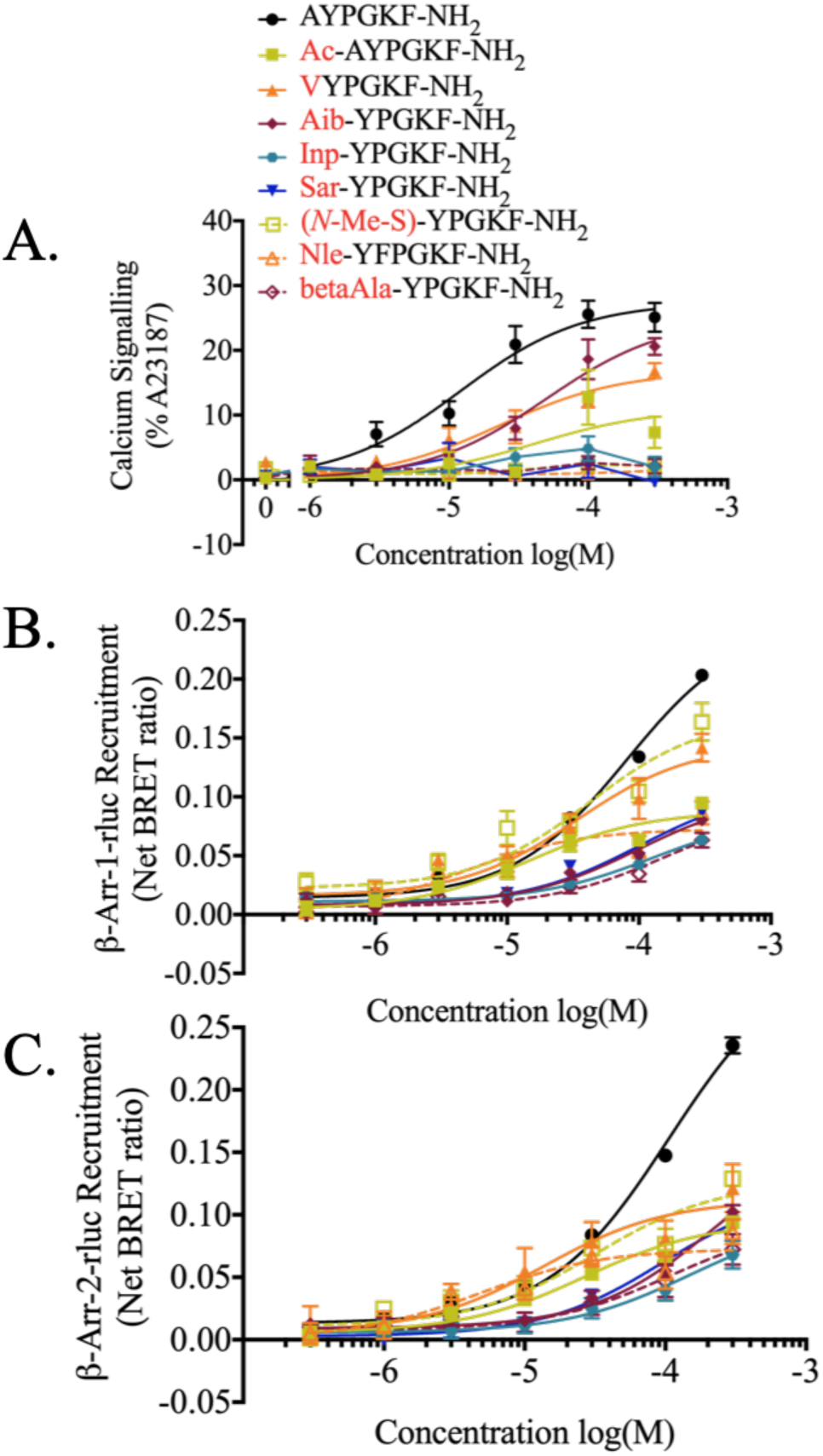
Calcium signalling and β-arrestin recruitment to PAR4 in response to AYPGKF-NH_2_ or peptides with N-terminal acetylation or additional substitutions in position 1 (alanine). **(A) Agonist-stimulated calcium signalling.** Calcium signalling is shown as a percentage of the maximum calcium signalling obtained by application of calcium ionophore (A23187, 3 µM). (*n = 3-7)* **Agonist-stimulated recruitment of β-arrestin-1 (B) and β-arrestin-2 (C) to PAR4.** Concentration effect curves of BRET values from cells treated increasing concentrations of peptide are presented as normalized net BRET ratio. (*n = 3*).

In both the native human PAR4 tethered-ligand sequence and the PAR4-AP, position 1 is occupied by small (tethered ligand – glycine, PAR4-AP – alanine) and non-polar (PAR4-AP – alanine) residues. To investigate the role of position 1 in the PAR4-AP, we generated seven peptides with position 1 substitutions. Introduction of steric bulk and increased hydrophobicity through substitution of alanine^1^ for valine resulted in partial agonism of both calcium signalling and β-arrestin recruitment (**V**YPGKF-NH_2_ EC_50_ = 22.9 ± 8.3 µM; β-arr-1 73.6 ± 13.0%, EC_50_ = 33.8 ± 13.7 µM; β-arr-2 55.2 ± 11.8%, EC_50_ = 13.7 ± 8.2 µM). Additionally, increasing hydrophobic bulk at position 1 through substitution with norleucine completely abolished calcium signalling (**Nle**-YPGKF-NH_2_, EC_50_ = *n.d.*) and resulted in partial agonism of β-arrestin recruitment (β-arr-1 40.2 ± 9.7%; β-arr-2 36.5 ± 8.8%). Together, data from these substitutions indicate that the precise steric bulk of the side chain in position 1 is important for PAR4 agonism.

Next, we evaluated the impact of steric bulk relocation in position 1. Relocation of the methyl group in alanine to the N-terminus and loss of side chain through substitution with sarcosine (**Sar**-YPGKF-NH_2_) resulted in poor agonism at both calcium signalling (EC_50_ = *n.d.*) and β-arrestin recruitment (β-arr-1 35.6 ± 2.0%, β-arr-2 39.8 ± 1.8%). Further, the addition of an α-methyl group through substitution of alanine^1^ with 2-aminoisobutyric acid (**Aib**-YPGKF-NH_2_) resulted in a significantly less potent partial agonist for calcium signalling (EC_50_ = 50.2 ± 15.3 µM) and also resulted in a significant reduction in β-arrestin recruitment (β-arr-1 38.6 ± 3.9%; β-arr-2 37.3 ± 3.6%). Similarly, replacement with *N*-methyl-serine [**(N-Me-S)**-YPGKF-NH_2_] also caused a significant attenuation of calcium signalling (EC_50_ = *n.d.*), while also revealing a decrease in β-arrestin recruitment compared to PAR4-AP (β-arr-1 78.1 ± 6.9%, β-arr-2 51.7 ± 6.3%). Thus, it is apparent that the precise position of the side chain in position 1 is important for agonism.

To further probe the importance of charge localization at the N-terminal region of the peptide, we made several additional substitutions to position 1. Substitution of alanine with either beta-alanine or isonipecotic acid significantly reduced the potency and activity of the PAR4-AP, similar to the results with PAR4-AP N-terminal acetylation (**betaAla**-YPGKF-NH2 calcium EC_50_ = *n.d.*, β-arr-1 25.7 ± 4.8%, β-arr-2 31.8 ± 4.4%; **Inp**-YPGKF-NH_2_ calcium EC_50_ = *n.d.*, β-arr-1 32.0 ± 3.3%, β-arr-2 26.9 ± 3.0%). Together these data suggest a role for N-terminal charge localization in agonist activity.

### Effect of additional substitutions in positions 2-4 on PAR4-AP-mediated signalling

Previously, structure activity studies with PAR4 activating peptides revealed an apparent requirement for an aromatic residue in position 2 (tyrosine) and found that alterations to position 3 (proline) and position 4 (glycine) were not well tolerated (Faruqi et al., 2000). In the present study, we observed that substitution of either tyrosine, proline, or glycine with alanine reduced calcium signalling potency and β-arrestin recruitment, however none of these substitutions completely abolished the activity of the peptide (Fig. 1). Interestingly, substitution of either L-tyrosine or L-proline with the respective D-isomer significantly reduced the activity of the PAR4-AP (Fig. 2, Table 1). Together, these data suggest that while no individual residue is strictly required for agonist activity, the side chains of these residues and their positioning are important for agonist function. To further probe the contribution of these residues to the PAR4-AP, we generated peptides with alterations in the aromatic side chain of position 2, the backbone conformation of position 3, or position 4 (Fig. 5, Table 1).

**Figure 5.**
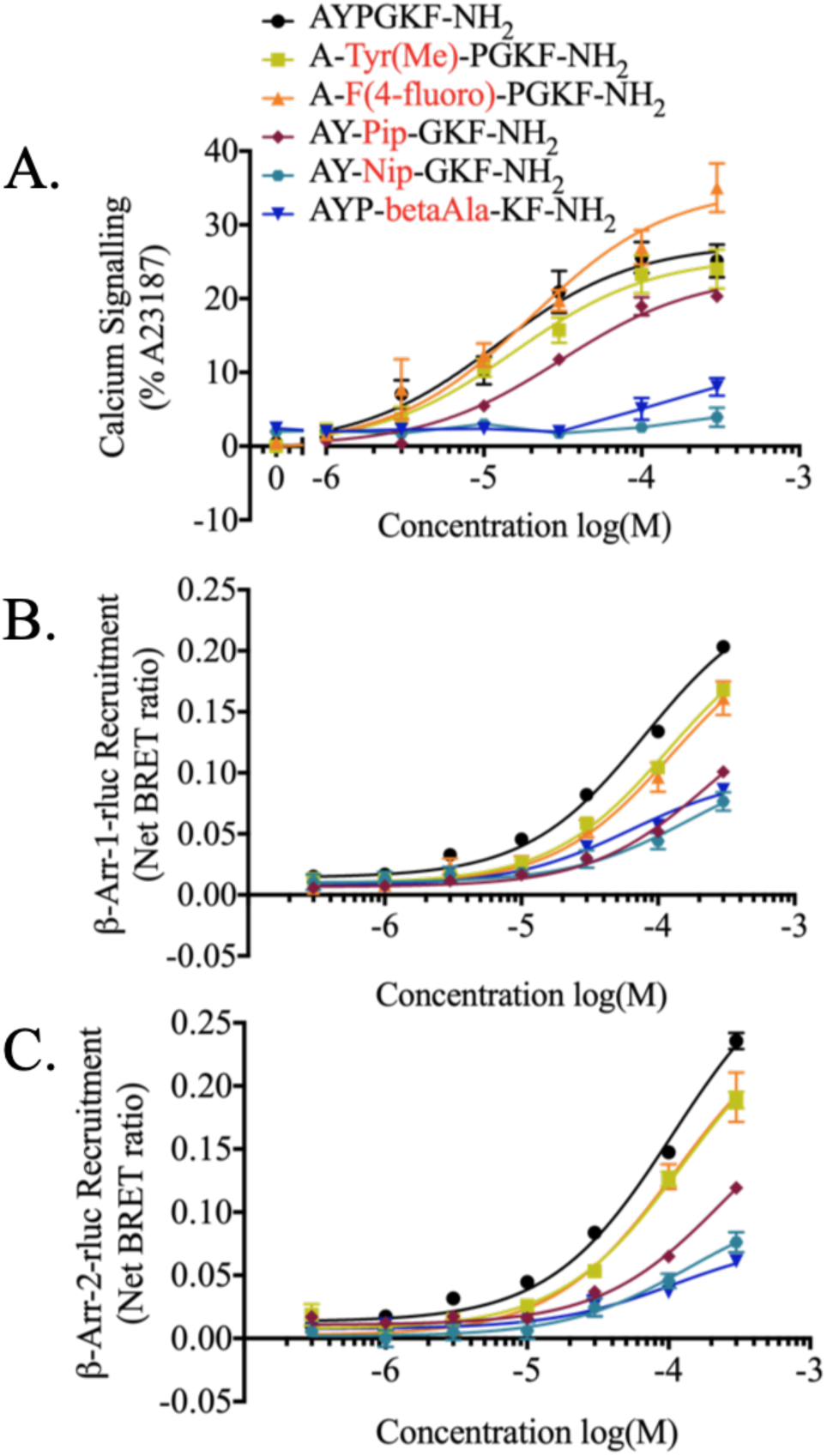
Calcium and β-arrestin recruitment to PAR4 in response to AYPGKF-NH_2_ or peptides with additional substitutions in tyrosine^2^, proline^3^, or glycine^4^. **(A) Agonist-stimulated calcium signalling.** Calcium signalling is shown as a percentage of the maximum calcium signalling obtained by application of calcium ionophore (A23187, 3 µM). (*n = 3-7)* **Agonist-stimulated recruitment of β-arrestin-1 (B) and β-arrestin-2 (C) to PAR4.** Concentration effect curves of BRET values from cells treated increasing concentrations of peptide are presented as normalized net BRET ratio. (*n = 3*).

Peptides with substitution of tyrosine^2^ with either 4-fluoro-L-phenylalanine [A-**F(4-fluoro)**-PGKF-NH2] or *O*-methyl-tyrosine [A-**Tyr(Me)**-PGKF-NH_2_] retained calcium signalling and β-arrestin recruitment with no significant changes in potency compared to the PAR4-AP. These data suggest that the side chain of tyrosine may be a site in which further modifications can be made to the peptide without affecting agonist potential; however, by comparing A**A**PGKF-NH_2_ the presence of an aromatic ring in position 2 is of importance. Given the results obtained with alanine or D-proline substitution of proline^3^ (Fig. 1 & 2, respectively), we furthered probed conformational contributions of this residue to the parental PAR4-AP. Proline was substituted with pipecolic acid (2-piperidinecarboxylic acid; Pip) which is structurally similar to proline but has an increased steric bulk and backbone elongation provided by a six-membered ring compared to the five-membered ring of proline. We observed partial agonism of calcium signalling in response to AY-**Pip**-GKF-NH_2_ stimulation with no significant difference in agonist potency (EC_50_ = 30.7 ± 4.9 µM) with a correlating partial agonism of β-arrestin recruitment (β-arr-1 38.8 ± 3.0%; β-arr-2 44.0 ± 2.7%). Thus, it appears that increasing steric bulk/backbone length may affect efficacy but still results in agonism. To further investigate the role of backbone conformation provided at position 3, proline was substituted with nipecotic acid (3-piperidinecarboxylic acid; Nip) which is similar to pipecotic acid, however, results in change of the peptide backbone direction (AY-**Nip**-GKF-NH_2_). With this modification, both calcium signalling (EC_50_ = *n.d.*) and β-arrestin recruitment (β-arr-1 32.9 ± 4.9%; β-arr-2 30.9 ± 4.5%) were significantly decreased in response to PAR4 stimulation. Elongation of the backbone in position four (glycine) by substitution with beta-alanine (AYP-**betaAla**-KF-NH_2_) was also significantly detrimental to both calcium signalling (EC_50_ = *n.d.*) and β-arrestin recruitment (β-arr-1 42.5 ± 1.8%, β-arr-2 24.9 ± 1.6%). Together these data suggest an important role for backbone conformation contributed by proline^3^ and glycine^4^ compactness in the PAR4-AP. As well, these data suggest peptide backbone length at proline^3^ and glycine^4^ has an important role; consistent with what we have observed with backbone elongation modifications at position 1 (**Inp**-YPGKF-NH_2_ and **betaAla**-YPGKF-NH_2_; Fig. 4, Table 1).

### Effect of additional substitutions in position 5 on PAR4-AP-mediated signalling responses

Previously, investigations into glutamine substitutions in the native tethered ligand mimicking peptide, GYPGQV, revealed that agonism is maintained when this position is substituted with a charged residue such as arginine or ornithine (Faruqi et al., 2000). In the present study, we have identified that substitution of position 5 (lysine) of the PAR4-AP (AYPGKF-NH_2_) with alanine does not significantly affect the agonist capacity of the peptide (Fig. 1). Further, we identified that substitution of L-lysine with D-lysine significantly decreases agonist stimulation of calcium signalling and β-arrestin recruitment (Fig. 2). To further probe the effect of positively charged residues in position 5, we generated several PAR4-AP analogues with lysine^5^ substitutions by either arginine or ornithine (Fig. 6, Table 1). Substitution of lysine^5^ with arginine resulted in a partial agonist with respect to calcium signalling but increased β-arrestin recruitment compared to the parental peptide (AYPG**R**F-NH_2_ calcium EC_50_ = 4.0 ± 1.1 µM, β-arr-1 154.2 ± 10.3%, β-arr-2 140.1 ± 9.4%). This substitution delocalizes the positive charge of the side chain compared to lysine and increases steric bulk. Interestingly, substitution with ornithine, which is similarly charged but has a shortened distance between the charge and the peptide backbone compared to lysine, also resulted in partial agonism of PAR4-mediated calcium signalling accompanied by a modest increase in the recruitment of β-arrestins (AYPG**O**F-NH_2_ calcium EC_50_ = 4.2 ± 1.1 µM, β-arr-1 121.7 ± 8.5%, β-arr-2 112.1 ± 7.7%). To further probe if steric bulk in the side chain of position 5 affects PAR4-AP potency, we increased steric bulk through methylation of the epsilon (*ε*) amine of lysine [AYPG-**(*N*_ε_-Me-K)**-F-NH_2_]. This substitution yielded a full agonist for calcium signalling (EC_50_ = 6.7 ± 1.9 µM) and a modestly more efficacious agonist for the recruitment of β-arrestins (β-arr-1 122.5 ± 12.4%; β-arr-2 116.7 ± 11.2%). All of these substitutions maintain a positive charge and are able to act as agonists for PAR4. To determine if charge in position 5 is required, we substituted lysine with citrulline, which maintains similar structure to arginine and lysine while removing a positive charge (AYPG-**Cit**-F-NH_2_). Interestingly, we observed partial agonism with respect to calcium signalling (EC_50_ = 5.4 ± 0.8 µM) with equipotent β-arrestin recruitment compared to the PAR4-AP, consistent with the alanine scan data for this position. Thus, positively charged residues in position 5 seem to contribute to agonism. Interestingly, we observed similar increases in β-arrestin recruitment and equipotent calcium signalling when stericity was either increased (AYPG**R**F-NH_2_) or decreased (AYPG**O**F-NH_2_), suggesting that the positive charge in this position is more important than the size of the side chain. Finally, although both the charge and size of the size chain have an effect, all modifications to position 5, with the exception of D-lysine (Fig. 2, Table 1) and *N*_α_-methyl-lysine (Fig. 3, Table 1) modifications, were generally well-tolerated and there appears to be modest freedom in the side-chain of this position. This makes the side chain a good candidate for further investigations of structure-activity relationships as well as a good region to label with an imagine modality.

**Figure 6.**
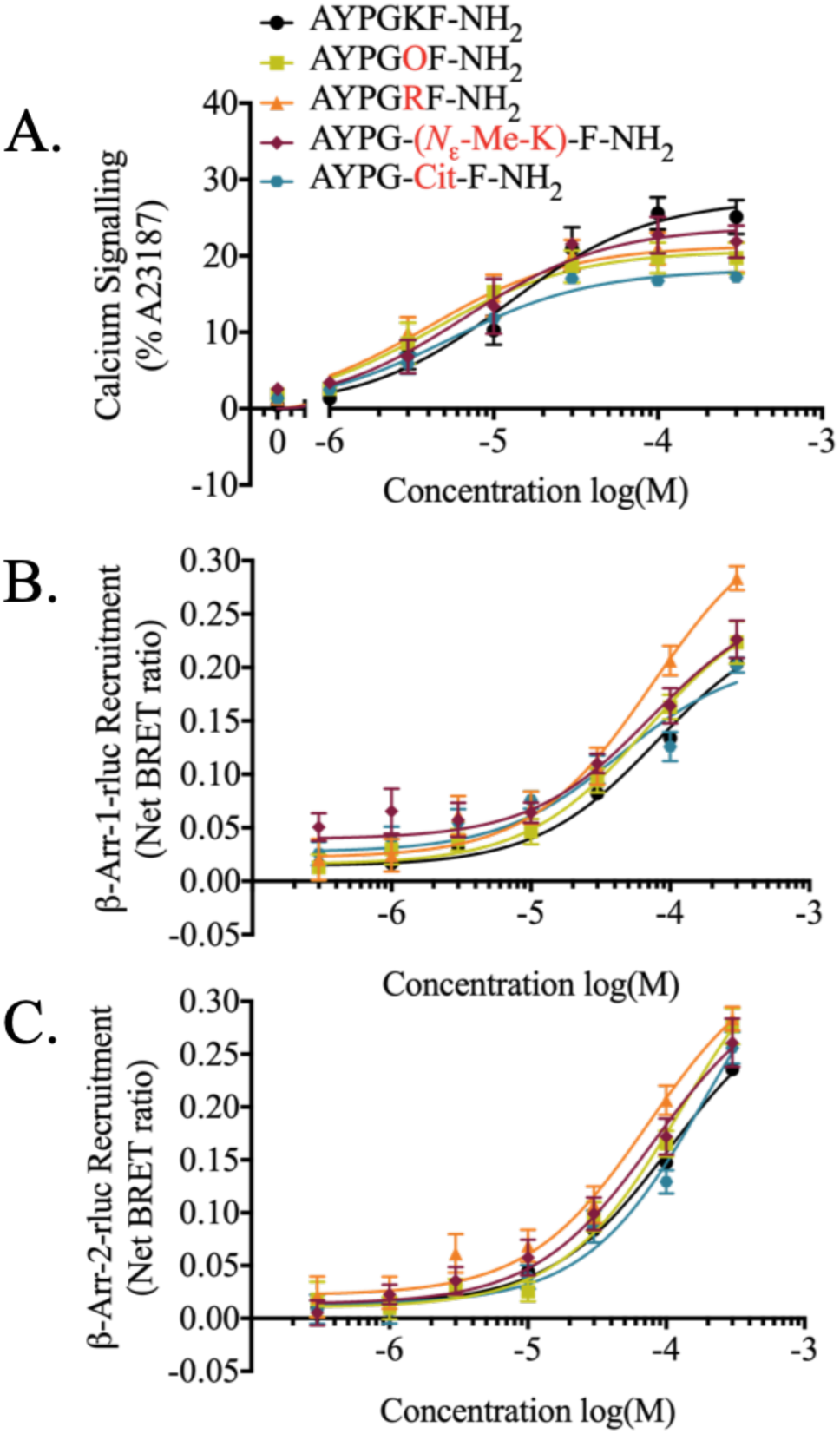
Calcium and β-arrestin recruitment to PAR4 in response to AYPGKF-NH_2_ or peptides with additional substitutions in lysine^5^. **(A) Agonist-stimulated calcium signalling.** Calcium signalling is shown as a percentage of the maximum calcium signalling obtained by application of calcium ionophore (A23187, 3 µM). (*n = 3-7)* **Agonist-stimulated recruitment of β-arrestin-1 (B) and β-arrestin-2 (C) to PAR4.** Concentration effect curves of BRET values from cells treated increasing concentrations of peptide are presented as normalized net BRET ratio. (*n = 3*).

### Effect of additional substitutions in position 6 on PAR4-AP-mediated signalling responses

Through our initial investigations into position 6 substitutions, we observed that replacement of phenylalanine^6^ with either alanine, D-phenylalanine, or *N*-methyl-phenylalanine had varied effects on PAR4-AP potency and efficacy. Previous investigations into position 6 substitutions in the native tethered ligand (GYPGQV) revealed the requirement for an aromatic residue, as substitution with non-aromatic residues such as lysine or ornithine significantly decreased or abolished the activity of the peptide (Faruqi et al., 2000). In addition to being an aromatic residue, phenylalanine is a hydrophobic residue so we investigated the impact of altering the hydrophobicity/hydrophilicity and stericity while maintaining an aromatic residue (Fig. 7, Table 1).

**Figure 7.**
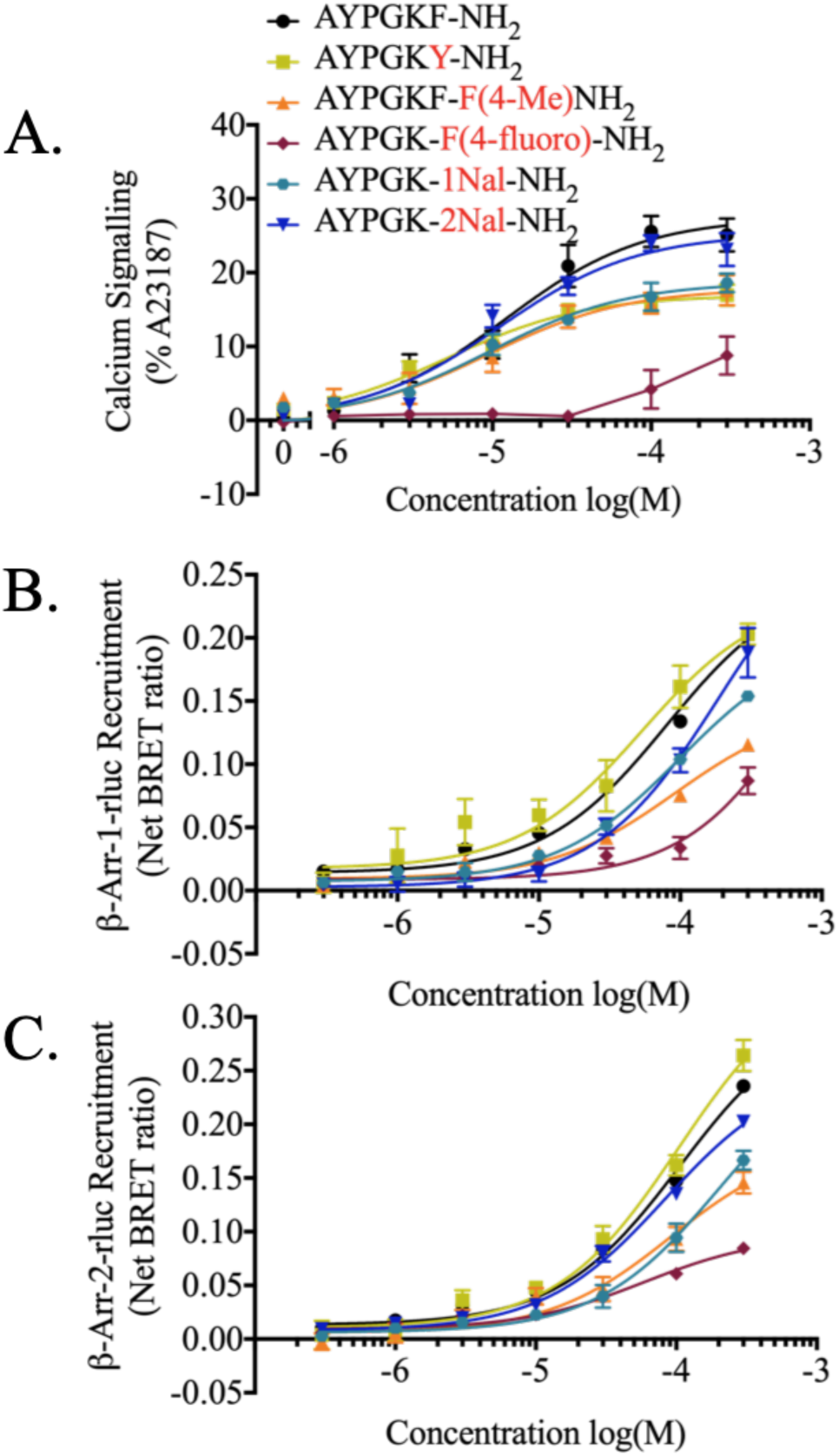
Calcium and β-arrestin recruitment to PAR4 in response to AYPGKF-NH_2_ or peptides with additional substitutions in phenylalanine^6^. **(A) Agonist-stimulated calcium signalling.** Calcium signalling is shown as a percentage of the maximum calcium signalling obtained by application of calcium ionophore (A23187, 3 µM). (*n = 3-7)* **Agonist-stimulated recruitment of β-arrestin-1 (B) and β-arrestin-2 (C) to PAR4.** Concentration effect curves of BRET values from cells treated increasing concentrations of peptide are presented as normalized net BRET ratio. (*n = 3*).

We investigated the effect of polar changes that decrease hydrophobicity. Substitution of phenylalanine^6^ with tyrosine results in the addition of a hydroxyl group to position 6 (AYPGK**Y**-NH_2_). We observed partial agonism of calcium signalling with no significant change in potency compared to the PAR4-AP (AYPGK**Y**-NH_2_, EC_50_ = 5.2 ± 1.0 µM). Interestingly, the peptide was able to recruit β-arrestins at least as well as the PAR4-AP (β-arr-1 120.5 ± 12.5%; β-arr-2 109.9 ± 11.4%). Further, replacing the tyrosine hydroxyl group with an electron withdrawing fluorine, by means of substitution with 4-fluoro-L-phenylalaine, caused significant impairment of agonist activity, both decreasing calcium signalling and significantly decreasing β-arrestin recruitment [AYPGK-**F(4-fluoro)**-NH_2_, calcium EC_50_ = *n.d.,* β-arr-1 25.2 ± 6.5%; β-arr-2 41.3 ± 5.9%]. The results of these two substitutions suggest that altering the polarity, while maintaining aromaticity, in position 6 reduces efficacy (Hoffmann et al., 2019). Since decreasing hydrophobicity did not increase PAR4-AP activity, we next investigated alterations that increase stericity and hydrophobicity. Substitution with 4-methyl-L-phenylalanine reduced the efficacy of the PAR4-AP to a partial agonist for both calcium signalling and β-arrestin recruitment [AYPGK-**F(4-Me)**-NH_2_ calcium EC_50_ = 9.4 ±3.0 µM, β-arr-1 56.4 ± 1.2%; β-arr-2 63.2 ± 1.1%]. Given these results, we wanted to assess the pharmacological impact of substituting phenylalanine with (1-naphthyl)-L-alanine (1Nal) and (2-naphthyl)-L-alanine (2Nal) which not only increase hydrophobicity but also have a modest increase in steric bulk and π-stacking potential. Substitution of phenylalanine^6^ with (1-naphthyl)-L-alanine (1Nal; AYPGK-**1Nal**-NH_2_) generated a partial agonist for both calcium signalling (EC_50_ = 9.6 ±1.9 µM) and β-arrestin recruitment (β-arr-1 77.5 ± 2.3%; β-arr-2 64.1 ± 2.0%) compared to PAR4-AP. Notably, substitution to (2-naphthyl)-L-alanine (2Nal; AYPGK-**2Nal**-NH_2_), which differs in orientation compared to (1-naphthyl)-L-alanine such that the alkyl chain is connected to position 2 of naphthyl group (instead of position one as in 1Nal), restored potency of the peptide to the levels observed with PAR4-AP. AYPGK-**2Nal**-NH_2_ was an equipotent agonist for calcium signalling (EC_50_ = 11.1 ± 2.3 µM) and β-arrestin recruitment (β-arr-1 77.0 ± 7.0%; β-arr-2 91.6 ± 6.4%). Together, these results indicate that hydrophobicity, stericity, and side chain positioning of the C-terminal residue all contributes to agonist peptide activity at PAR4.

### Probing antagonism of PAR4-AP-stimulated calcium signalling by calcium signalling null peptides

The failure of some substituted peptides to stimulate calcium signalling could reflect either a failure to bind or a loss of efficacy. To determine if such peptides can act as antagonists of PAR4-AP stimulated calcium signalling, we pre-incubated HEK-293 cells stably expressing PAR4-YFP prior to stimulation with PAR4-AP. Pre-treatment and competition with 100 μM of calcium null peptides were largely unable to affect the calcium response elicited by 30 μM PAR4-AP (AYPGKF-NH_2_) compared to vehicle treated controls. Interestingly, two calcium null peptides with substitution in position 3 or 4 were able to decrease calcium signals elicited by PAR4-AP. Calcium signalling in response to AYPGKF-NH_2_ (30 µM) was significantly reduced following pre-treatment with AY-**Nip**-GKF-NH_2_ (10.6 ± 0.9% of A23187; p<0.05) or AYP-**Sar**-KF-NH_2_ (13.0 ± 1.5% of A23187; p<0.05) compared to non-pre-treated control (20.8 ± 1.3% of A23187; p<0.05) (Fig. 8).

**Figure 8.**
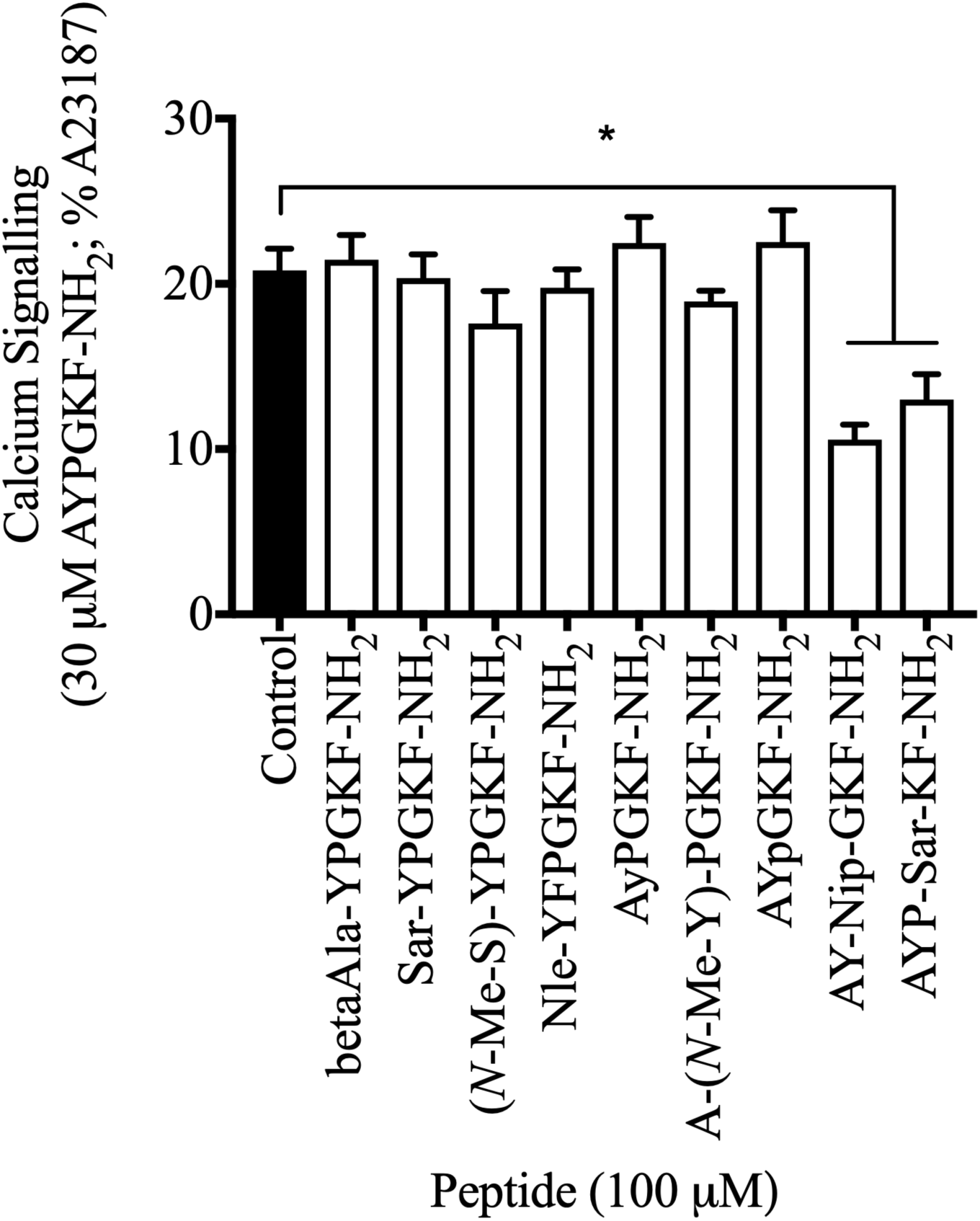
PAR4-AP (AYPGKF-NH_2_) stimulated calcium signalling responses following pre-treatment with calcium signalling null peptides. Calcium signalling (Fluorescence Em: 530) in HEK-293 cells stably expressing PAR4-YFP were recorded following 30 µM AYPGKF-NH_2_ addition after 30-minute pre-treatment incubation with 100 µM of calcium signalling-null peptides. Data are shown as a percentage of maximum response obtained in cells treated with the calcium ionophore A23187 (3 µM). Agonist peptide concentrations are in M. Pre-treatment with calcium-null peptides did not affect AYPGKF-NH_2_-stimulated calcium signalling in most cases with the exception of AY-Nip-GKF-NH_2_ and AYP-Sar-KF-NH_2_, which significantly decreased AYPGKF-NH_2_ stimulated calcium signalling compared to vehicle treated control. (*p > 0.05; *n = 3*)

### PAR4-dependent calcium signalling is Gα_q/11_-dependent

In order to verify that PAR4-dependent calcium signalling was G*α*_q/11_-dependent, HEK-293 cells stably expressing PAR4-YFP were stimulated with 100 µM AYPGKF-NH_2_ following pre-incubation with selective Gα_q/11_-inhibitor YM254890. YM254890 inhibited AYPGKF-NH_2_-stimulated calcium signalling in a concentration-dependent manner, with complete inhibition observed at 100 nM YM254890. The control vehicle (0.001% DMSO) had no effect on calcium signalling (Fig. 9).

**Figure 9.**
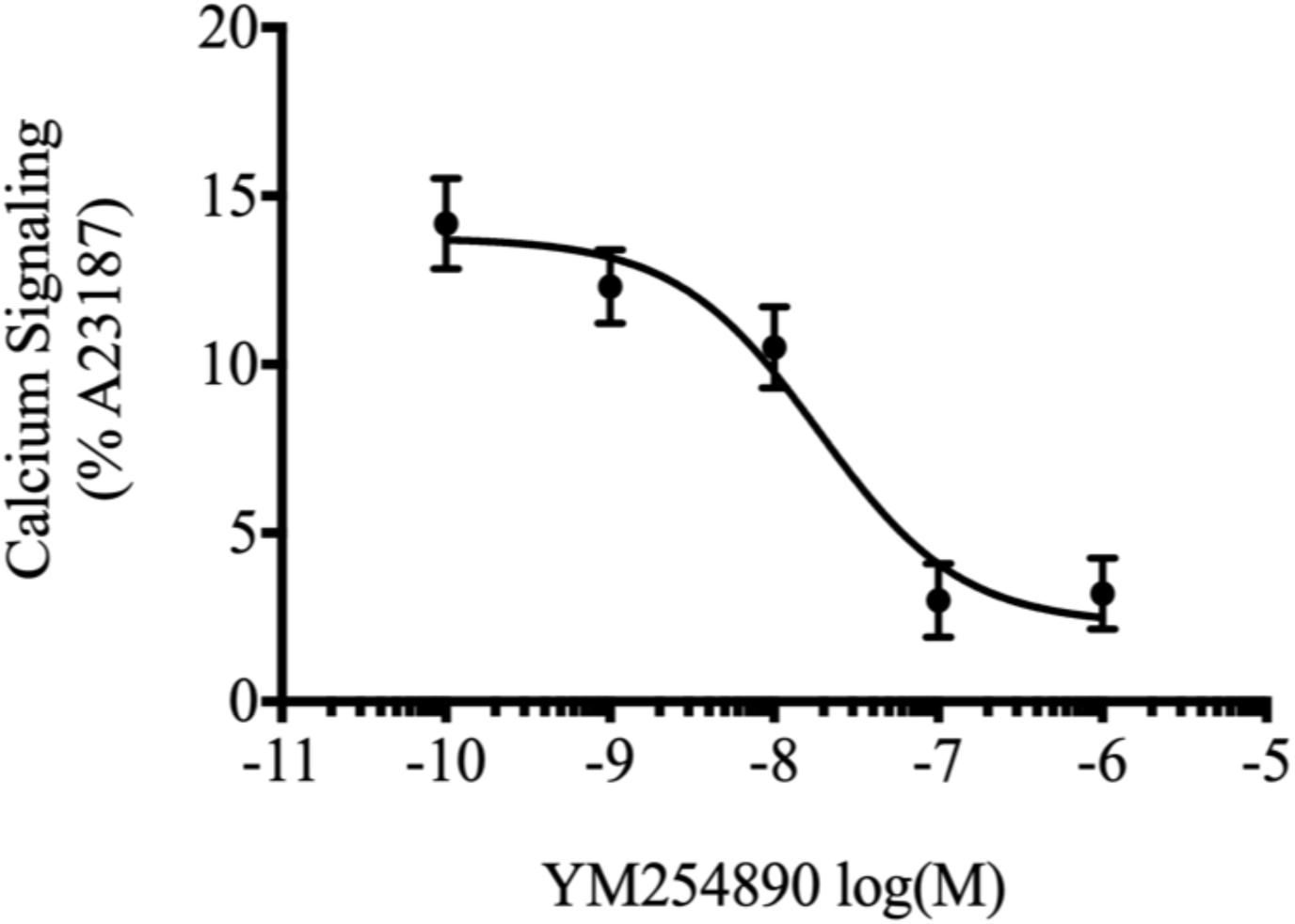
PAR4-mediated calcium signalling is Gα_q/11_-dependent. HEK-293 cells stably expressing PAR4 were pre-treated with either DMSO vehicle control (0.001%) or increasing concentration of the selective Gα_q/11_ inhibitor YM254890. Calcium signalling following stimulation with AYPGKF-NH_2_ (100µM) was recorded. PAR4-mediated calcium signalling was blocked by YM254890 at concentrations greater than 100 nM. (*n = 4*)

### Calcium signalling-null peptides do not activate MAPK signalling

Activation of G-protein and β-arrestin pathways downstream of GPCRs often converge on the activation of the p44/42 MAP kinase-signalling pathway (Jean-Charles et al., 2017). We therefore monitored phosphorylation of p44/42 MAPK in response to the 37 AYPGKF-NH_2_ derivative peptides. HEK-293 cells, stably expressing PAR4-YFP, were incubated with control vehicle (0.001% DMSO), 100 μM of AYPGKF-NH_2_, or derivative peptides (100 μM) for 10 minutes. Data were normalized as a ratio of phosphorylated-p44/42 (p-p44/42) to total-p44/42 (p44/42). Interestingly, we find that most peptides that were unable to stimulate calcium signalling were also unable to activate p44/42 MAPK signalling (*i.e.* A**y**PGKF-NH_2_, AY**p**GKF-NH_2,_ **(*N*-Me-S)**-YPGKF-NH_2_, **Inp**-YPGKF-NH_2_, **Sar**-YPGKF-NH_2_, AY-**Nip**-GKF-NH_2_, AYP-**Sar**-KF-NH_2_). Similarly, peptides that perform as partial agonists of Gα_q/11_-mediated calcium signalling were also partial agonists for p44/42 phosphorylation. Several peptides were able to cause a significantly greater increase in phosphorylation compared to the parental peptide (*i.e.* **Nle**-YPGKF-NH_2_, AYPG**O**F-NH_2,_ AYPG-**(*N*_ε_-Me-K)**-F-NH_2_, AYPG-**Cit**-F-NH_2_, AYPGK**Y**-NH_2_, AYPGK-**1Nal**-NH_2_,) (Fig. 10, Table 1).

**Figure 10.**
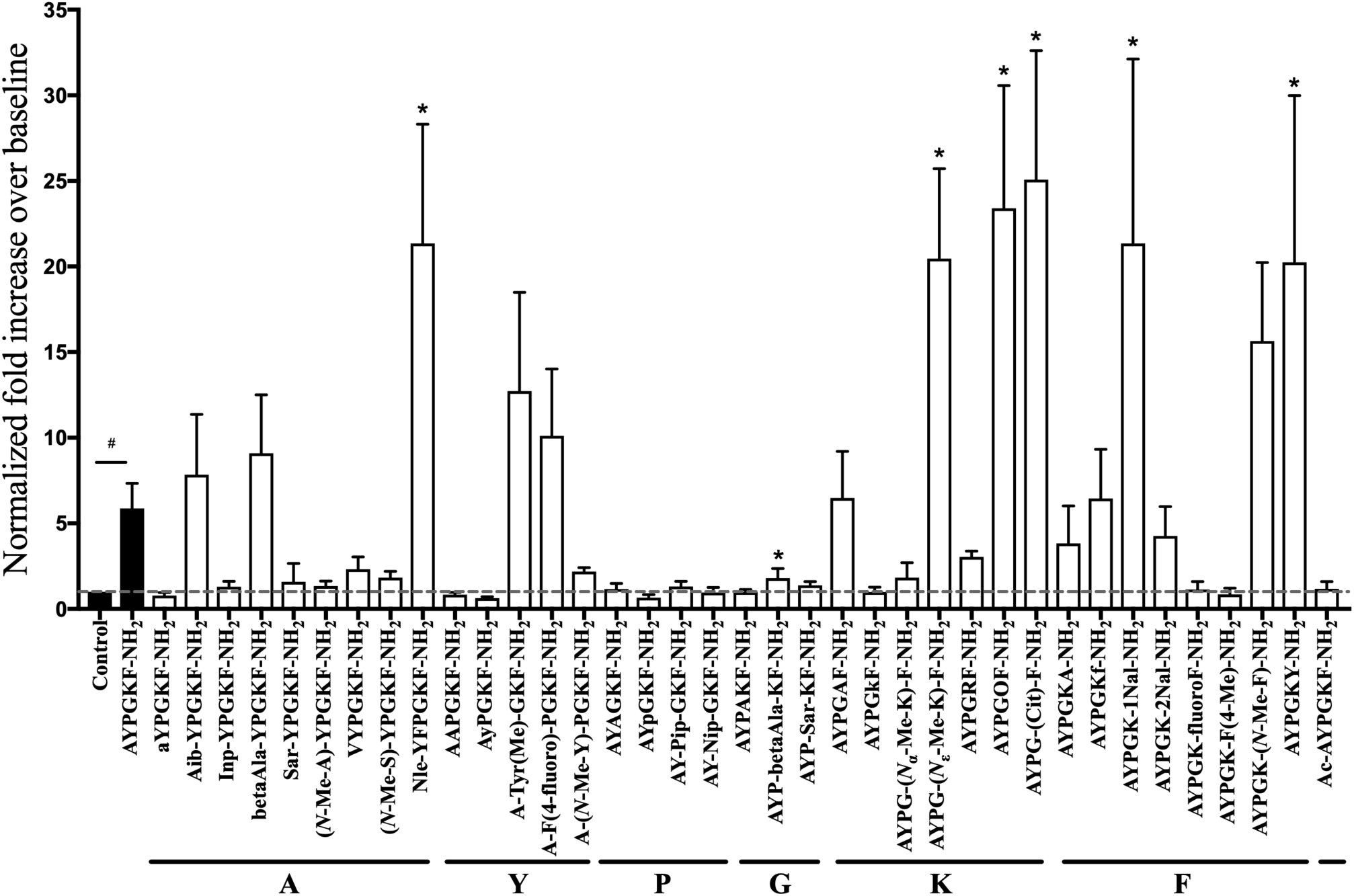
PAR4-mediated phosphorylation of p44/42(ERK) is attenuated upon stimulation of PAR4 with calcium-null peptide. HEK-293 cells stably expressing PAR4-YFP were stimulated with AYPGKF-NH_2_ (30 µM) or peptide (30 µM) from library of compounds and Western blot for p44/42 phosphorylation (p-p44/42). Data are shown as normalized fold increase over unstimulated control cells. Dashed baseline shows mean of unstimulated baseline for comparison. Dotted baseline shows mean response achieved following AYPGKF-NH_2_ stimulation. (# p > 0.05 AYPGKF-NH_2_ compared to untreated control; *p > 0.05 AYPGKF-NH_2_ compared to other peptide agonists; one-way ANOVA). Peptides that were unable to stimulate calcium signalling were also generally unable to stimulate phosphorylation of p44/42. Interestingly, several peptides were able to significantly increase phosphorylation of p44/42 compared to AYPGKF-NH_2_. (Representative blots shown in supplemental figure S2; *n* = *4-5*)

### PAR4-mediated MAPK signalling is Gα_q/11_-dependent and β-arrestin independent

In order to further probe the contribution of different pathways downstream of PAR4 to p44/42 phosphorylation, we examined the effective of blocking G*α*_q/11_-signalling with the specific antagonist YM254890 and examined the role of β-arrestins using a β-arrestin-1/-2 double knockout HEK-293 cell derived by CRISPR/Cas9 mediated targeting (Supp. Fig. 3). Pre-treatment of PAR4-YFP HEK-293 cells with 100 nM YM254890 for 20 minutes abolished PAR4-AP (30 μM) stimulated activation of MAP kinase signalling and subsequent phosphorylation of p-p44/42 at five minutes time (Fig. 11 A.). In contrast, there was no significant difference in the phosphorylation of p44/42 MAPK between wild-type HEK-293 and β-arrestin-1/-2 knockout HEK-293 cells stimulated with 30 μM PAR4-AP for 0-90 minutes (Fig. 11 B.).

**Figure 11.**
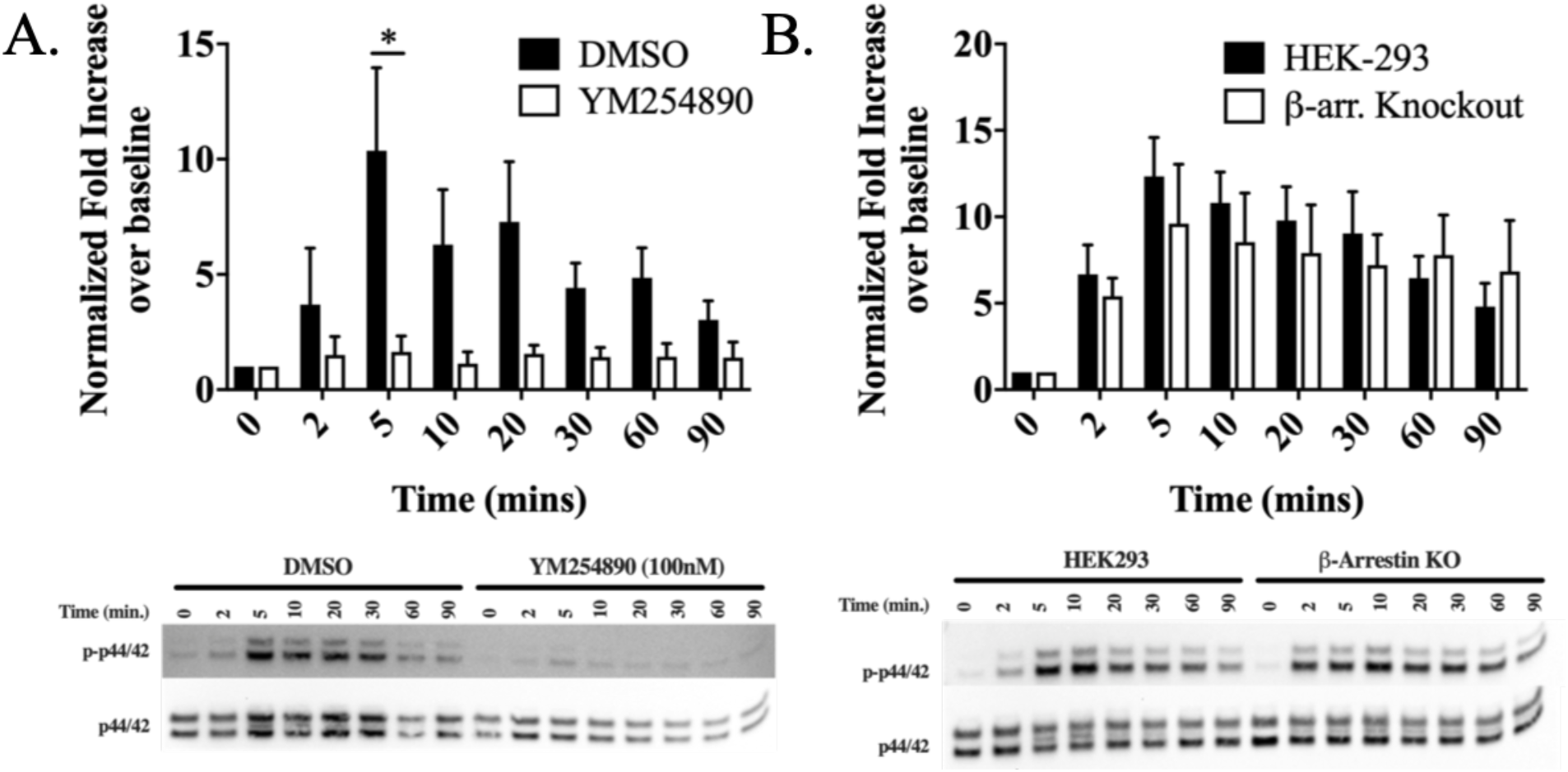
PAR4-mediated phosphorylation of p44/42 (ERK) is Gα_q/11_-dependent and β-arrestin-independent. To evaluate contribution of Gα_q/11_ and β-arrestin to p44/42 phosphorylation, p-p44/42 was monitored with western blot following stimulation with AYPGKF-NH_2_ (30 µM) for various time points. To evaluate contribution of Gα_q/11_-signalling cells were incubated with vehicle control (0.001% DMSO) or Gα_q/11_-inhibitor, YM254890 (100 nM) prior to agonist stimulation (A). To determine contribution of β-arrestin-signalling to p44/42 phosphorylation, HEK-293 and β-arrestin-1/2 CRISPR knockouts were assayed following agonist stimulation (B). Values are expressed as fold increase over unstimulated 0-minute time [(p-p44/22 / total p44/42) / (baseline p-p44/42 / total p44/42)]. Data are analysed with two-way ANOVA (*p > 0.05). We find that Gα_q/11_ inhibition with YM254890 results in a significant reduction of p44/42 phosphorylation. We observe no statistical difference in ERK activation between HEK-293 and β-arrestin-1/2 knockouts HEK-293 cells. (Representative blots shown; *n = 4*)

### Homology modelling of PAR4 and *in silico* docking of AYPGKF-NH_2_

In order to gain a better understanding of the ligand-binding pocket in PAR4 we turned to homology modelling and *in silico* docking experiments. A homology model of PAR4 was generated using known coordinates of thermostabilized human PAR2 crystal structure in complex with AZ8838 (2.8Å resolution, 5NDD.pdb, Cheng et al., 2017). Sequences of human PAR4 were aligned with the structure-resolved residues of PAR2 using Clustal Omega (Madeira et al., 2019; Sievers et al., 2011). We generated a homology model of wild-type PAR2 on the PAR2 structure, which contained modifications necessary for crystallization. Twenty human PAR4 homology models, using the wild-type human PAR2 model as a template, were generated in MODELLER (Webb and Sali, 2014). The PAR4 model with the lowest discrete optimized protein energy (DOPE) score was utilized for *in silico* analyses.

Docking AYPGKF-NH_2_ to the PAR4 homology model (GalaxyPepDock; Lee et al., 2015) revealed a number of highly predicted peptide-receptor interaction sites (Fig. 12 A.; Ser^67^, Tyr^157^, His^229^, Asp^230^, Leu^232^, Leu^234^, Asp^235^, Ala^238^, Gln^242^, His^306^, Tyr^307^, Pro^310^, Ser^311^, Ala^314^, Gly^316^, Tyr^319^, and Tyr^322^). Of these, the most frequently predicted interactions were receptor residues His^229^, Asp^230^, and Gln^242^. In peptide-docked models, Gln^242^ made contact with alanine, tyrosine, and lysine, with the most frequent prediction being with phenylalanine. His^229^ was predicted to interact with lysine and phenylalanine. Asp^230^ was frequently predicted to interact with lysine and phenylalanine and contacts with lysine appeared in 60% of predictions (Fig. 12 B.). Interestingly, when visualizing receptor residues predicted to interact with the peptide, we found that a large number of these residues were localized on extracellular loop 2 (ECL2) (Fig. 12 A. and B.).

**Figure 12.**
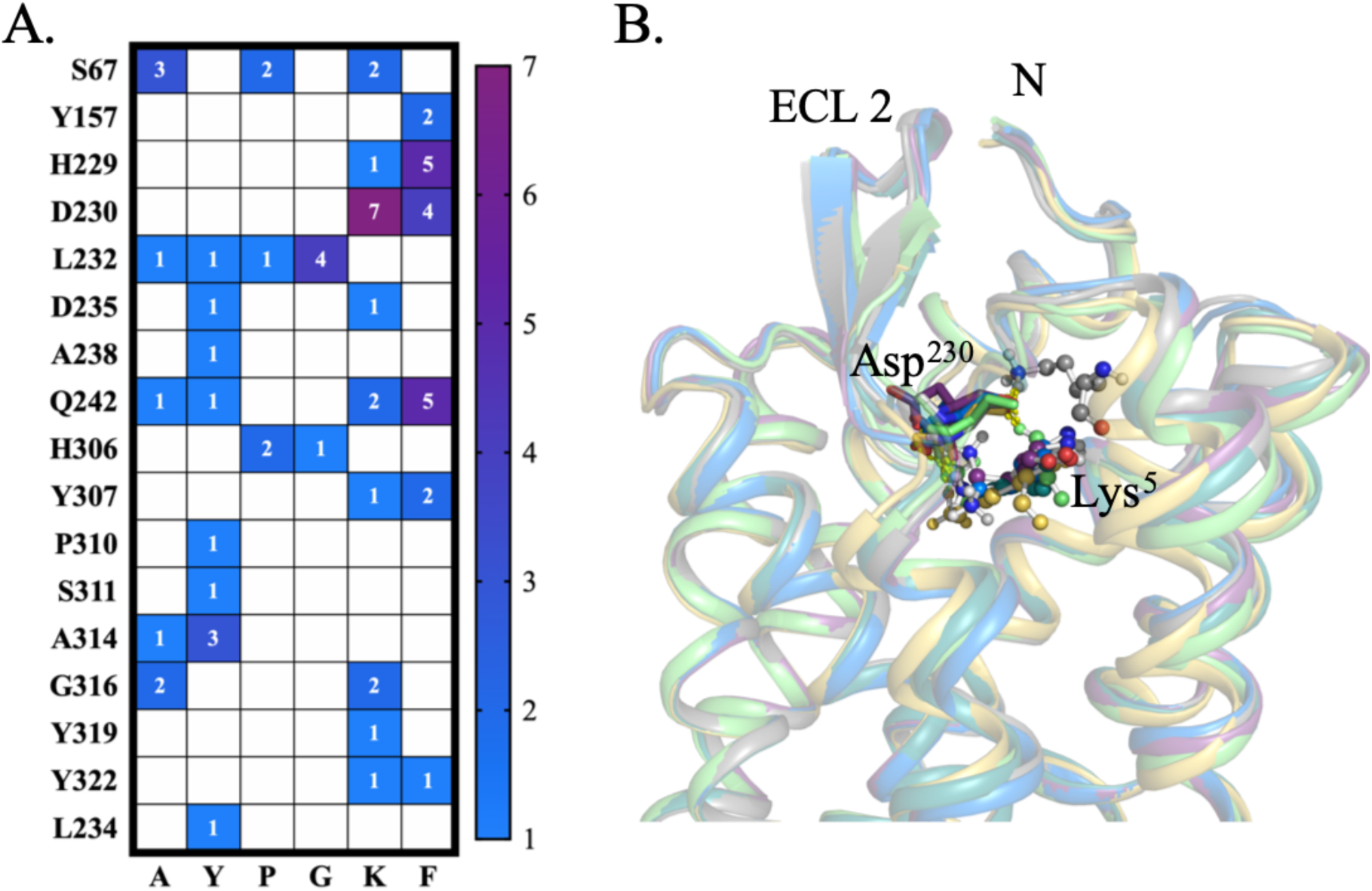
(A) Frequency table of PAR4/AYPGKF-NH_2_ interactions as predicted by *in silico* peptide docking (GalaxyPepDock). Numbers represent the frequency with which an atom in a PAR4-AP residue was predicted to interact with an atom of a PAR4 residue. **(B) Representation of the most highly predicted AYPGKF-NH_2_ binding site (Asp^230^, shown as sticks) interacting (shown as yellow dashed lines) with position 5 (Lys^5^; shown as ball and sticks) of AYPGKF-NH_2_**. Interaction of Lys^5^ and Asp^230^ was highly predicted in 7/10 models (B). Alignment of the receptor backbone (shown as transparent cartoon) reveals similar binding poses adopted by the side chain position of Asp^230^ and the position of Lys^5^ of AYPGKF-NH_2_.

### Calcium signalling and β-arrestin recruitment to predicted peptide binding-site mutant-PAR4

To experimentally examine *in silico* predictions, site-directed mutagenesis was performed introducing alanine substitutions at the three most highly predicted interacting residues, His^229^, Asp^230^, and Gln^242^. Additionally, we generated a double mutation, PAR4^H229A, D230A^, to determine if combined loss of His^229^ and Asp^230^ would result in additive detriment to receptor activation. We evaluated the functional consequences of these mutations by recording calcium signalling in response to AYPGKF-NH_2_ with all mutants. We observed no calcium response in cells expressing PAR4^D230A^-YFP or PAR4^H229A, D230A^-YFP mutant receptors up to 300 μM AYPGKF-NH_2_ (Fig. 13 A.). Similarly, peptide-induced β-arrestin-1/-2 recruitment to PAR4^D230A^-YFP and PAR4^H229A, D230A^-YFP was significantly reduced compared to the wild-type PAR4. Both PAR4^H229A^ and PAR4^Q242A^ mutations were fully able to recruit β-arrestin-1/-2 to the same levels observed with wild-type PAR4-YFP (Fig. 13 B. and C.).

**Figure 13.**
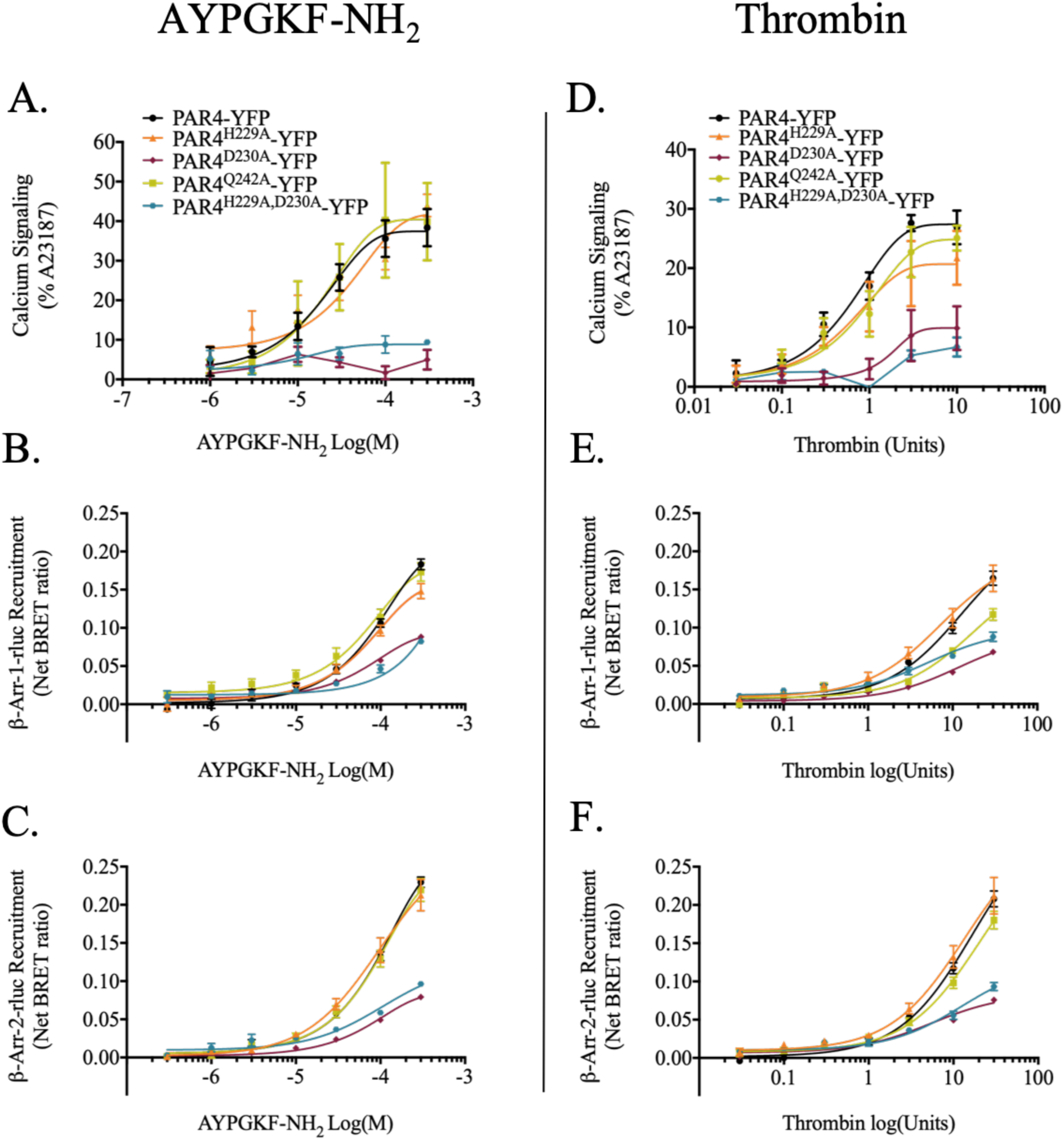
AYPGKF-NH_2_-stimulated calcium signalling and β-arrestin-1/-2 recruitment in PAR4 extracellular loop 2 mutants. Agonist-stimulated Calcium signalling. Calcium signalling in response to AYPGKF-NH_2_ (A) or thrombin (D) was recorded in HEK-293 cells transiently expressing PAR4-YFP, single site-directed mutant PAR4-YFP (PAR4^H229A^-YFP, PAR4^D230A^-YFP, PAR4^Q242A^-YFP), or double site-directed mutant PAR4-YFP (PAR4^H229A, D230A^-YFP, PAR4^H229D, A231D^-YFP). Data are shown as a percentage of maximum response obtained in cells treated with the calcium ionophore A23187 (3 µM) for each transient cell line, per experimental day. AYPGKF-NH_2_ concentration is in M and thrombin concentration is shown as units. (N=3-5) **Agonist-stimulated β-arrestin-1/-2 recruitment** HEK-293 cells were transiently transfected with BRET pair of either β-arrestin-1-rluc or -2-rluc and PAR4-YFP or site-directed mutant PAR4-YFP (described above) and agonist-stimulated arrestin recruitment to PAR4 was monitored. Cells were incubated with increasing concentration of either AYPGKF-NH_2_ (B, C) or thrombin (E, F) for 20 minutes and BRET values were recorded. BRET values from cells treated with HBSS were also collected to normalize the data. Renilla-luciferase substrate coelenterazine-h (5 µM) was added 10 minutes prior to data collection. Data are presented as the normalized net BRET ratio [eYFP/rluc - baseline (HBSS control)]. AYPGKF-NH_2_ concentration is in M and thrombin concentration is shown as units. (*n = 4*).

To investigate whether these sites are important for tethered ligand activation of the receptor, we stimulated cells expressing wildtype and mutant constructs with thrombin. Thrombin cleavage of the receptor reveals the native tethered ligand, GYPGQV (Xu et al., 1998). Comparable to activation with AYPGKF-NH_2_, we observed significantly diminished calcium signalling in HEK-293 cells expressing PAR4^D230A^-YFP and PAR4^H229A, D230A^-YFP mutants, even up to 10 units/mL thrombin (Fig. 13 D.). Additionally, β-arrestin-1/-2 recruitment was also impaired to PAR4^D230A^ and PAR4^H229A, D230A^ mutants (Fig. 13 E. and F.). We observed no appreciable difference in calcium signalling or β-arrestin recruitment when PAR4^H229A^-YFP and PAR4^Q242A^-YFP mutants were stimulated with thrombin, with the exception of a modest decrease in β-arrestin-1 recruitment to PAR4^Q242A^-YFP mutant receptor. Thus, Asp^230^ interactions with either PAR4-AP or the tethered ligand are critical for receptor activation.

### Platelet aggregation response to differential activation of PAR4 signalling

To determine the effects of targeted signalling observed with several peptides we investigated PAR4-mediated platelet aggregation in response to the β-arrestin biased, calcium signalling-null peptide, A**y**PGKF-NH_2,_ as well as a modestly more potent calcium agonist AYPG**R**F-NH_2_ (compared to AYPGKF-NH_2_). Thrombin (1 unit/mL) and 100 µM AYPGKF-NH_2_ were used as controls to ensure successfully washed platelet preparations. In response to thrombin stimulation, we observed a characteristic aggregation trace resulting in 100% aggregation at 1 unit/mL. In agreement with previously published data, AYPGKF-NH_2_ (100 µM) stimulation resulted in approximately 50% platelet aggregation (Ramachandran et al., 2017). We observed that platelet aggregation was not stimulated by 100 µM A**y**PGKF-NH_2_. Interestingly, when platelet preparations were stimulated with AYPG**R**F-NH_2_, with which we observed a modest increase in potency in calcium signalling assays, enhanced β-arrestin recruitment, and an increased phosphorylation of p44/42, we observed platelet aggregation response comparable to levels observed with thrombin (1 unit/mL) (Fig. 14).

**Figure 14.**
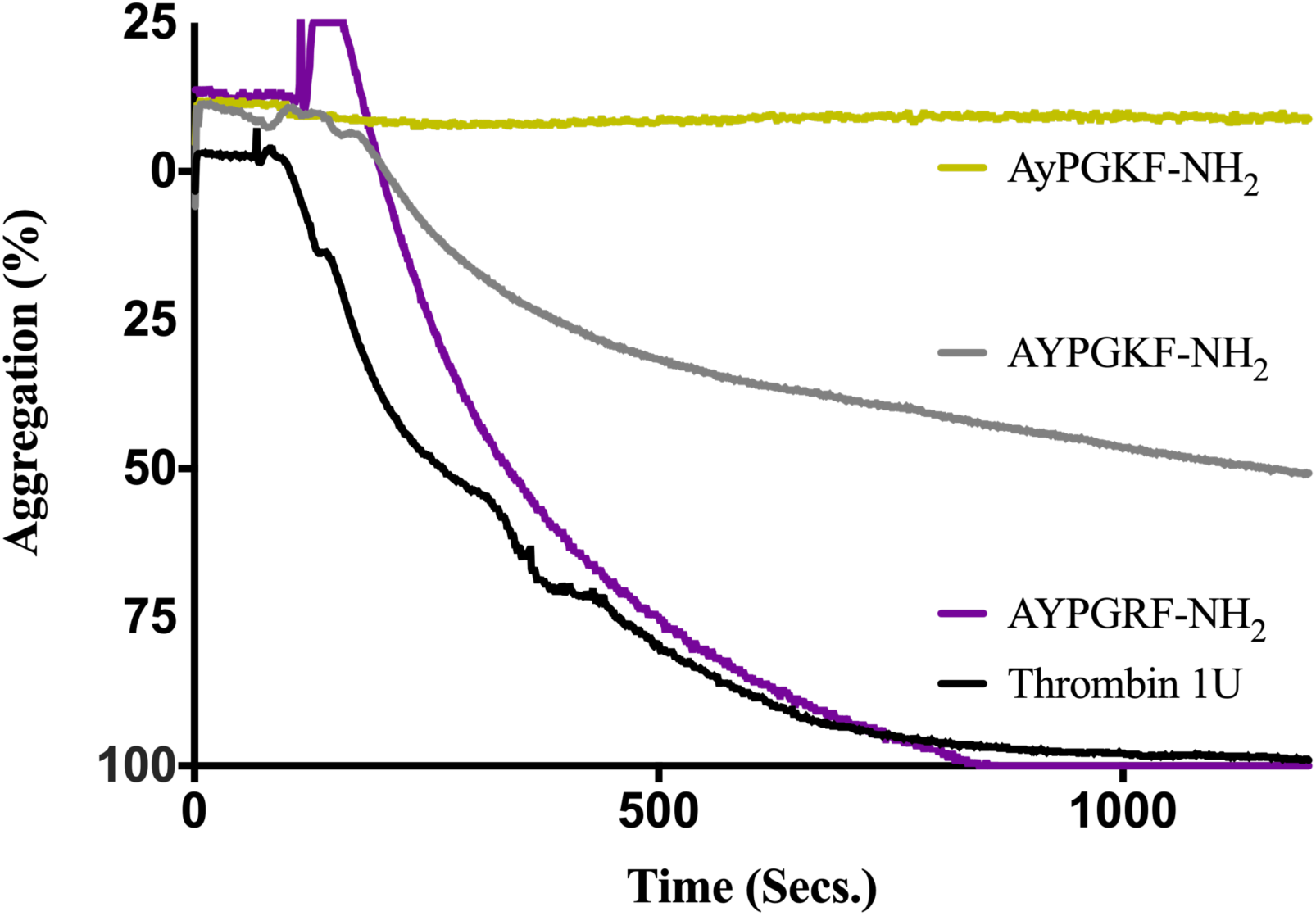
PAR4 agonist-mediated platelet aggregation. Both thrombin (1 unit/mL; shown in black) and AYPGKF-NH_2_ (100 µM; shown in grey) initiated PAR4-dependent rat platelet aggregation. We observed that stimulation of washed platelet samples with a calcium signalling null, β-arrestin biased peptide, AyPGKF-NH_2_ (100 µM; shown in yellow) did not stimulate platelet aggregation. Further, stimulation of washed platelets with the peptide agonist AYPGRF-NH_2_ (100 µM; shown in purple), caused a robust platelet aggregation comparable to the response with 1unit/mL thrombin response (*n = 2-4*).

## Discussion

PAR4 is emerging as a novel anti-platelet drug target. Here we examine structure activity requirements for PAR4 activation of calcium signalling, β-arrestin recruitment and MAP kinase activation. PAR cleavage by a number of different enzymes has been reported (Soh et al., 2010). Canonically, thrombin has been described as a PAR1 and PAR4 activator while trypsin is described as a PAR2 and PAR4 activator. While PAR3 can be cleaved by thrombin, its ability to act as a signalling receptor remains ambiguous. In each case, cleavage by serine proteases occurs at an arginine-serine site which reveals a unique tethered-ligand specific to each PAR (Hollenberg, 2003) - human PAR1-SFLLRNP… (Liu et al., 1991; Vu et al., 1991), human PAR2-SLIGKV… (Al-Ani et al., 2002; Nystedt et al., 1994) and human PAR4-GYPGQV… (Kahn et al., 1998; Xu et al., 1998). While these tethered-ligand sequences are diverse, they retain some highly-conserved properties including a highly-conserved charged residue in position five of the revealed tethered-ligands (arginine in PAR1; lysine in PAR2). Additionally, PAR2 and PAR4 have a conserved glycine in position three and valine in position six of their tethered-ligands. The similarities of the proteolytically-revealed tethered ligands is also observed in the optimized tethered-ligand-mimicking synthetic agonist peptides. Hydrophobic residues leucine, isoleucine, and proline are preferred in position three of PAR1, PAR2, and PAR4 agonist peptides, respectively. Additionally, there is a basic residue in position five of the tethered ligand peptide in all three receptors (arginine in PAR1 and PAR2 and lysine in PAR4 activating peptides). These conserved residue properties highlight some of the key requirements that a PAR4 agonist peptide must meet. Previous structure-activity work starting with the proteolytically-revealed sequences of human (GYPGQV) and murine (GYPGKF) PAR4 identified the hexapeptide AYPGKF-NH_2_ or SYPGKF-NH_2_ to be potent and selective agonists for PAR4 (Faruqi et al., 2000). In this study we started with the hexapeptide AYPGKF-NH_2_, as our parental peptide agonist, and incorporated various modifications at each of the six positions in this peptide to investigate determinants of agonist-induced PAR4-coupling to G*α*_q/11_, MAP kinase and β-arrestin-1/-2 recruitment to guide development of biased agonists of PAR4.

Previously, it was demonstrated that substitution of alanine or serine in position 1 of the GYPGKF-NH_2_ activating peptide retained the ability to trigger tritiated inositol 1,4,5-triphosphate release to a level comparable to that observed with thrombin activation and much higher than release caused by the GYPGKF-NH_2_ (Faruqi et al., 2000). Interestingly, substitution of position 1 with threonine was reported to significantly reduce the potency of the activating peptide (Faruqi et al., 2000). In keeping with results from previous studies, we find a requirement for small aliphatic side chains in position 1 of PAR4 activating peptides, with substitutions that increased bulk in this position consistently resulting in poor agonists for both calcium signalling and β-arrestin recruitment (*N*-Me-A Fig. 3; *N*-Me-S, Aib, Inp, V, Nle Fig 4; Table 1). Additionally, we observed that both stereochemical inversion of L-alanine to D-alanine (D-alanine, Fig. 2) and backbone elongation (betaAla., Fig. 4) were detrimental to both calcium signalling and β-arrestin recruitment. Additionally, we observed that capping of the N-terminal charge via acetylation significantly decreases agonist potency. Together these results indicate that a positive charge and the localization of that charge in addition to precise side-chain positioning at position 1 are critical to effective PAR4-agonism.

The aromatic sidechain, and to a lesser extent the hydroxyl group, of the tyrosine^2^ residue of the parental peptide also proved to be important for peptide activity with alanine or D-isomer substitutions resulting in decreased activity of all pathways tested. These results are consistent with previous reports that replacement of tyrosine with other aromatic residues, such as phenylalanine, significantly reduced both the selectivity and the potency of the activating peptide (Faruqi et al., 2000). Interestingly, substituting L-tyrosine^2^ for either D-tyrosine (Fig. 2 A., Fig. 10) or *N*-methyl-tyrosine (Fig. 3 A. & 10) resulted in significant decreases in calcium and p44/42 signalling, while having relatively modest effects on β-arrestin recruitment (Fig, 2 B. & C., Fig. 3 B. & C., respectively; Table 1). Also, substitution of tyrosine with either 4-fluoro-phenylalanine or *O*-methyl-tyrosine did not significantly affect calcium transients, p44/42 phosphorylation or β-arrestin recruitment in comparison to AYPGKF-NH_2_ (Fig. 5 & 10; Table 1). Our findings, together with previously reported data (Faruqi et al., 2000), point to the necessity of an aromatic residue in position 2 to maintain agonist potency. Further, our data reveal that backbone orientation in position 2 is a key determinant of agonist potency. Future experiments fine-tuning the aromatic substituents may be beneficial for peptide analogue development.

Our data reveal that alterations affecting proline backbone conformation or glycine residue compactness are not well-tolerated, as these decreased calcium signalling, p44/42 MAP kinase signalling, and β-arrestin recruitment to PAR4. Calcium signalling has previously been reported to be negatively impacted by alterations to either of these two residues (Faruqi et al., 2000). Both proline and glycine residues are frequently found in β-turns, in which proline provides a characteristic kink, and the available data support the notion that the conformation(s) (proline^3^ and glycine^4^) and rigidity (proline^3^) of PAR4 agonists permitted by these residues is/are important for receptor activation. To test this hypothesis further, we made several substitutions in position 3 (proline) aimed at evaluating favoured backbone conformations. We observed that a return to a predominantly trans-amide conformation by substitution of proline with alanine (Fig. 1), results in decreased potency in all pathways studied. Interestingly, this substitution did not abolish signalling but did significantly decrease agonist potency, which suggests that the backbone conformation of proline may be necessary for receptor binding but not receptor activation. Substitution of proline to pipecolic acid (Pip., Fig. 5), which has a similar peptide backbone orientation to proline with a six-membered aromatic ring, decreased β-arrestin recruitment with relatively modest effects for calcium signalling activation. Interestingly, substitution of proline with nipecotic acid (Nip., Fig. 5) abolished both calcium signalling and p44/42 phosphorylation while also significantly reducing β-arrestin recruitment. Nipecotic acid is similar to pipecolic acid with the exception that nipecotic acid changes the direction of the peptide backbone compared to that adopted by proline, while pipecolic acid favours a similar backbone directionality and conformation similar to proline. Interestingly, we observed a similar pharmacological profile observed with AY-**Nip**-GKF-NH_2_ when we substituted D-proline in position 3, which also changes the backbone orientation from that of the of the parent peptide. These findings demonstrate that the backbone conformation provided by proline in position 3 of the parental peptide is necessary for agonism of PAR4 signalling pathways. Further, substitution of a proline^3^ with alanine (AY**A**GKF-NH_2_, Fig. 2) significantly and detrimentally altered the pharmacological profile compared to the parental peptide PAR4-AP, suggesting the possibility that agonist competency contributed by position 3 is largely conformational and not chemical. Substitution of sarcosine or beta-alanine at position 4 also negatively affected all signalling pathways, while substitution of glycine^4^ with alanine resulted in a decrease in potency with calcium signalling as well as a reduction of β-arrestin recruitment. Together these substitutions reveal the requirement for a small flexible residue in position 4, consistent with previous reports (Faruqi et al., 2000).

It has been previously demonstrated that substitution of position 5 glutamine in the native tethered ligand (GYPGQV) with arginine or ornithine results in decreased potency in the PAR4 agonist peptide for calcium signalling pathways (Faruqi et al., 2000). However, there is a basic residue found in position 5 of PAR1 (arginine, TFLL**R**), PAR2 (arginine, SLIG**R**L), and PAR4 (lysine, AYPG**K**F) tethered-ligand mimicking peptides. Given the conservation of charged residues in position 5 of PAR tethered-ligand mimicking peptides, we generated PAR4-AP derivative peptides with lysine^5^ substitutions that maintained or altered charge in this position to determine if this is a requirement for activity at PAR4. Substitution of lysine^5^ with alanine had no appreciable effect on calcium or MAP kinase signalling, while resulting in decreased β-arrestin recruitment (Fig. 1). Substitution of L-lysine^5^ with D-lysine, however, significantly reduced potency and efficacy in calcium signalling and β-arrestin recruitment assays and furthermore resulted in a loss of stimulation of MAP kinase signalling (Fig. 2). We continued our exploration of this position with an *N*_α_-methyl-lysine substitution, which increases steric bulk through the addition of a methyl group to the backbone while also removing a hydrogen bond donor and increasing the amount of peptide in the cis-transformation of the amide bond between position 4 and 5. We found this substitution detrimental to calcium signalling but advantageous for β-arrestin recruitment, resulting in comparable maximal levels of recruitment to those observed with PAR4-AP but at lower concentrations (Fig. 3). Thus, backbone conformation and bulk at position 5 are important for agonism of calcium signalling and MAP kinase activation, while possibly being an important site for modification in the search for β-arrestin biased ligands.

We next turned our attention to analogues that resemble the side chain of lysine^5^, with a specific interest on structure-activity in addition to ascertaining the importance of charge in this residue. In contrast to the observation with similar substitutions to the tethered-ligand-mimicking peptide (GYPGQV) (Faruqi et al., 2000) wherein signalling was decreased compared to PAR4-AP, we found that both calcium signalling and β-arrestin recruitment are only modestly impacted by position 5 substitution to ornithine in the PAR4-AP, which decreases steric bulk and shortens the distance between the backbone and the positively charged amine compared to lysine (AYPG**O**F-NH_2_; Fig. 6). Interestingly, however, we observed significantly increased p44/42 phosphorylation in response to AYPG**O**F-NH_2_ stimulation (Fig. 10). This is somewhat puzzling since we found p44/42 phosphorylation to be G*α*_q/11_-dependent (Fig. 11); however, it is important to note that MAP kinase and other cellular signalling pathways downstream of receptor activation are subject to amplification/deamplification processes within the cell (Chidiac, 2016) and these observations may point to additional differences in signalling stimulated by this peptide that needs to be examined. We further probed stericity in this position through increasing bulk with substitution of lysine^5^ with arginine (AYPG**R**F-NH_2_). Interestingly, this substitution also results in a delocalization of charge compared to the charge on the lysine^5^ side chain. This modification provided a similar calcium signalling profile compared to that observed with ornithine substitution, however, it also resulted in a significant increase of β-arrestin recruitment compared to the parental PAR4-AP. Interestingly, there was no significant change in p44/42 phosphorylation unlike what was observed with ornithine substitution. Another peptide with increased steric bulk, *N*_ε_-methyl-lysine substitution in position 5, improved calcium signalling potency, modestly improved β-arrestin recruitment, and significantly increased p44/42 phosphorylation [AYPG-**(*N*_ε_-Me-K)**-F-NH_2_, Fig. 6]. We further investigated whether charge was a significant determinant of agonism by substituting lysine^5^ with citrulline (AYPG-**Cit**-F-NH_2_), which lacks charge but retains a similar structure compared to arginine and, to a lesser extent, lysine. We observed decreased calcium, equivalent β-arrestin recruitment, and significantly increased p44/42 phosphorylation compared to the parental peptide (Fig. 6 & 10). Together these data reveal that changes to side chain charge, bond donors/acceptors, and steric bulk result in varied pharmacological responses that may be exploited to bias signalling responses in favour of β-arrestin recruitment (AYPG**R**F-NH_2_) or p44/42 phosphorylation (AYPG**O**F-NH_2_; AYPG-**(*N*_ε_-Me-K)**-F-NH_2_) with modest improvements to potency in calcium signalling. Interestingly, we observed that charge is not wholly required for receptor activation as both alanine and citrulline substitutions resulted in agonism of PAR4.

Finally, our structure-activity investigation turned to position 6, phenylalanine, of the parental peptide. Given the significant impact that alterations to position 2 (tyrosine) had on agonist potency, we predicted that there may be a pi-pi (*π*-*π*) stacking interaction between tyrosine and phenylalanine which is supported by the backbone kink provided by proline in position 3. Substitution of position 6 with alanine or D-phenylalanine reduced calcium signalling potency while reducing β-arrestin recruitment and p44/42 phosphorylation. These alterations suggest that phenylalanine in position 6 is not essential for PAR4 activation by agonist peptide but is important for agonist potency and full activation of all signalling pathways. Methylation of the backbone nitrogen in phenylalanine (AYPGK-**(*N*-Me-F)**-NH_2_) resulted in improved activation of calcium signalling, β-arrestin recruitment, and p44/42 phosphorylation compared to AYPGKF-NH_2_ (Fig. 3). Addition of either methyl (AYPGK-**F(4-Me)**-NH_2_; Fig. 7), fluorine (AYPGK-**F(4-fluoro)**-NH_2_; Fig. 7), or hydroxyl (AYPGK**Y**-NH_2_; Fig. 7) off of the *para-*position of the aromatic side chain of phenylalanine was deleterious to all signalling pathways studied. Substitution of unnatural residues that increase the size of steric bulk and *π*-stacking potential yielded mixed results based on the orientation of the naphthyl. Substitution of phenylalanine with (1-naphthyl)-L-alanine (1Nal) increased p44/42 phosphorylation but decreased both calcium signalling and β-arrestin recruitment (Fig. 7). Changing the alkyl chain connection from position 1 to position 2 of the naphthyl (AYPGK-2Nal-NH_2_; Fig 7) resulted in activity similar to AYPGKF-NH_2_. With (2-naphthyl)-L-alanine showing some flexibility in the steric bulk of position 6, it would be interesting for future work to investigate increasing the *π*-stacking potential at position 2 while simultaneously increasing the *π*-stacking potential at position 6, since we hypothesize that these two residues’ aromatic regions of the PAR4-AP may interact.

To determine if any of the calcium signalling-null peptides were able to act as antagonists, cells were pre-treated with calcium null peptides and subsequently stimulated with AYPGKF-NH_2_ to see if the parental peptide was able to still elicit a signal. We observed that both AY-**Nip**-GKF-NH_2_ and AYP-**Sar**-KF-NH_2_ were both able to significantly decrease the calcium signal elicited by PAR4-AP (Fig. 8). Interestingly, each of these substitutions resulted in ablation of both calcium and p44/42 signalling as well as significantly decreasing β-arrestin recruitment. These results may indicate that targeting PAR4 with small molecules mimicking position 3 and 4 dipeptides may provide a starting place for the design of novel PAR4 antagonists. Additionally, peptides that further explore modifications to position 3 and 4 could be investigated for potent PAR4 antagonists.

In platelets, it has previously been demonstrated that PAR4-mediated calcium mobilization induces activation and aggregation (Edelstein et al., 2014; Holinstat et al., 2006). Both thrombin and AYPGKF-NH_2_ initiated PAR4-dependent platelet aggregation to levels comparable to those previously reported in the literature in the same rat platelet system (Ramachandran et al., 2017). As was hypothesized, we observed that stimulation of washed platelet samples with a calcium signalling null, β-arrestin biased peptide, A**y**PGKF-NH_2_ (100 µM) did not stimulate platelet aggregation (Fig. 14). Further, stimulation of washed platelets with the agonist peptide, AYPG**R**F-NH_2_ (Fig. 6), enhanced platelet aggregation to levels comparable to those observed with thrombin activation of PAR4 (Fig. 14). AYPG**R**F-NH_2_stimulates calcium signalling to levels comparable to those seen with PAR4-AP but triggers more β-arrestin-1 recruitment than the parental peptide. These data suggest that calcium signalling is essential for triggering platelet activation, but arrestin recruitment may also play a role.

Having ascertained some of the governing peptide residue characteristics enabling PAR4 activation and validated their signalling consequences in an *ex vivo* system, we turned our attention to investigating the binding site of the receptor. The extracellular loops of many GPCRs have been demonstrated to be important facilitators of ligand entry into the orthosteric site, with ECL2 possessing the most diversity in sequence and structure (Venkatakrishnan et al., 2014; Woolley and Conner, 2017). Analysis of common mechanisms of GPCR ligand interaction derived from crystal structure and biochemical experiments reveal a possible role of ECL2 secondary loop in binding and facilitating entry, while the diversity of residue sequences across class A GPCRs is thought to confer their ligand specificity (Venkatakrishnan et al., 2014; Woolley and Conner, 2017). In many class A GPCRs, a disulfide bond between ECL2 and a transmembrane 3 cysteine (Cys^3.25^, Ballesteros-Weinstein numbering) is thought to stabilize the extracellular topology such that ligand recognition at ECL2 and subsequent orthosteric binding can occur (Wheatley et al., 2012). All PAR receptors possess this canonical TM3 Cys^3.25^ (PAR1 Cys175^3.25^, PAR2 Cys148^3.25^, PAR3 Cys166^3.25^, PAR4 Cys149^3.25^) and residues adjacent to this conserved cysteine are important for ligand binding and receptor activation. Investigations into the role of PAR ECL2 domains have previously demonstrated that all PAR receptors possess a homologous CHDxL motif (…CHDVL… - PAR1^C254-L258^ and PAR2^C226-L230^; …CHDVH… PAR3^C245-L249^ …CHDAL; … PAR4^C226-L230^) in their ECL2 (Al-Ani et al., 1999; Xu et al., 1998). Given that the activating peptides of PAR1 and PAR2 (Al-Ani et al., 1999; Lerner et al., 1996) are similar (SFLLRN, PAR1; SLIGRL, PAR2), additional studies were conducted that revealed that ligand specificity is conferred by acidic residues distal to the CHDxL motif (PAR1, N^259^ETL^261^; PAR2 P^231^EQL^234^; (Gerszten et al., 1994; Lerner et al., 1996; Nanevicz et al., 1995). Interestingly, the equivalent human PAR4 ECL2 residues distal to the CHDAL domain, P^233^LDA^236^, share sequence homology with those distal to the human PAR2 CHDVL domain (P^231^EQL^234^), both of which contain an acidic residue. Al-Ani *et al*. (1999) demonstrated that glutamic acid residues in the rat PAR2 ECL2 (PEEVL) conferred specificity to PAR2 ligands. In previous studies, a series of aspartic acids in the PAR4 extracellular loop 2 (ECL2) and amino terminal (N)-tail were proposed to be key for thrombin binding to PAR4 (Nieman, 2008; Nieman and Schmaier, 2007; Sánchez Centellas et al., 2017). Mutations of ECL2 aspartic acid residues D^224^A, D^230/235^A (double mutation), and D^224/230/235^A mutants were all found to decrease thrombin cleavage of PAR4 and activation in yeast models (Sánchez Centellas et al., 2017). Further, the activity of the PAR4 agonist peptide AYPGKF-NH_2_ was also abolished in PAR4 mutants containing ECL2 single mutant D224A, double mutant D^230/235^A, or triple mutant D^224/230/235^A aspartic acid mutations to alanine which the authors posit indicates that these aspartic acid residues may be important for both agonist peptide and tethered ligand bindings and activation of PAR4. The D^230/235^A mutation to alanine removed two acidic residues in the highly conserved …CHD^230^AL domain (PAR1 Asp^256^; PAR2 Asp^228^) as well as the homologous PLD^235^A… specificity conferring domain observed in PAR1 (Glu^260^) and PAR2 (Glu^232^) making it difficult to assess whether the loss of cleavage and activation is due to the CHD^230^AL or PLD^235^A aspartic acid.

In the present study, *in silico* docking to a homology model of PAR4 revealed 17 predicted sites of interaction with AYPGKF-NH_2_ (Ser^67^, Tyr^157^, His^229^, Asp^230^, Leu^232^, Leu^234^, Asp^235^, Ala^238^, Gln^242^, His^306^, Tyr^307^, Pro^310^, Ser^311^, Ala^314^, Gly^316^, Tyr^319^, Tyr^322^; Fig. 12 A.). Interestingly, these predicted sites reveal residues that have been shown or predicted to have a role in PAR4 activation by thrombin or agonist peptide. These sites include aspartic acid residue Asp^230^ found in the highly conserved CHDxL domain in PARs as well as Asp^235^ in the downstream region that is homologous to the specificity conferring motif found in PAR1 and PAR2 (Sánchez Centellas et al., 2017). Additionally, previous *in silico* docking revealed a role for aspartic acid and histidine in this motif in binding of AYPGKF-NH_2_ to PAR4 (Moschonas et al., 2017). We generated mutations of the most frequently predicted interaction sites (His^229^, Asp^230^, Gln^242^, His^229^/Asp^230^) to alanine which revealed that Asp^230^ is essential for productive interaction of the peptide with the receptor, as mutation to alanine abolished calcium signal potentiation as well as β-arrestin recruitment. Interestingly, this loss of activity was observed with both peptide agonist AYPGKF-NH_2_ as well as with thrombin stimulated activation of PAR4, which agrees with previous reports of the importance of these sites while also elucidating the importance of the conserved CHD^230^AL motif in PAR4 activation, consistent with the role of this domain in PAR1 and PAR2 (Al-Ani et al., 1999; Blackhart et al., 1996; Gerszten et al., 1994; Lerner et al., 1996; Nanevicz et al., 1995; Sánchez Centellas et al., 2017). Another frequently predicted residue belonging to the CHDAL domain, His^229^, when mutated to alanine did not alter the ability of either AYPGKF-NH_2_ or the tethered ligand as revealed by thrombin to activate calcium signalling or β-arrestin recruitment pathways. Further, we observed that a double mutation of both of these sites to alanine (H229A, D230A) did not result in any additive detriment to PAR4 activation; therefore, we conclude that Asp^230^ interaction with PAR4 peptide and tethered ligand agonist performs an essential role in activation of PAR4.

Finally, in the context of platelet activation we find that our modified peptides that are able to cause a greater level of calcium signalling and β-arrestin recruitment than PAR4-AP could also trigger greater platelet aggregation. In our hands, PAR4-AP was un able to elicit greater than 50% of the aggregation observed in response to 1 unit/mL of thrombin, while the peptide AYPG**R**F-NH_2_ was able to trigger aggregation comparable to that seen with thrombin. A role for PAR4-dependent calcium signalling is well established (Edelstein et al., 2014; French and Hamilton, 2016; Tourdot et al., 2018), and consistent with this, our calcium signalling-null peptide A**y**PGKF-NH_2_ failed to induce aggregation. A role for β-arrestin mediated PAR4 signalling in platelets has also been demonstrated (Khan et al., 2014; Ramachandran et al., 2017). Together these data suggest that calcium signalling is critical for initiating PAR4 mediated platelet aggregation, but the precise role hierarchy of the different PAR4 signalling pathways in platelet responses remains to be fully established.

In summary, the present studies identify key mechanisms involved in PAR4 activation and signalling. We also present and characterize a novel toolkit of PAR4 agonist peptides that could be used to study biased signalling through PAR4 in platelets as well as other physiological systems. We further show that platelet activation by PAR4 critically depends on the G*α*_q/11_ pathway, and selective targeting of this pathway might yield a useful anti-platelet agent. Overall, our work advances our understanding of agonist binding to PAR4 and will support future efforts aimed at defining signalling contribution in homeostatic signalling and the discovery of novel therapeutics targeting PAR4.

## Materials and Methods

All chemicals were purchased from Millipore-Sigma (Burlington, Massachusetts, United States), Thermo Fisher Scientific (Hampton, New Hampshire, United States), or BioShop Canada Inc. (Burlington, Ontario, Canada) unless otherwise stated. AYPGKF-NH_2_ was purchased from and synthesized by Genscript (Piscataway, New Jersey, United States). Thrombin from human plasma was obtained from Millipore-Sigma (Cat. # 605195, 5000 units; Burlington, Massachusetts, United States).

### Molecular Cloning and Constructs

The plasmid encoding PAR4-YFP was described previously (Ramachandran et al., 2017). Plasmid DNA mutations in the predicted orthosteric binding site of PAR4 were generated using the QuikChange XL Multi Site-Directed Mutagenesis Kit (Agilent Technologies, Mississauga, ON, Canada) to generate all mutants described in this study. All constructs were verified by sanger sequencing (Robarts DNA sequencing facility, University of Western Ontario). Renilla luciferase-tagged β-arrestin-1/-2 constructs were a kind gift from Michel Bouvier (Université de Montréal). PX458 vector for CRISPR/Cas9 targeting was a kind gift from Feng Zhang (Massachusetts Institute of Technology; Addgene # 48138) *<u>Site-directed mutagenesis of predicted PAR4 receptor-peptide</u> <u>interaction sites.</u>* Site-directed mutagenesis using Agilent QuikChange-XL was undertaken to generate mutations in highly predicted receptor residues to validate GalaxyPepDock model and determine key binding site residues for the native PAR4 tethered ligand and PAR4 agonist peptide AYPGKF-NH_2_. Residues were mutated by single-nucleotide substitutions replacing native residues with alanine. Successful mutations were confirmed by DNA sequencing (London Regional Genomics Centre, London, ON).

### Cell Lines and Culture Conditions

All media and cell culture reagents were purchased from Thermo Fisher Scientific (Hampton, New Hampshire, United States). HEK-293 cells (ATCC; Manassas, Virginia, United States) and CRISPR/Cas9 HEK-293 β-arrestin double knockout (β-arr knockout) cell lines were maintained in Dulbecco’s modified Eagle’s medium supplemented with 10% fetal bovine serum, 1% sodium pyruvate, and 1% streptomycin penicillin. Cells stably transfected with PAR4-YFP vector was routinely cultured in the above media supplemented with 600 µg/mL G418 sulfate (Geneticin; ThermoFisher Scientific Waltham, Massachusetts, United States). Since trypsin activates PAR4, cells were routinely sub-cultured using enzyme-free isotonic phosphate-buffered saline (PBS), pH 7.4, containing 1 mM EDTA followed by centrifugation to remove PBS-EDTA and plated in appropriate culture plates for further experimentation. Cells were transiently transfected with appropriate vectors using a modified calcium phosphate transfection (Nuclease free water, 2.5M CaCl_2_, and 2x HEPES Buffered Saline). Transient transfections were rinsed with phosphate buffered saline (PBS) 24 hours post-transfection and replaced with growth media. Transiently transfected cells were assayed or imaged at 48 hours post-transfection. *<u>Generation of β-</u> <u>arrestin-1/-2 double knockout HEK-293 cells using CRISPR/Cas9</u>*. A β-arrestin-1/-2 double knockout HEK-293 cells cell line (β-arr. knockout) was generated using CRISPR/Cas9 mediated gene targeting. Guides sequences targeted to β-arrestin-1 or β-arrestin-2 were designed using the design tool at http://crispr.mit.edu. Gene targeting guides (β-arrestin 1; TGTGGACCACATCGACCTCG, β-arrestin 2; GCGGGACTTCGTAGATCACC, were ligated into the PX458 vector (Kind gift from Dr. Feng Zhang, MIT, Addgene plasmid #48138), verified by direct sequencing and transfected in HEK-293 cells via calcium phosphate transfection (Ferguson and Caron, 2004; Graham and van der Eb, 1973). GFP expressing cells were single-cell selected by flow cytometry (FACSAria III; London Regional Flow Cytometry Facility) into wells of a 96-well plate. Clonal cell populations were expanded and screened by western blot to identify successful knockout (anti-β-arrestin-1/-2 antibody, rabbit mAb D24H9; Cell Signalling Technologies, Danvers, Massachusetts, United States; anti-β-actin antibody, mouse mAb 8H10D10; Cell Signalling Technologies; Supplemental Figure 3).

### General Methods for Peptide Synthesis

All reagents were purchased from Sigma-Aldrich, ChemImpex, or Thermo Fisher Scientific and used without further purification. Peptides were synthesized using standard Fmoc-solid phase peptide synthesis (SPPS) on Rink amide MBHA resin (256 mg, 0.1 mmol, 0.39 mmol/g) with a Biotage® Syrowave^TM^ automated peptide synthesizer (0.4 mmol of HCTU, 0.4 mmol of Fmoc-amino acids, 0.6 mmol of DIPEA, 1 h coupling). Some peptides also involved additional manual synthesis (see procedures below). Following automated and manual synthesis, peptides were cleaved off the resin by being shaken in a cleavage cocktail solution of 95% TFA, 2.5% TIPS, and 2.5% H_2_O (4 mL, 5 h), precipitated in ice cold *tert*-butyl methyl ether, lyophilized, purified by preparative RP-HPLC, and further lyophilized to obtain a dry powder. Purity was assessed by analytical RP-HPLC and characterized by mass spectrometry (MS). The analytical RP-HPLC was performed on a system consisting of an analytical Agilent Zorbax SB-C8 column (4.6 x 150 mm, 5 µm), Waters 600 controller, Waters in-line degasser, and Waters Masslynx software (version 4.1). Two mobile phases were used; eluent A (0.1% TFA in acetonitrile) and eluent B (0.1% TFA in MilliQ water). The flow rate was set at 1.5 mLmin^-1^ over 10 minutes with an additional 5-minute wash (95% solvent A in solvent B). A Waters 2998 Photodiode array detector (200-800 nm) and an ESI-MS (Waters Quattro Micro API mass spectrometer) were used to monitor the column eluate. The preparative RP-HPLC used the same system, eluents, and detection method as mentioned above for the analytical RP-HPLC, except that a preparative Agilent Zorbax SB-C8 column (21.2 x 150 mm, 5 µm) at a flow rate of 20 mLmin^-1^ was used. The mass spectra for all peptides were determined in positive mode using an ESI ion source on either a Bruker micrOToF II mass spectrometer or a Xevo QToF mass spectrometer (Table S1). All peptides had purity >95% as determined by analytical HPLC (see supplemental information, Figure S1).

### Synthesis of Ac-AYPGKF-NH_2_

The hexapeptide (AYPGKF) was synthesized using the standard automated Fmoc-SPPS discussed in the general methods for peptide synthesis. While still on resin, with all of the side chains protected, and with the N-terminal Fmoc-deprotection completed, the N-terminus of this peptide was acetylated by being shaken in a solution of acetic anhydride and DMF (v/v 1:4, 5 mL, 30 min). Following this, the peptide was cleaved, purified, and analyzed as mentioned in the general methods for peptide synthesis.

### Synthesis of Peptides Involving N-methylation of the Peptide Backbone

Peptides [(*N*-Me-A)-YPGKF-NH_2_, (*N*-Me-S)-YPGKF-NH2, A-(*N*-Me-Y)-PGKF-NH2, AYPG-(*N*_α_-Me-K)-F-NH2, and AYPGK-(*N*-Me-F)-NH_2_] were synthesized using abovementioned automated Fmoc-SPPS up to the desired amino acid that was *N*-methylated (e.g. KF-resin for AYPG-(*N*_α_-Me-K)-F-NH_2_). Following Fmoc-deprotection of the resin-bound N-terminal amino acid (e.g. lysine for AYPG-(*N*_α_-Me-K)-F-NH_2_), site selective *N*-methylation of the N-terminal amino acid was completed as follows: First, protection of the primary amine was completed to favour monomethylation by adding 2-nitrobenzenesulfonyl chloride (110.8 mg, 0.5 mmol) and 2,4,6-trimethylpyridine (132 µL, 1.0 mmol) in NMP (4 mL) to the on-resin peptide (0.1 mmol) and shaken (15 min). Following completion of the protection reaction, the resin was washed thrice with NMP (3X 5 mL) and dry THF (3X 5 mL). Second, *N*-methylation was completed through Mitsunobu conditions by adding triphenylphosphine (131.2 mg, 0.5 mmol), diisopropyl azodicarboxylate (98 µL, 0.5 mmol), and methanol (41 µL, 1.0 mmol) in dry THF (4 mL) to the on-resin peptide and shaken under N_2_ (10 min). Following completion of the *N*-methylation reaction, the resin was washed thrice with dry THF (3X 5mL) and NMP (3X 5 mL). Third, the methylated N-terminal amine was then deprotected by adding 2-mercaptoethanol (70 µL, 1.0 mmol) and 1,8-diazabicyclo(5.4.0)undec-7-ene (75 µL, 0.5 mmol) in NMP (4 mL) to the on-resin peptide and shaken (5 min). The solution turned bright yellow as expected. The solution was removed and then this third step was repeated, followed by being washed five times with NMP (5X 5 mL) and thrice with DMF (3X 5 mL). Addition of the next sequential amino acid (except for inapplicable peptides i.e. (*N*-Me-A)-YPGKF-NH_2_ and (*N*-Me-S)-YPGKF-NH_2_) was added manually [0.3 mmol of HATU, 0.3 mmol of Fmoc-amino acid (e.g. Fmoc-Gly-OH for AYPG-(*N*_α_-Me-K)-F-NH_2_), 0.6 mmol of DIPEA, 4 mL of DMF, 3 h coupling]. Following completion of the sequential amino acid reaction, the resin was washed 5 times with DMF (5X 5mL) and then the next sequential amino acid was added (on applicable peptides i.e AYPG-(*N*α-Me-K)-F-NH_2_ and AYPGK-(*N*-Me-F)-NH_2_) using the automated procedure mentioned in the general methods for peptide synthesis. Following completion of the synthesis, the peptide was cleaved, purified, and analyzed as mentioned in the general methods for peptide synthesis. The amino acid *N*-methylglycine, commonly known as sarcosine, was obtained from a commercial source, and therefore, *N*-methylglycine containing peptides (i.e. Sar-YPGKF-NH_2_ and AYP-Sar-KF-NH_2_) were synthesized solely by the standard automated Fmoc-SPPS discussed in the general methods for peptide synthesis.

### Synthesis of AYPG-(N_ε_-Me-K)-F-NH_2_

The hexapeptide (AYPGKF) was synthesized using the standard automated Fmoc-SPPS discussed in the general methods for peptide synthesis, except that Fmoc-Lys(Alloc)-OH was used in place of standard Fmoc-Lys(Boc)-OH. While still on resin, with all of the side chains protected, and with the N-terminal Fmoc-deprotection completed, the N-terminus of this peptide was Boc protected by adding Boc-ON (98.5 mg, 0.4 mmol) and DIPEA (70 µL, 0.4 mmol) in DMF (4 mL) and shaken (4 hours). The solution was removed and this Boc protection was repeated overnight (20 hours) to ensure reaction completion. The resin was washed thrice with DMF (3X 5 mL) and dry DCM (3X 5 mL) and was placed under N_2_ for site selective Alloc deprotection of the on-resin lysine side chain. Phenylsilane (296 μL, 2.4 mmol) in dry DCM (2 mL) was added to the resin. Tetrakis(triphenylphosphine) palladium(0) (23.1 mg, 20 µmol) was dissolved in dry DCM (1 mL) and added to the resin. The peptide column was flushed with N_2_ (2 min) followed by being shaken (5 min). The resin was washed thrice with dry DCM (3X 5 mL). The procedure was repeated and shaken again (30 min). The resin was washed four times with DCM (4X 5 mL) and DMF (4X 5 mL). Following this N-terminal Boc protection and subsequent side chain Alloc deprotection, *N*-methylation of the on-resin lysine side chain was completed through the site selective *N*-methylation procedure mentioned above. Following completion of the synthesis, the peptide was cleaved, purified, and analyzed as discussed in the general methods for peptide synthesis.

### SPPS Reaction Monitoring

Three methods were used to monitor SPPS reactions. Two methods (the Kaiser Test and the small-scale resin cleavage) were used to monitor general peptide synthesis. Three methods (the Kaiser Test, the small-scale resin cleavage, and the Chloranil Test) were used to monitor synthesis of peptides involving *N*-methylation. For the Kaiser Test, several resin beads were placed in a test tube followed by the addition of 42.5 mM phenol in ethanol (50 µL), 20 µM potassium cyanide in pyridine (50 µL), and 280.7 mM ninhydrin in ethanol (50 µL). The mixture was then heated (100 °C, 5 min); where a positive test indicates the presence of a free primary amine. For the small-scale resin cleavage method, several beads and cleavage cocktail (500 µL) are shaken, worked up as per the full cleavage of the resin procedure described above, and the desired peptide was confirmed through HPLC-MS. For the Chloranil Test, several resin beads were placed in a test tube followed by the addition of 357.8 mM acetaldehyde in DMF (50 µL) and 81.3 mM *p*-chloranil in DMF (50 µL) and allowed to stand at room temperature (5 min); where a positive test indicates the presence of a free secondary amine.

### Calcium Signalling Assay

HEK-293 cells or HEK-293 cells stably expressing PAR4-YFP were loaded with calcium sensitive dye (Fluo-4 NW, F36206, Life Technologies, Thermo Fisher) and aliquoted into cuvettes with Hanks buffered salt solution (HBSS containing Ca^2+^ and Mg^2+^). Samples were excited at 480nm and emission recorded at 530nm. Baseline fluorescence from PAR4-YFP excitation remains constant throughout experiment. Following baseline recording, agonist induced changes in fluorescence was recorded. Change in fluorescence from baseline to maximum response at a given agonist concentration (delta) were expressed as a percentage of maximum cellular response elicited by calcium ionophore (A23187, Sigma-Aldrich). *<u>Calcium Signalling Assay with PAR4 orthosteric site-directed</u> <u>mutants</u>.* HEK-293 cells were transiently transfected with 2µg of wild-type YFP tagged PAR4 or PAR4 mutant receptors (described above) using the calcium phosphate transfection protocol as described above. Calcium signalling was investigated as described above for the wild type receptor.

### β-arrestin Recruitment - Bioluminescence Resonance Energy Transfer (BRET)

HEK-293 cells were co-transfected with PAR4-YFP (2µg) or mutant PAR4-YFP and β-arrestin-1- or -2-renilla luciferase (rluc, 0.2µg) using modified calcium phosphate transfection. Cells were re-plated (24 hours post-transfection) into white 96-well plates (Corning). At 48-hours post-transfection, agonist-stimulated β-arrestin recruitment to PAR4 was detected by measuring BRET following 20 minutes of agonist stimulation. Coelenterazine (5µM; Nanolight Technology, Pinetop, AZ) was added ten minutes prior to BRET collection using a Berthold Mithras LB 940 plate reader.

### Testing Peptide library specificity for PAR4

Previous studies have pointed to the possibility of PAR agonist peptides cross activating other members of this family. For example, the PAR1 tethered ligand peptide SFLLRN is able to activate PAR2 and substitution of a phenylalanine residue in certain PAR4 agonist peptides confers PAR1 activity (Boitano et al., 2011; Faruqi et al., 2000; Lerner et al., 1996). Activating peptides typically require an aromatic sidechain in position 2 (F in PAR1) and we wondered if substitution of tyrosine for phenylalanine may decrease selectivity at PAR4 and increase crosstalk with PAR1. We examined whether substitutions made to AYPGKF-NH_2_ affected their selectivity using a calcium signalling assay in HEK-293 cells that endogenously express PAR1 and PAR2. None of the peptides tested elicited a signalling greater than 5% of the calcium ionophore A23187 showing that they retain selectivity towards PAR4 (Supp. Fig. 4). Additionally, we investigated specificity using BRET for β-arrestin-2 recruitment to cells expressing PAR4-YFP or PAR2-YFP. None of the peptides were able to induce β-arrestin recruitment in PAR2-YFP expressing HEK-293 cells (NET BRET of 0.05 and over) with the exception of AYPGK-1Nal-NH_2_ (Supp. Fig. 5).

### PAR4-stimulated Mitogen-activated Protein Kinase (MAPK) signalling

*<u>HEK-293 and</u> <u>CRISPR/Cas9 β-arrestin-1/-2 double knockout HEK-293.</u>* Agonist-stimulated mitogen-activated protein kinase (MAPK) signalling in CRISPR/Cas9 HEK-293 β-arrestin double knockout cells (β-arr. knockout) and wild-type HEK-293 cells transfected with PAR4-YFP were analysed by western blot. Cells were placed in serum free media (serum-free DMEM, Gibco) for 3 hours before beginning experiment. Cells were then stimulated with 30μM AYPGKF-NH_2_ for varying time periods (0, 2, 5, 10, 20, 30, 60, 90 minutes) at 37 °C. Treated cells were placed on ice following stimulating period. Total protein was extracted adding kinase lysis buffer with phosphatase and protease inhibitors (20 mM Tris-HCl, pH 7.5, 100 mM NaCl, 10 mM MgCl_2_, 1 mM EDTA, 1 mM EGTA, 0.5% NP-40, 2.5 mM sodium pyrophosphate, 1 mM b-glycerophosphate, 1 mM Na_3_VO_4_, 25 mM NaF, 1 mg/mL leupeptin, 1 mg/m aprotinin, 1 mM phenylmethylsulfonyl fluoride, and 1mM dithiothreitol) for 20 minutes and cell membranes were cleared by centrifugation (13,300g for 3 minutes). Protein concentrations were measured using the Pierce BCA Protein Assay (Thermo Fisher Scientific). Protein samples were heat-denatured at 98°C for 8 minutes in denaturing Laemmli buffer (containing 2-mercaptoethanol) and resolved on 4–12% gradient Invitrogen Bolt Bis-Tris Plus gels (Thermo Fisher Scientific). The resolved proteins were transferred via wet transfer in buffer (1X Tris-glycine, 20% methanol) to a polyvinylidene difluoride membrane and blocked in TBST buffer (TBS with 0.1% (v/v) Tween-20) supplemented with 1% ECL Advance Blocking Agent (GE Healthcare, Waukesha, WI) for 40 minutes at room temperature. p44/42 (Thr202/Tyr204) phosphorylation was detected with specific antibodies (Cell Signalling Technology, Danvers, MA; diluted 1:1000 in TBST) overnight at 4°C. phospho-p44/42 immunoreactivity was detected using the horseradish peroxidase-conjugated anti-rabbit secondary antibody (Cell Signalling Technology; 1:10,000 in TBST for 1 hour). After washing the membrane with TBST (3 x 10-minute washes), the peroxidase activity was detected with the chemiluminescence reagent ECL Advance (GE Healthcare) on iBright image Station (Invitrogen, Burlington, ON). Membranes were then stripped with blot stripping buffer (Thermo Fisher Scientific) at room temperature and blocked in TBST with 1% ECL Advance Blocking Agent before incubation with the p44/42 (1:1000 in TBST with 1% ECL Advance Blocking Agent) overnight at 4°C. Following incubation with primary antibody, blots were rinsed in TBST and incubated with appropriate horseradish peroxidase -conjugated anti-rabbit or anti-mouse secondary antibody (1:10,000 in TBST for 1 hour). Membranes were washed and imaged as previously described. Band intensities representing activated MAPK were quantified using FIJI (Schindelin et al., 2012). Phospho-kinase levels were normalized for differences in protein loading by expressing the data as a ratio of the corresponding total-p44/42 signal. Data are expressed as a fold increase over unstimulated baseline control. *<u>PAR4-</u> <u>stimulated MAPK signalling in HEK-293 +/-YM254890.</u>* Agonist-stimulated MAPK signalling in wild-type HEK-293 cells transfected with PAR4-YFP were analysed by western blot. Cells were placed in serum free media (serum-free DMEM, Gibco) for 3 hours before beginning experiment. Prior to agonist stimulation cells were treated with either control (0.001% DMSO) or YM254890 (100nM) for 20 minutes. Cells were then stimulated with 30μM AYPGKF-NH_2_ for varying time periods (0, 2, 5, 10, 20, 30, 60, 90 minutes) at 37 degrees Celsius. Cells were lysed, protein quantified, and samples run and imaged as previously described. *<u>MAPK</u> <u>signalling in PAR4-YFP stable cells with peptide library</u>*. Agonist-stimulated mitogen-activated protein kinase (MAPK) signalling in HEK-293 cells stably expressing PAR4-YFP were analysed by western blot following stimulation with PAR4 peptide agonists. Cells were placed in serum free media (serum-free DMEM, Gibco) for 3 hours before beginning experiment. Cells were stimulated with either control (no agonist), AYPGKF-NH_2_ or agonist peptides described at 100μM concentration. Cells were stimulated for 10 minutes with agonist and lysed, quantified, and blot as previously described.

### Preparation of washed platelets and light-transmission aggregometry

Blood was drawn from the abdominal aorta or heart of Sprague-Dawley rats into trisodium citrate (3.4% w.v). The blood was centrifuged at 200g for 15 minutes at 22 degrees Celsius to isolate platelet-rich plasma (PRP). PRP was transferred to a 14 mL polypropylene round-bottom tube (Falcon) and centrifuged at 1800 rotations per minute (rpm.; 594g) for 10 minutes at 22°C to pellet platelets and obtain platelet-poor plasma (PPP). PPP was decanted and used for 100% light transmission control. To obtain washed platelet sample, platelets were washed twice in Tyrode’s buffer (12 mM NaHCO_3_, 127 mM NaCl, 5 mM KCl, 0.5 mM NaH_2_PO_4_, 1 mM MgCl_2_, 5 mM glucose, and 10 mM HEPES; Ramachandran et al., 2017) and resuspended to a volume equivalent to that of the initial blood draw. Washed platelet sample was equilibrated for one hour at ambient temperature following the addition of 1M CaCl_2_ (1 µl/mL). 400 µl of this washed platelet suspension was used for light transmission aggregometry on a platelet aggregometer (Model 700; Chrono-Log Corp, Havertown, PA).

### Structural homology modelling of PAR4

Residues resolved in the PAR2 crystal (5NDD.pdb) were used to generate a homology model of wild-type human PAR2. Homology model of wild-type human PAR2 was then aligned with the wild-type human PAR4 amino acid sequence [National Center for Biotechnology Information (NCBI) accession number NM_003950) using Clustal Omega (Madeira et al., 2019; Sievers et al., 2011). Only residues resolved in the crystal structure of PAR2 were used as template coordinates for PAR4 homology modelling. Twenty human PAR4 homology models were generated using the human PAR2 crystal structure coordinates (5NDD.pdb; Cheng et al., 2017) as a template in MODELLER (version 9.16, Webb and Sali, 2017). The model with the lowest DOPE score was taken as the best structure and visualized using PyMOL (version 1.7.4, Schrödinger, LLC). Electrostatic surface potential was calculated after assignment of partial charges using PDB2PQR and using the Adaptive Poisson-Boltzmann Solver plugin in PyMOL (Dolinsky et al., 2004).

### In silico docking of AYPGKF-NH_2_ to the PAR4 homology model

*in silico* docking of PAR4 agonist peptide, AYPGKF-NH_2_, to PAR4 homology model was undertaken using GalaxyPepDock (Lee et al., 2015). The lowest energy PAR4 homology model PDB file was loaded into GalaxyPepDock along with a text file of the PAR4 agonist peptide sequence which GalaxyPepDock uses to predict a peptide structure for docking analyses (Ala-Tyr-Pro-Gly-Lys-Phe; amidation is not included in the structure prediction). Ten models of PAR4-AP docking were generated. Frequency of receptor-peptide residue interactions, taken as atom-atom distances of less than 3 Angstroms, was quantified manually to determine highly recurrent interactions.

### Data analysis and statistical tests

Data are shown as mean ± SEM. Calcium signalling and β-arrestin-1/-2 recruitment concentration effect curves were calculated using nonlinear regression curve fitting (three parameter; Prism7). EC_50_ values are reported (Table 1) where signal has saturated. Where signalling is not saturated EC_50_ is reported for calcium as not determined (*n.d.*). The F-statistic calculated to compare EC_50_ values was calculated using the extra sum-of-squares analysis (Kim et al., 2019). For table 1, values comparing percentage of signalling or recruitment at 100 µM to values obtained from the parental PAR4-AP, AYPGKF-NH_2_, are shown. Significance was determined for comparisons with 100 µM AYPGKF-NH_2_ and p44/42 phosphorylation by one-way analysis of variance (ANOVA). Western blot protein densitometry was quantified using FIJI. Transformation and analysis were conducted on raw data and significance is indicated by ‘*’ (p < 0.05).

## Acknowledgements and Funding

These studies were funded by a Canadian Institutes of Health Research (CIHR) grant (RR) and grants from the Natural Sciences and Engineering Research Council of Canada (LL and PS). PT is the recipient of a doctoral QEII Graduate Scholarships in Science and Technology. JCL is the recipient of a Canada Graduate Scholarship and an Ontario Graduate Scholarship.

## Supplemental tables and figures

**Figure S1:**
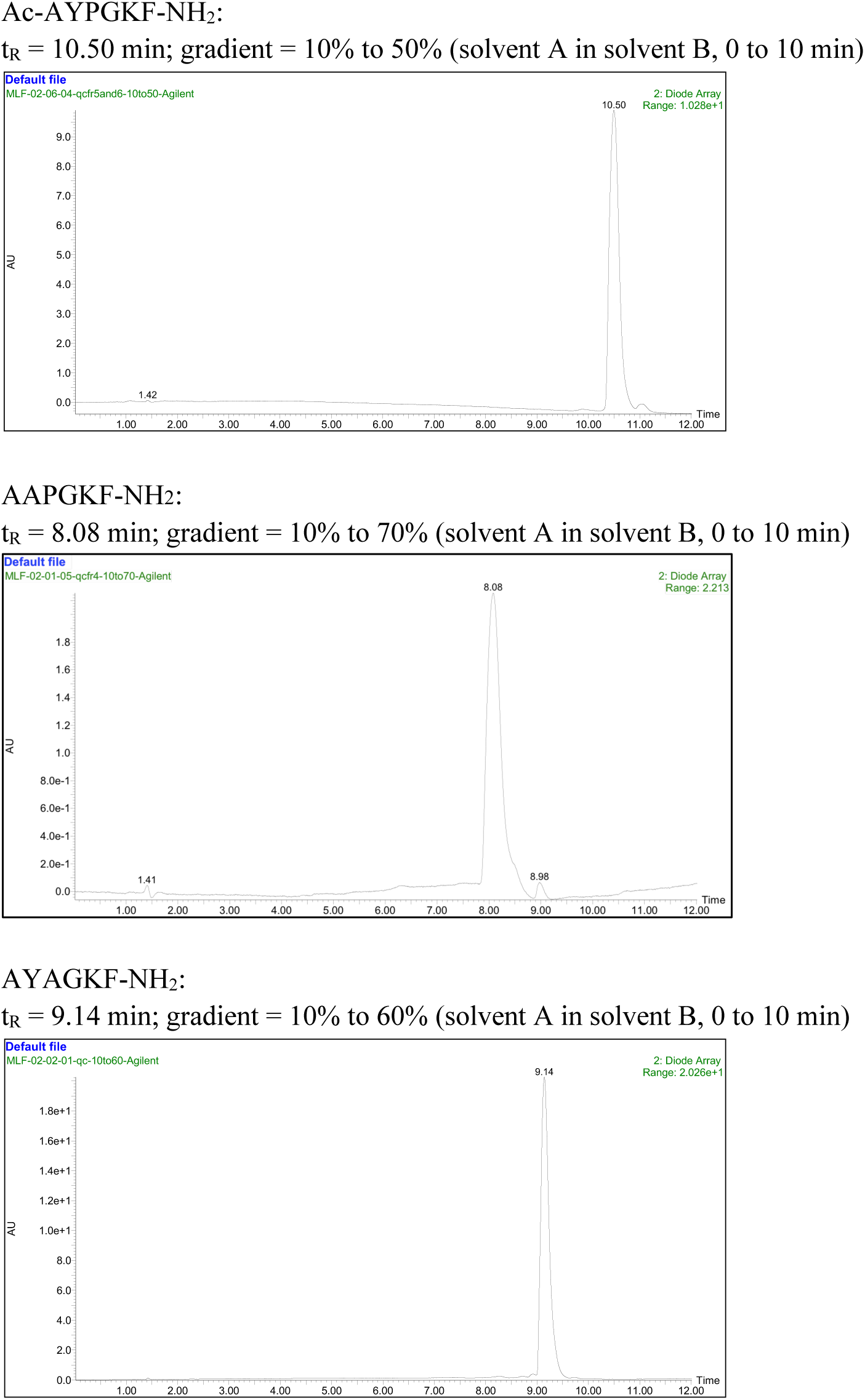
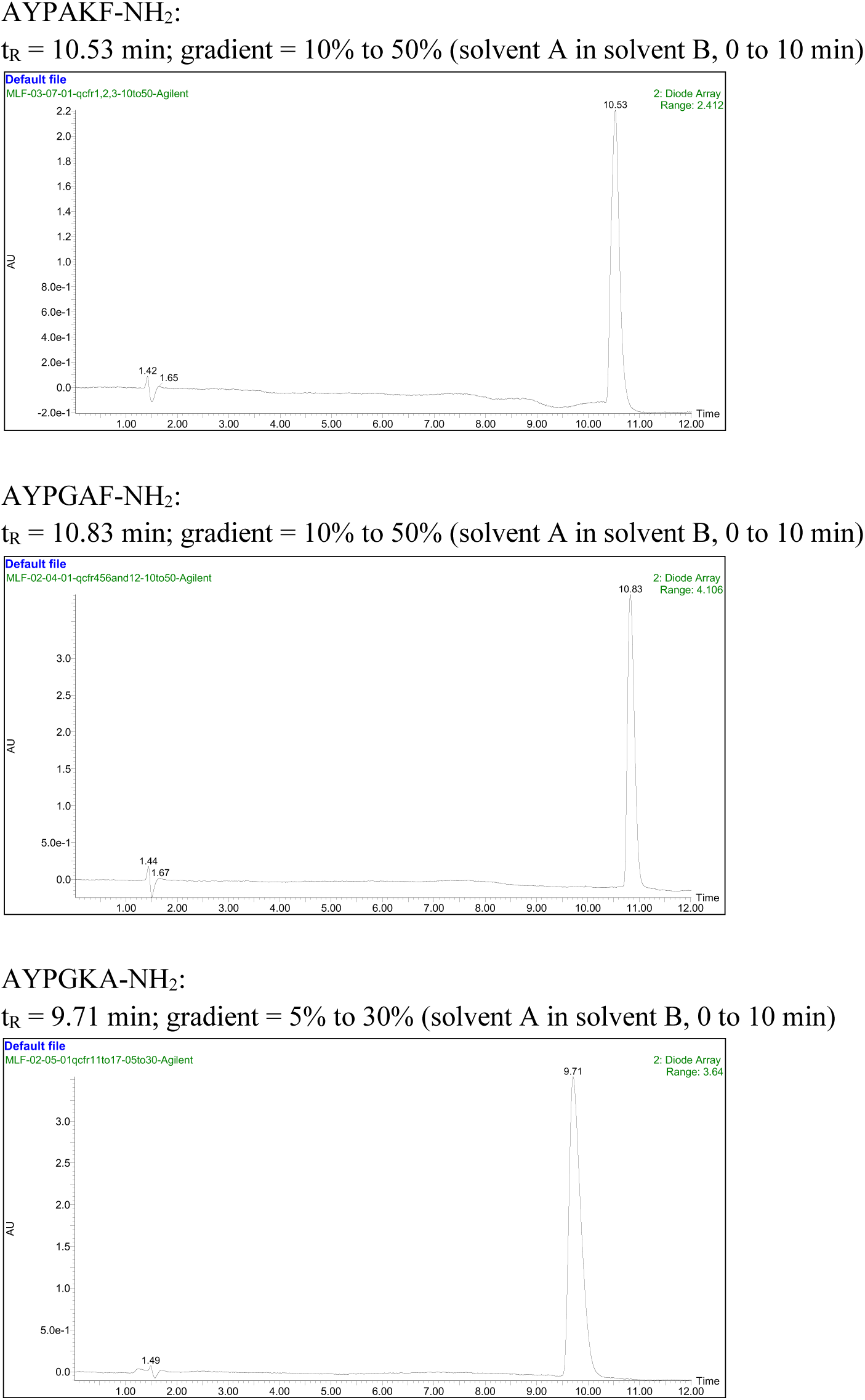
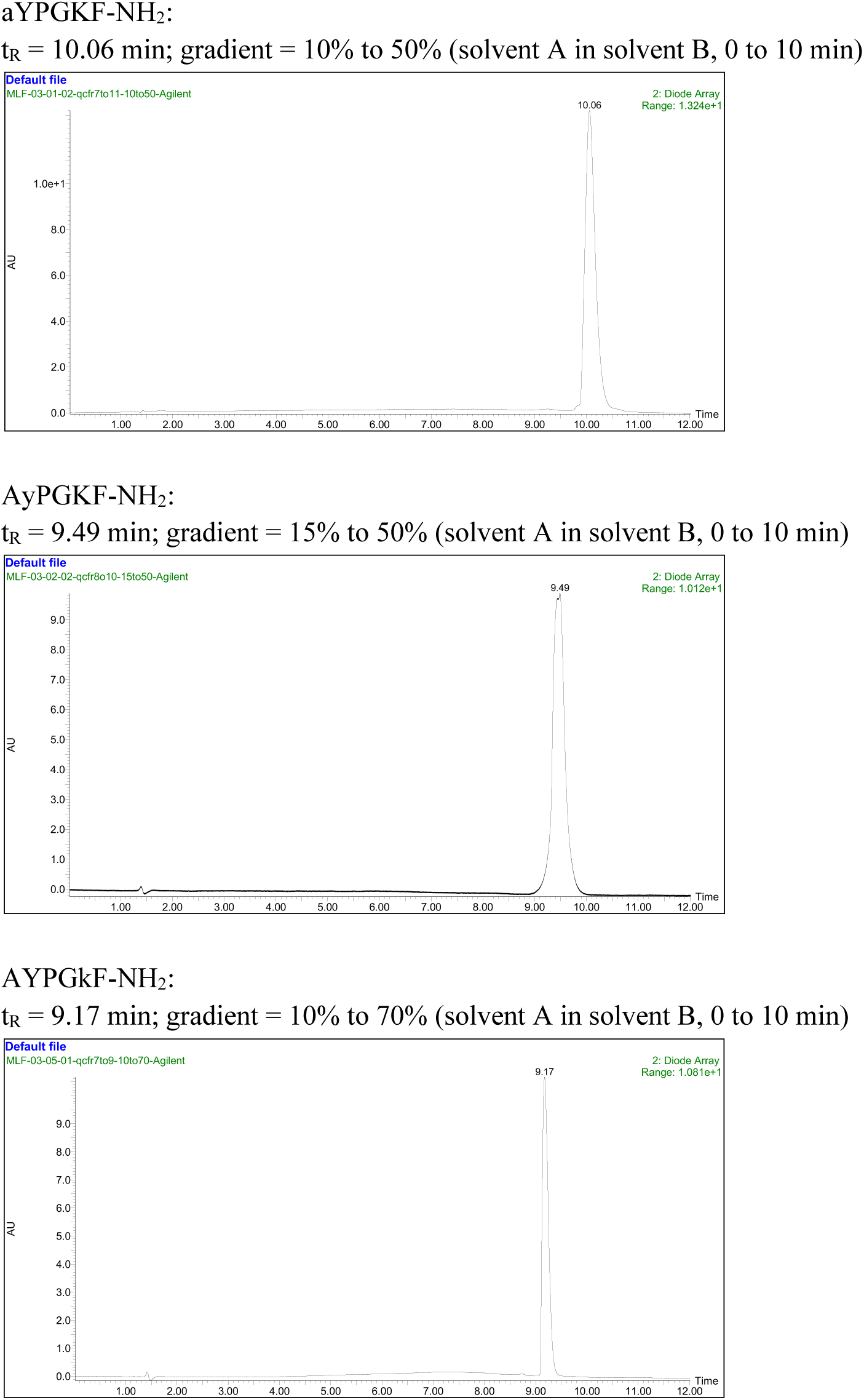
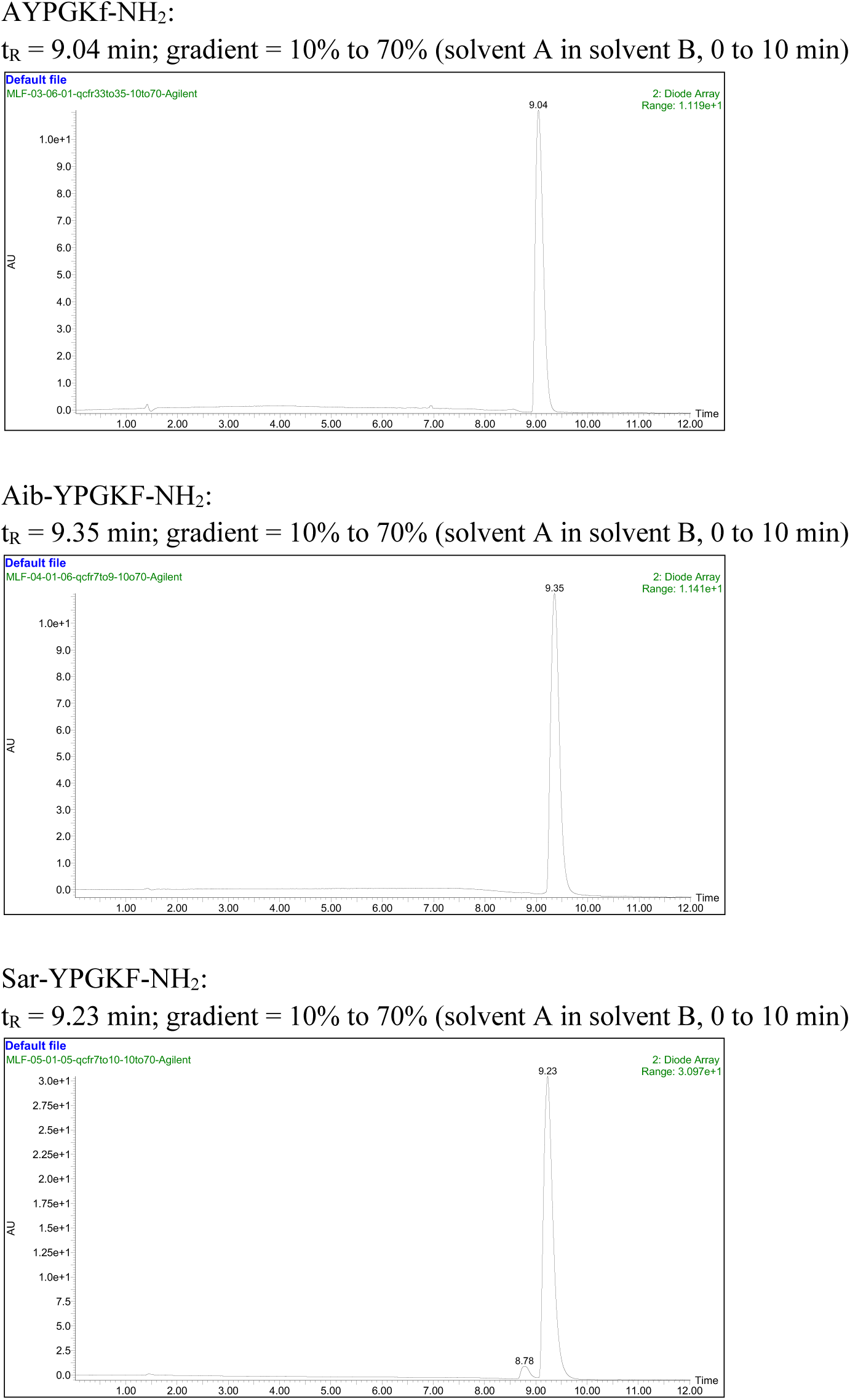
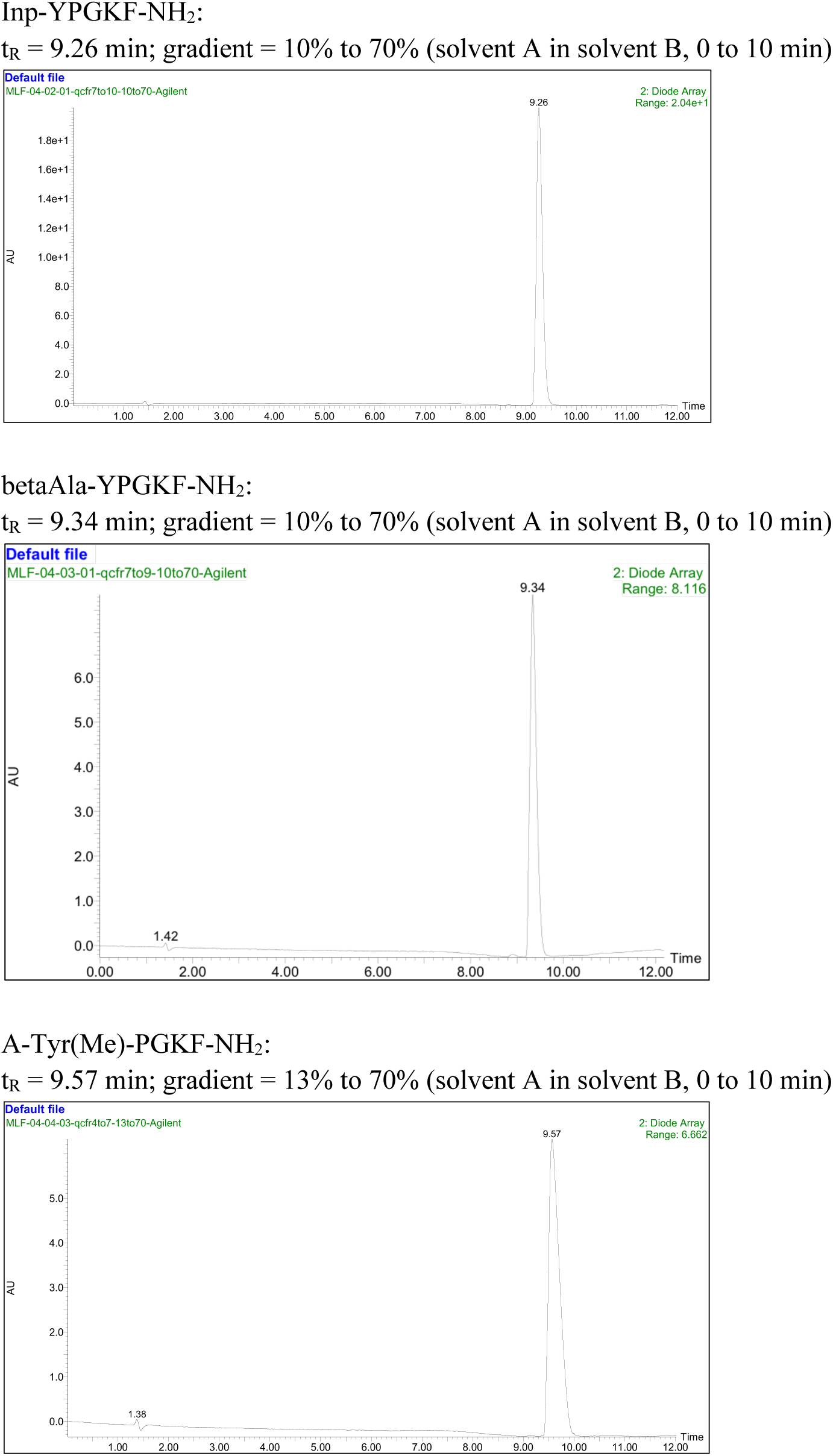
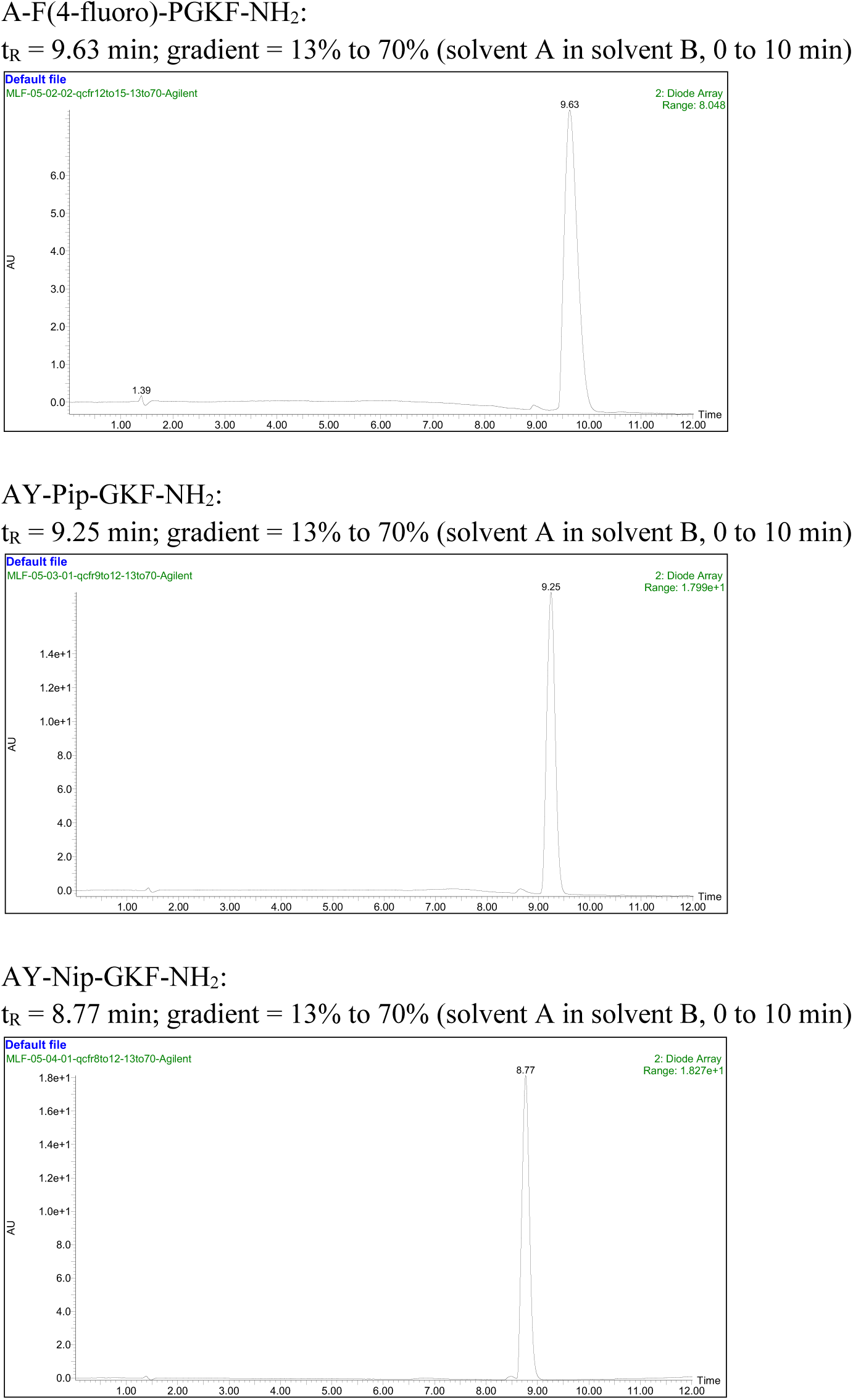
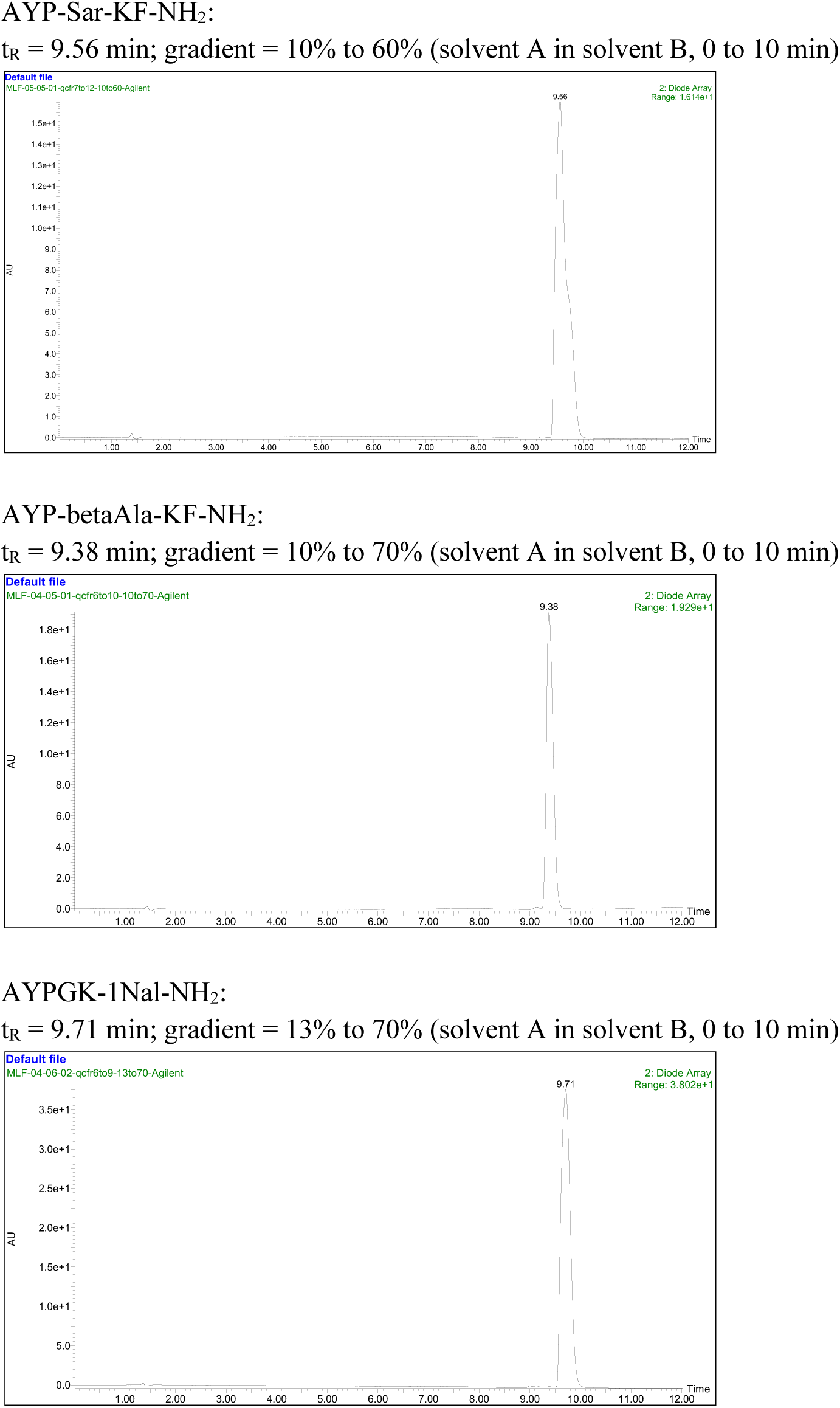
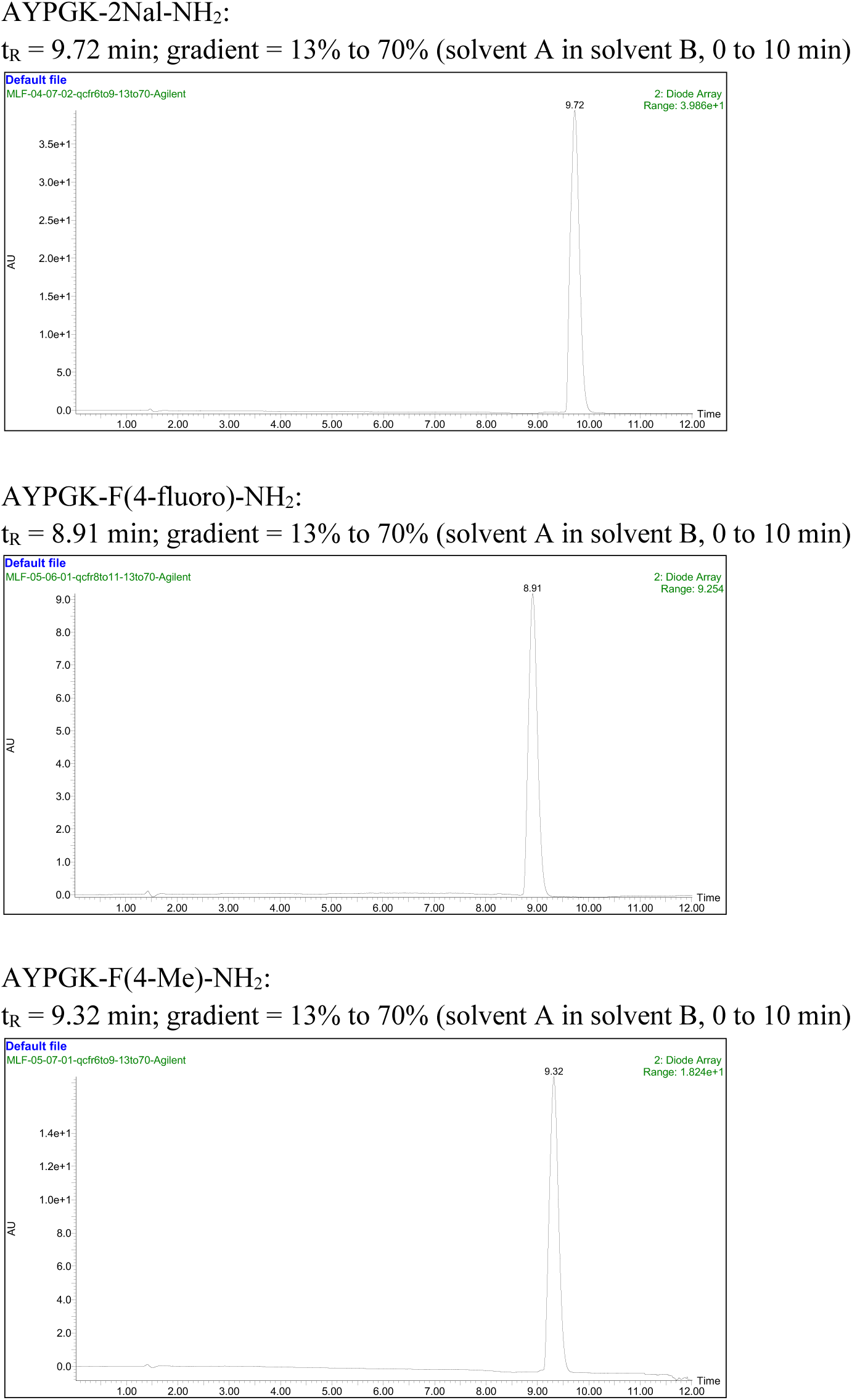
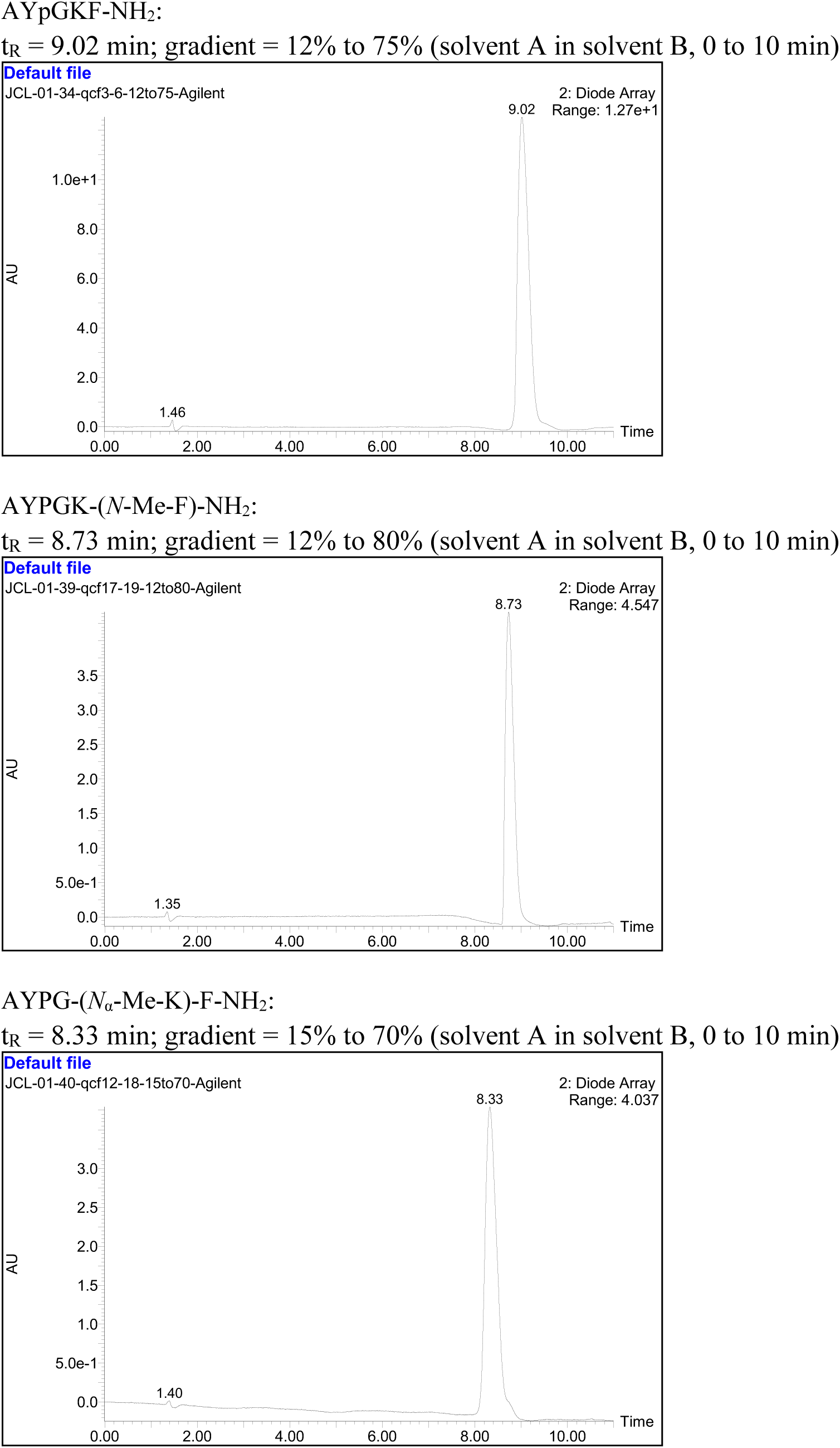
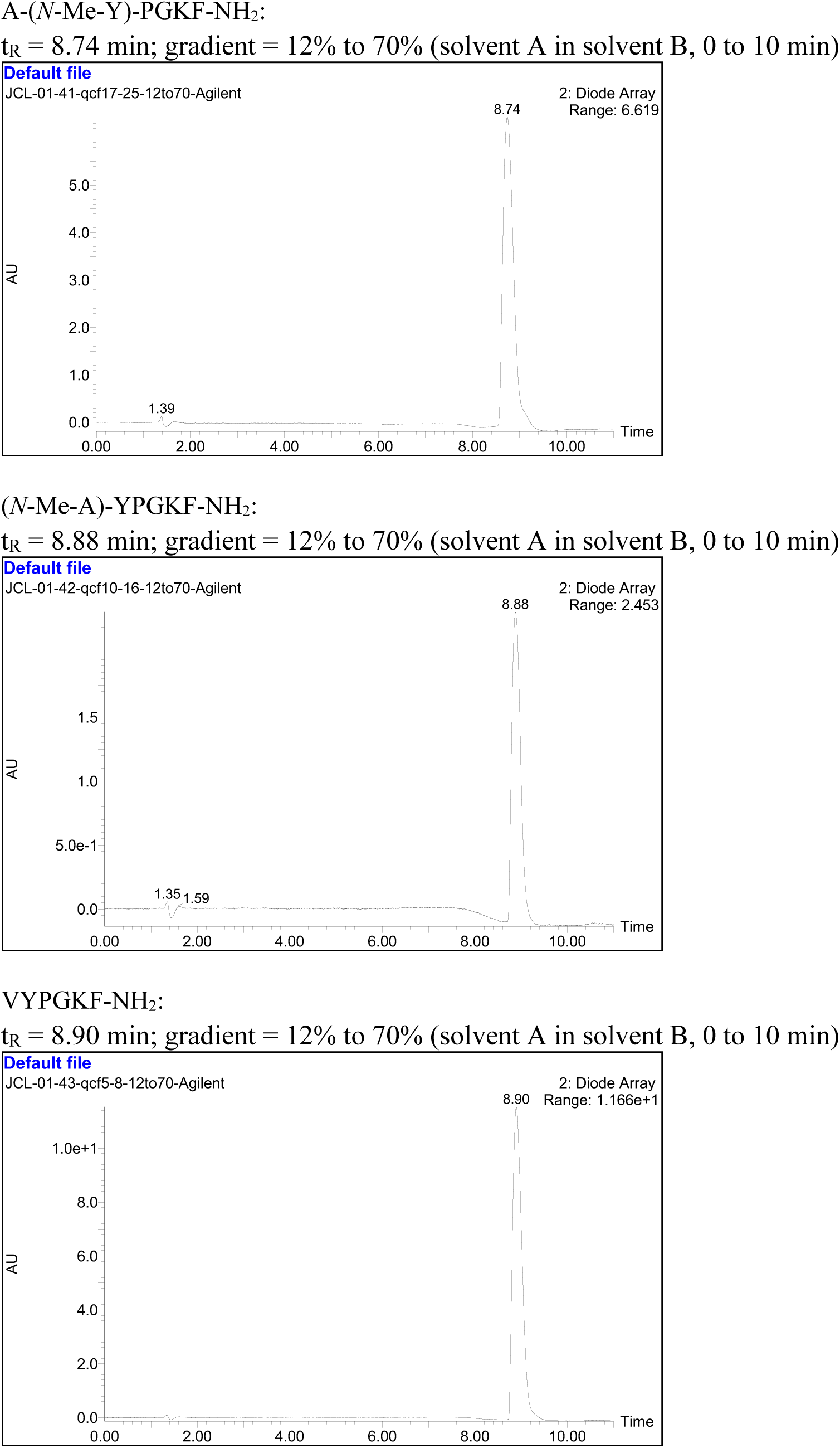
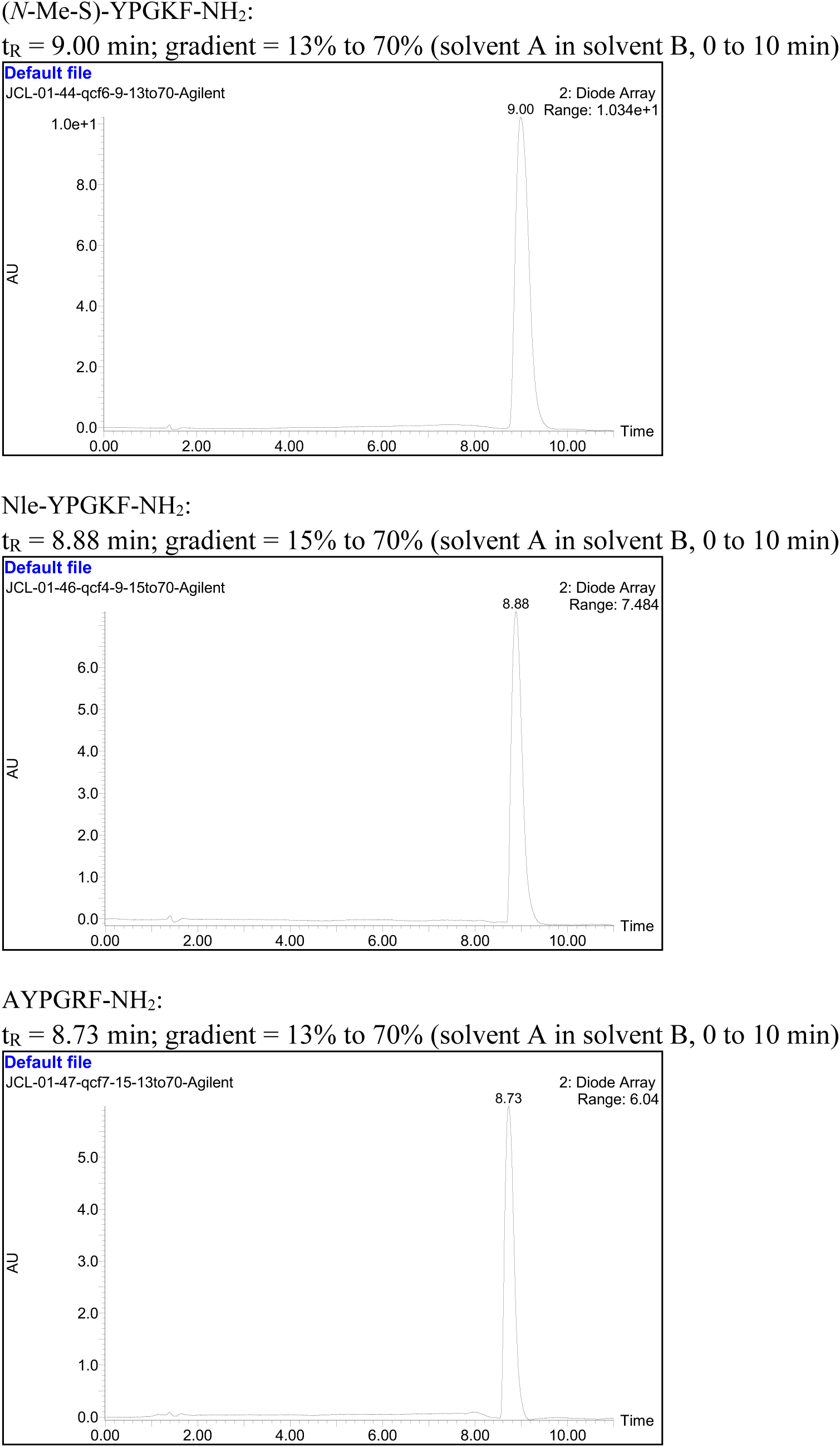
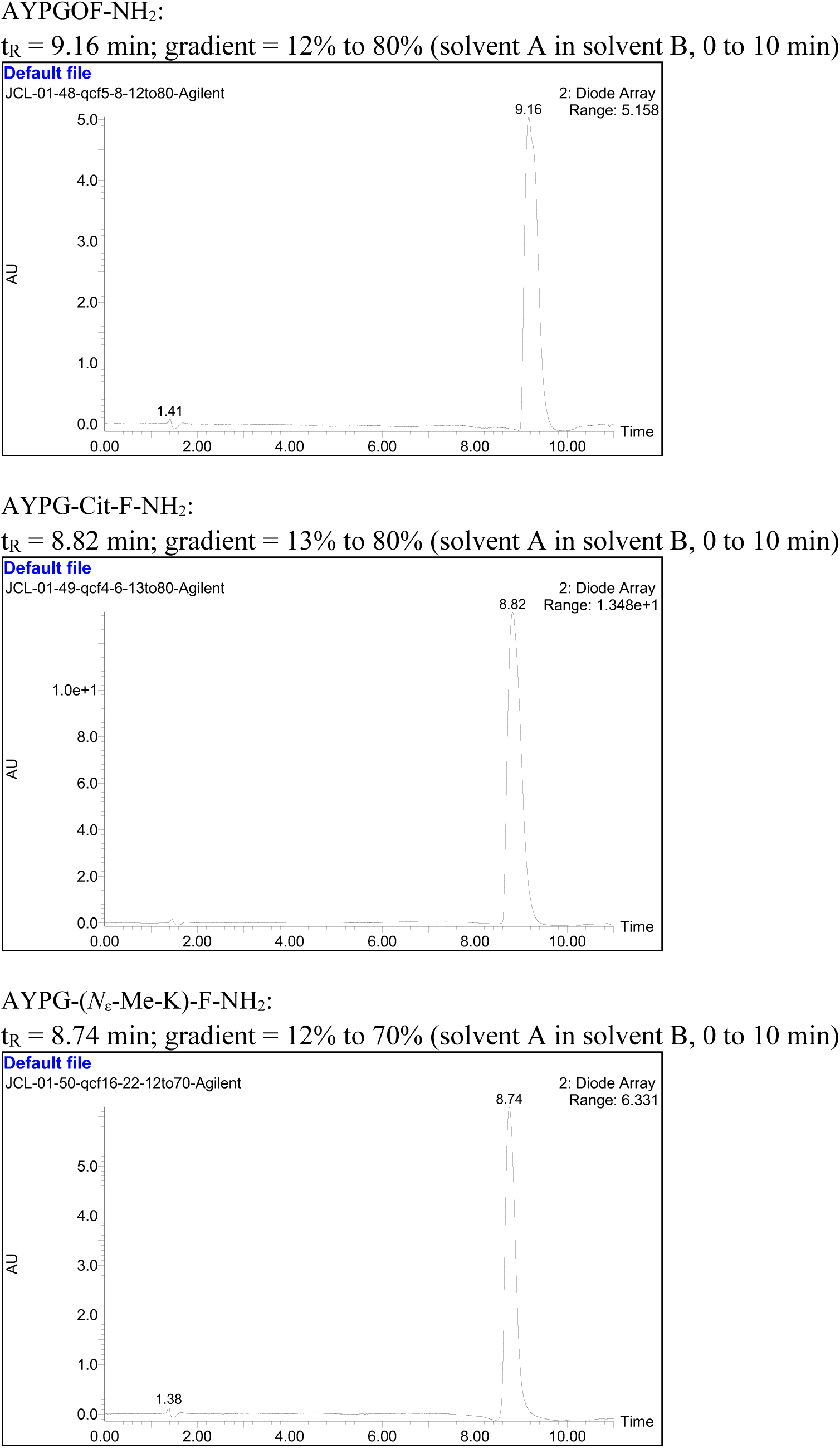
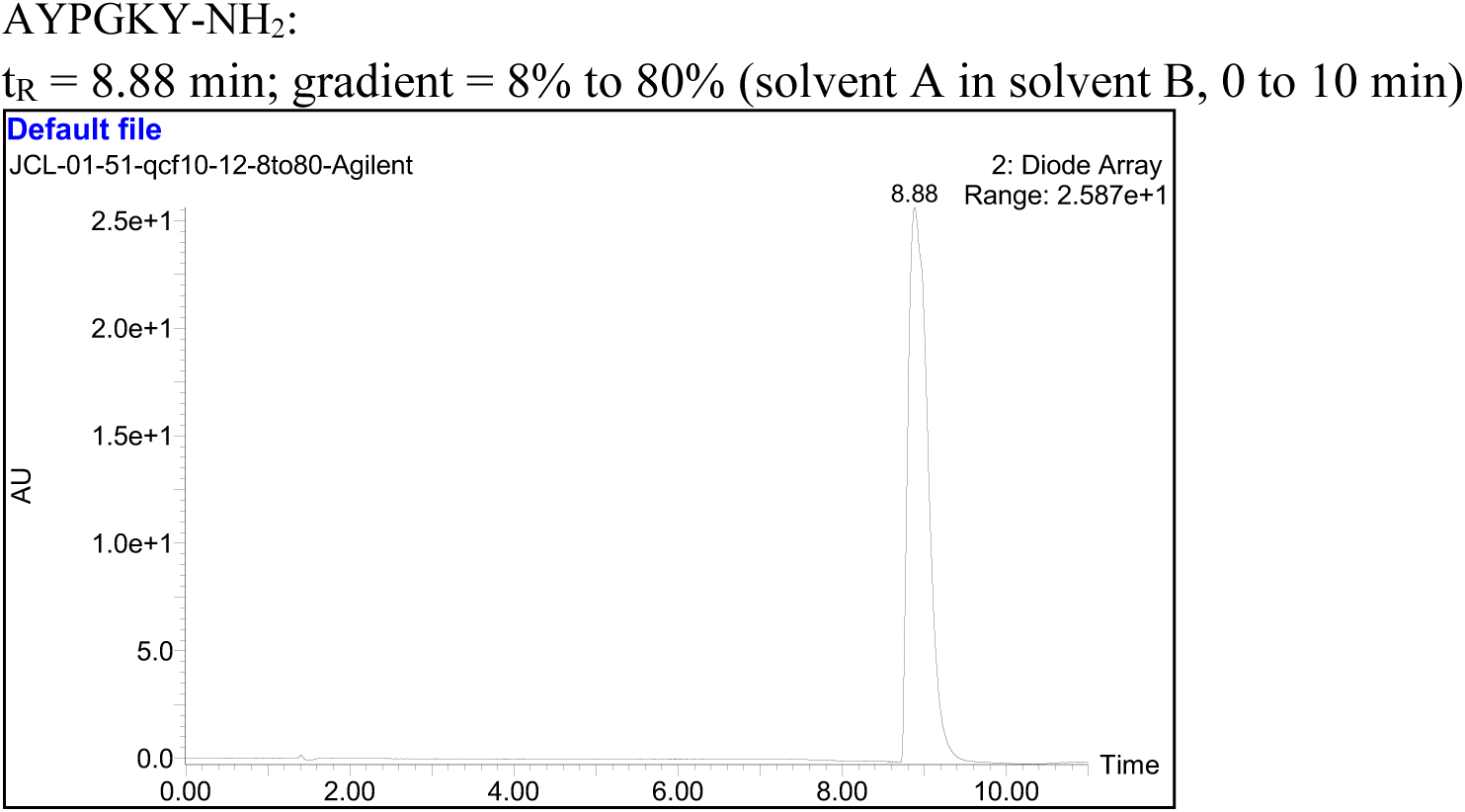
Analytical RP-HPLC UV detection chromatograms of synthesized peptides. Purity of all peptides are > 95% as determined by analytical RP-HPLC UV detection (see traces below).

**Figure S2.**
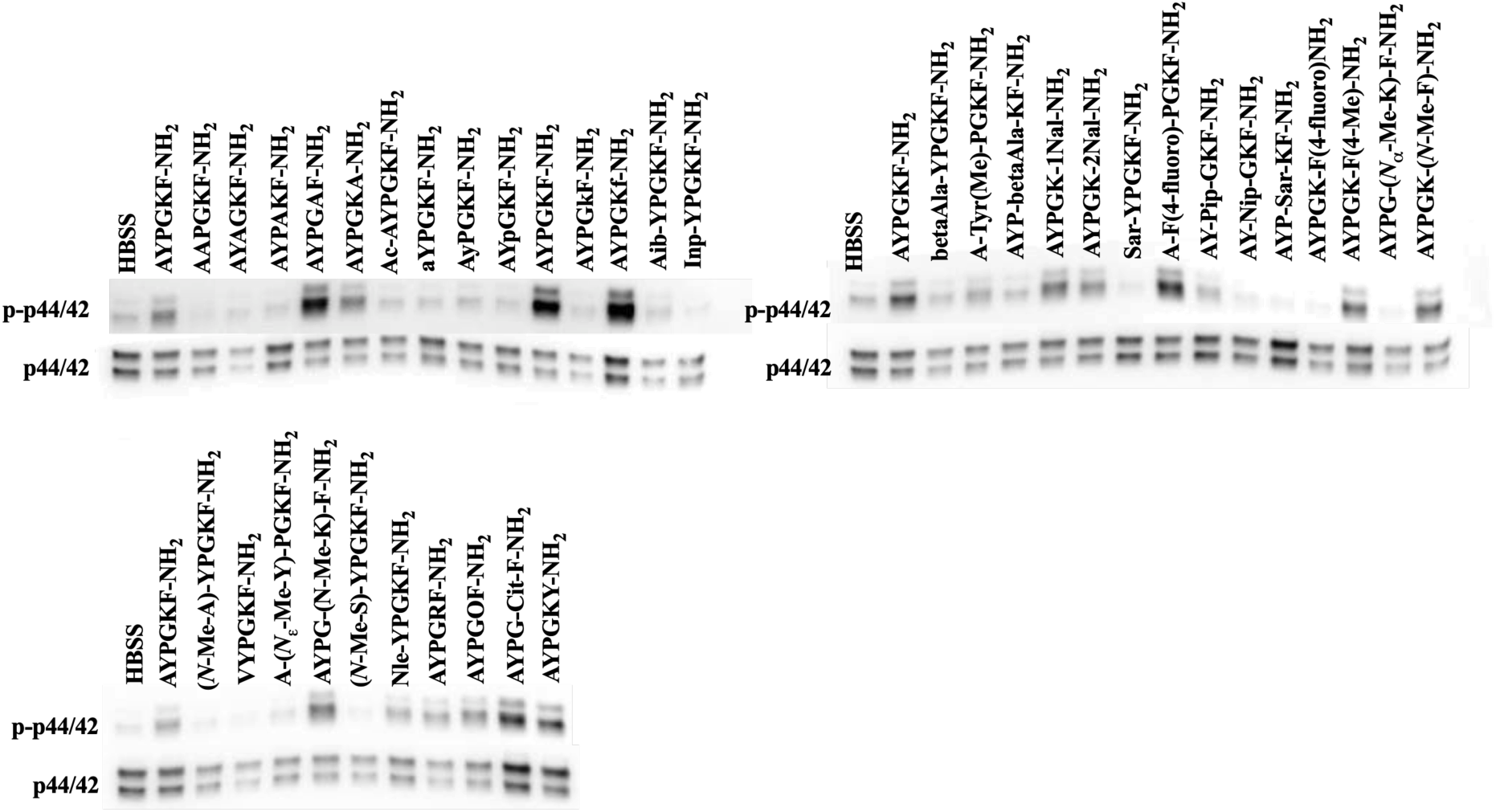
Representative blots of ERK phosphorylation in response to 100 µM of AYPGKF-NH_2_ and derivative peptides. Data are summarized in figure 10.

**Figure S3.**
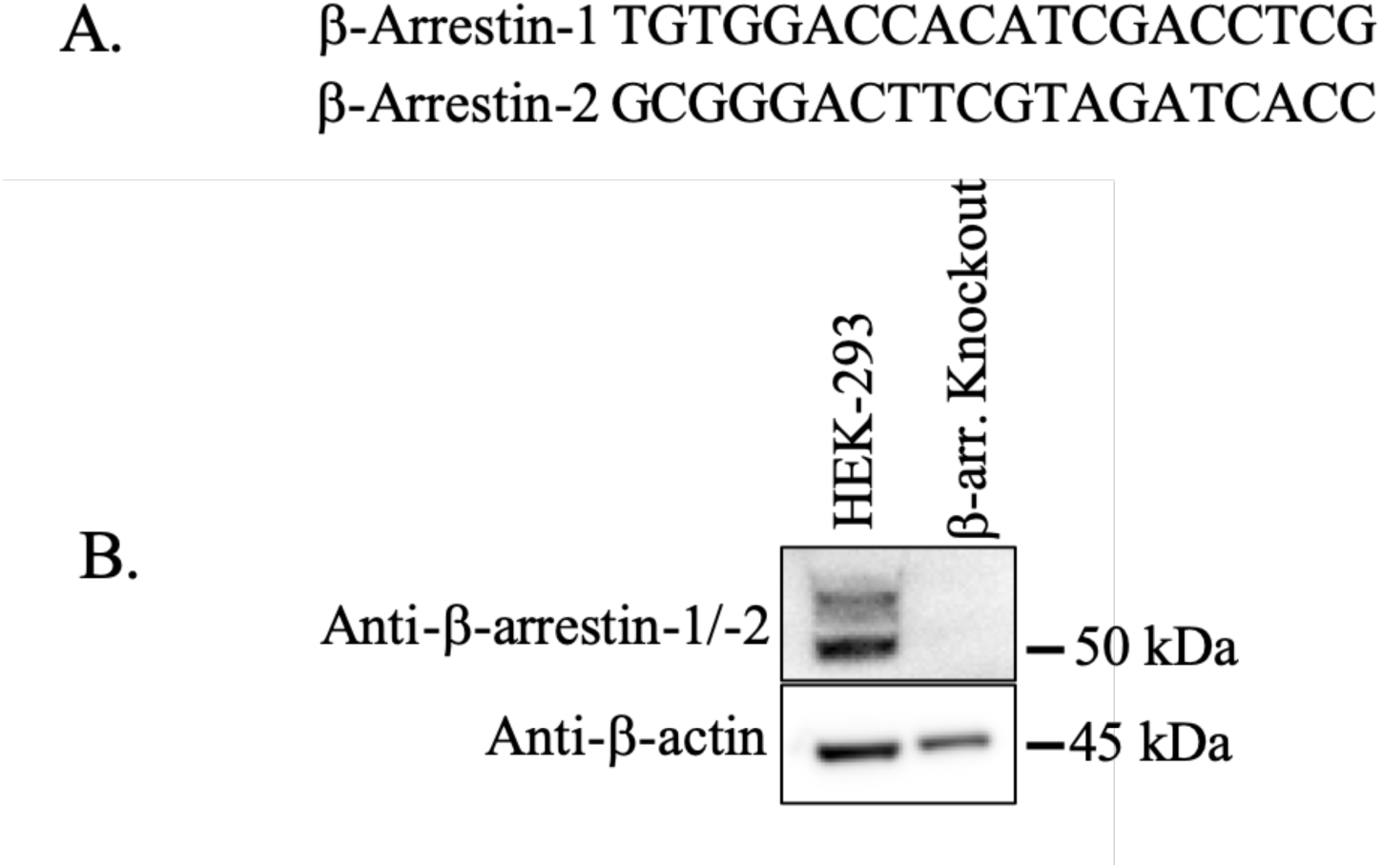
(A) Guide RNA sequences designed to target either β-arrestin-1 or -2. (B) Representative blots of β-arrestin-1/-2 protein from HEK-293 and β-arrestin-1/-2 double knockout HEK-293 (β-arr. Knockout) cells. We observe no β-arrestin-1/-2 protein in the β-arr. knockout HEK-293 cell line.

**Figure S4.**
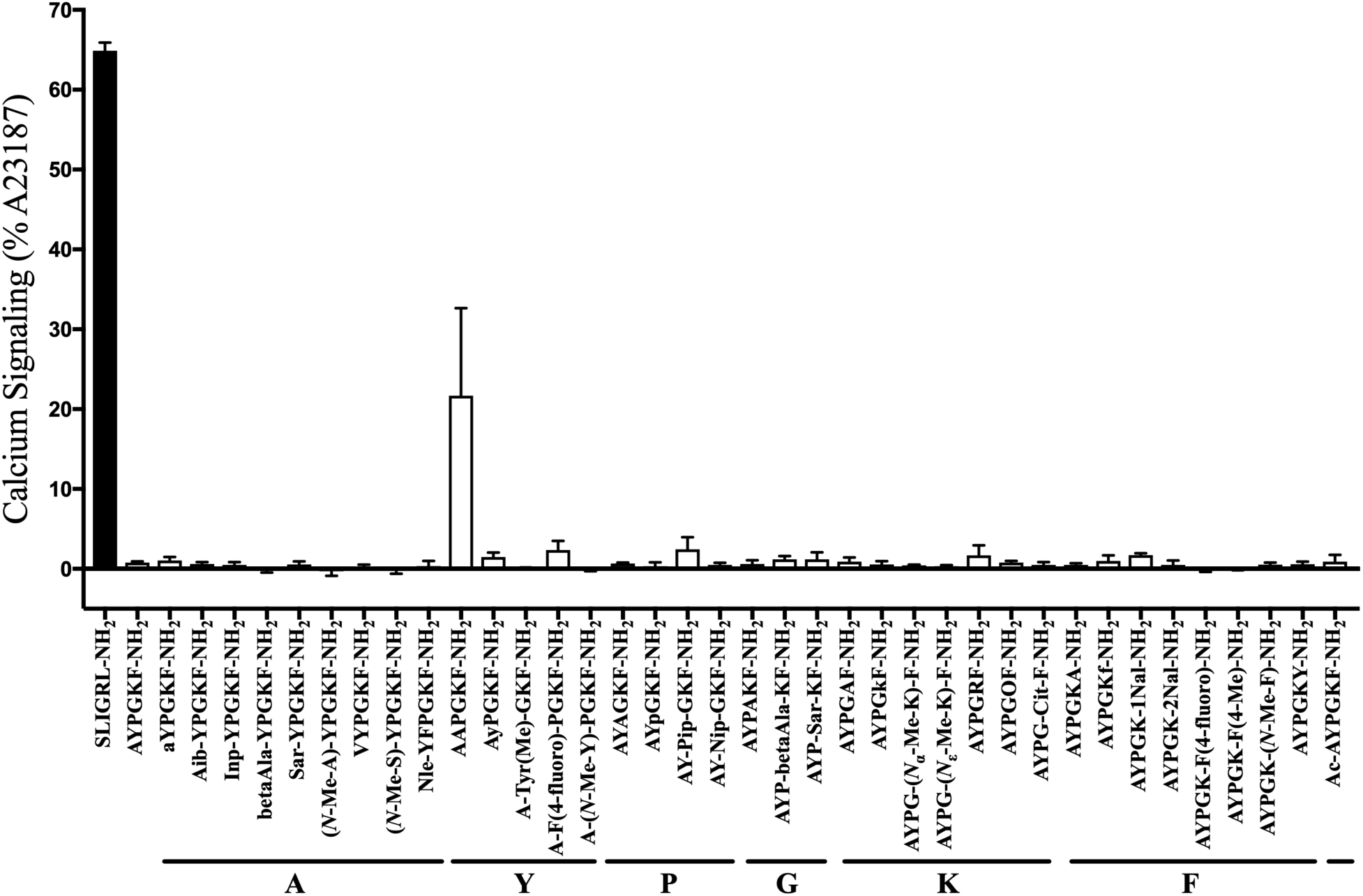
PAR4 peptide library is specific for PAR4 as shown by lack of calcium signalling in HEK-293 cells which do not endogenously express PAR4. Peptide AAPGKF-NH_2_ was found to have basal fluorescence even in the absence of cells. (*n = 3*)

**Figure S5.**
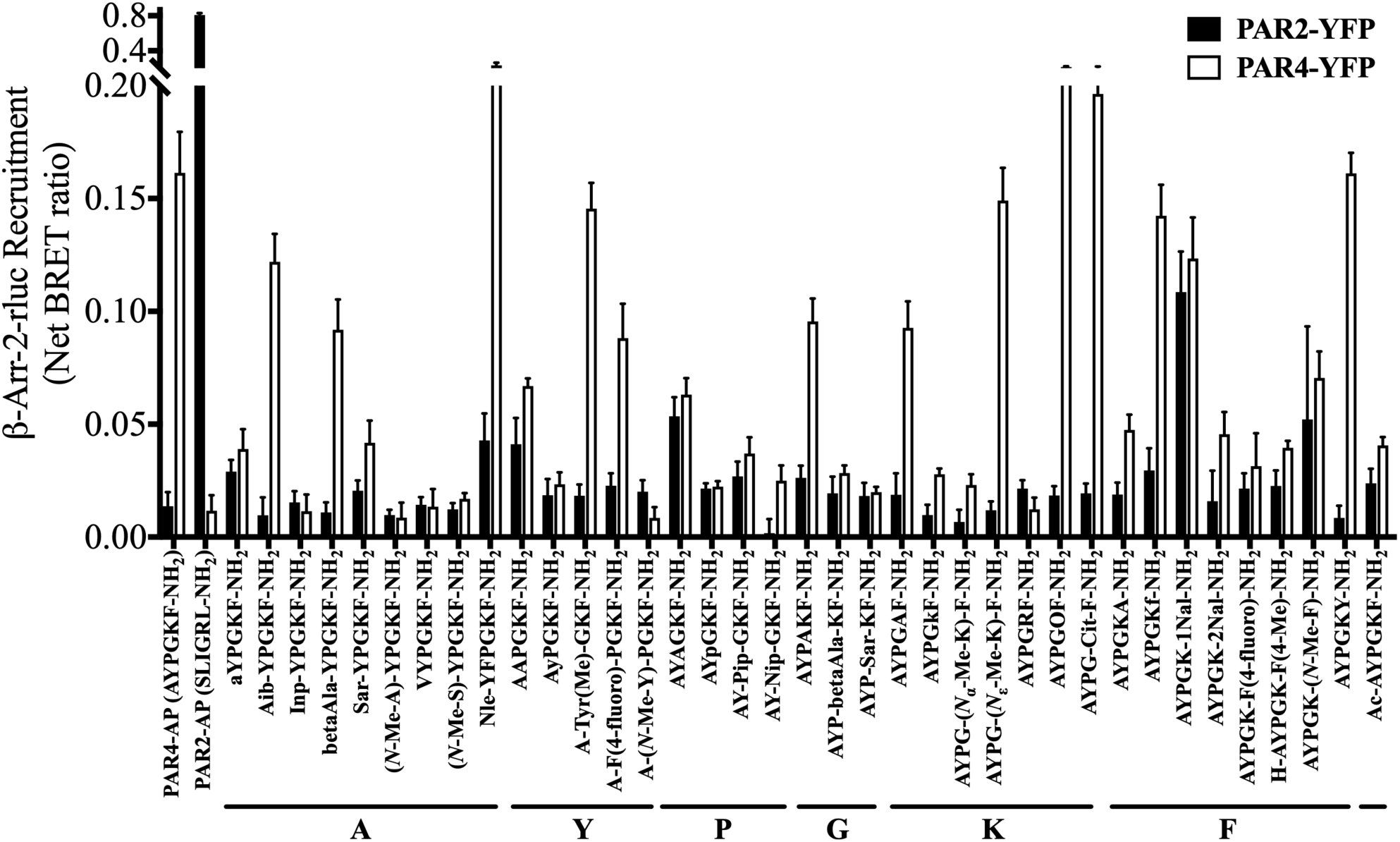
PAR4 peptide library is specific for PAR4 and does not cause recruitment of β-arrestin-2 to PAR2 (shown as black bars) at 100 µM, with the exception of AYPGK-1Nal-NH_2_ which shows some β-arrestin recruitment to PAR2. BRET ratios obtained with the parental peptides for both PAR4 (AYPGKF-NH_2_) and PAR2 (SLIGRL-NH_2_) are shown. (*n = 3*)

**Table S1:**
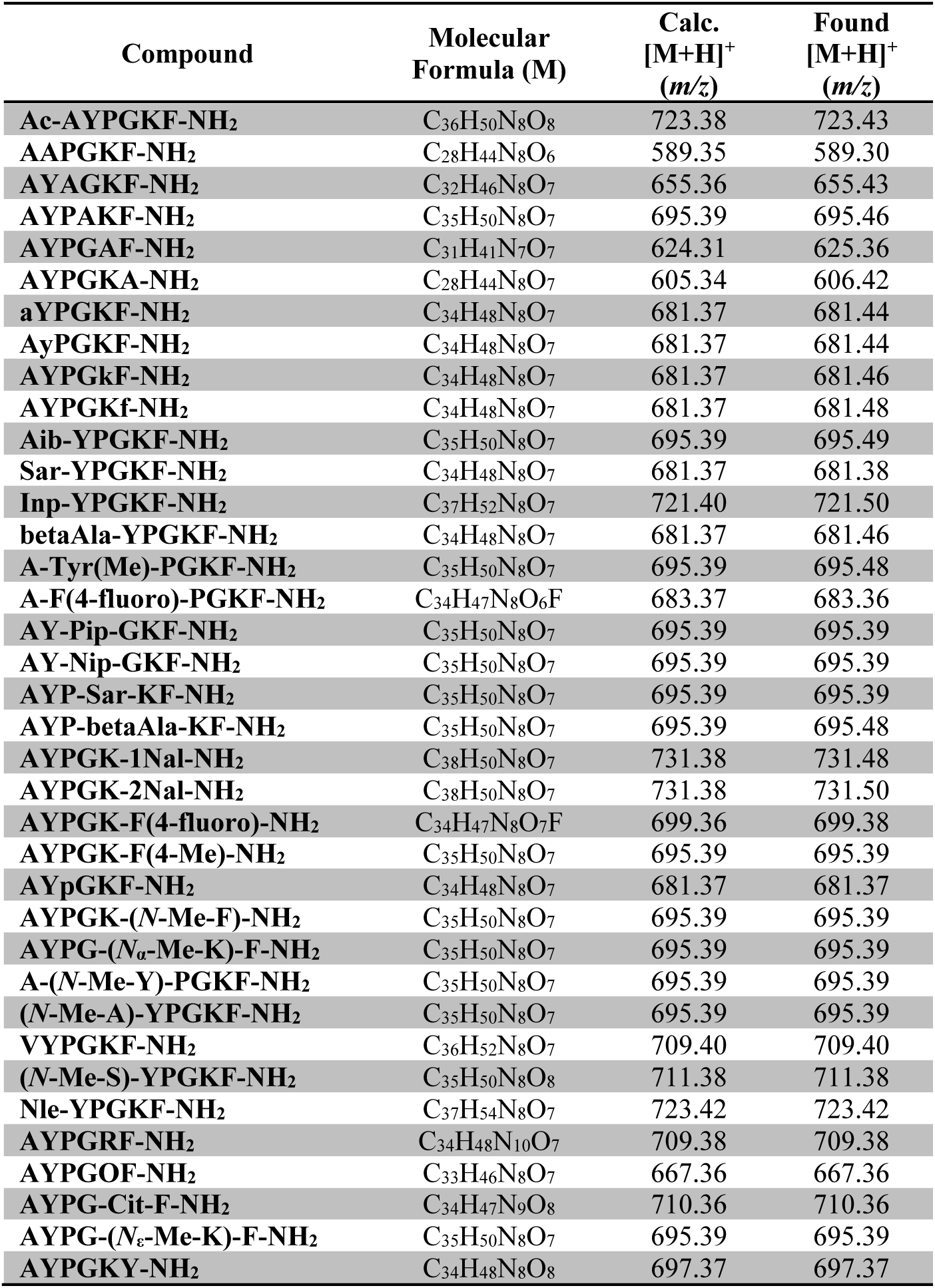
MS data of synthesized peptides.

